# CINner: modeling and simulation of chromosomal instability in cancer at single-cell resolution

**DOI:** 10.1101/2024.04.03.587939

**Authors:** Khanh N. Dinh, Ignacio Vázquez-García, Andrew Chan, Rhea Malhotra, Adam Weiner, Andrew W. McPherson, Simon Tavaré

## Abstract

Cancer development is characterized by chromosomal instability, manifesting in frequent occurrences of different genomic alteration mechanisms ranging in extent and impact. Mathematical modeling can help evaluate the role of each mutational process during tumor progression, however existing frameworks can only capture certain aspects of chromosomal instability (CIN). We present CINner, a mathematical framework for modeling genomic diversity and selection during tumor evolution. The main advantage of CINner is its flexibility to incorporate many genomic events that directly impact cellular fitness, from driver gene mutations to copy number alterations (CNAs), including focal amplifications and deletions, missegregations and whole-genome duplication (WGD). We apply CINner to find chromosome-arm selection parameters that drive tumorigenesis in the absence of WGD in chromosomally stable cancer types. We found that the selection parameters predict WGD prevalence among different chromosomally unstable tumors, hinting that the selective advantage of WGD cells hinges on their tolerance for aneuploidy and escape from nullisomy. Direct application of CINner to model the WGD proportion and fraction of genome altered (FGA) further uncovers the increase in CNA probabilities associated with WGD in each cancer type. CINner can also be utilized to study chromosomally stable cancer types, by applying a selection model based on driver gene mutations and focal amplifications or deletions. Finally, we used CINner to analyze the impact of CNA probabilities, chromosome selection parameters, tumor growth dynamics and population size on cancer fitness and heterogeneity. We expect that CINner will provide a powerful modeling tool for the oncology community to quantify the impact of newly uncovered genomic alteration mechanisms on shaping tumor progression and adaptation.

## INTRODUCTION

Chromosomal instability (CIN) is a hallmark of cancer, characterized by structural and numerical chromosomal alterations in tumor tissues over time. Key manifestations of chromosomal instability include chromosome missegregation, whole-genome doubling (WGD) and extrachromosomal DNA [1]. CIN generates genetic diversity and phenotypic variation among cancer cells, which can facilitate their adaptation to different environmental challenges, such as metastasis, drug resistance, and immune evasion [2]. On the other hand, CIN can also impair cell fitness by causing cellular stress, impaired DNA repair, and reduced proliferation. The role of CIN in cancer is therefore complex and context-dependent, and depends on the balance between its benefits and costs. To better understand how CIN affects cancer evolution and cell fitness, it is necessary to integrate experimental and computational approaches that can capture the temporal dynamics and consequences of CIN in tumor tissues.

We present CINner, a framework for modeling chromosomal instability during cancer evolution. CINner is designed to output data that are compatible with both bulk and single-cell DNA sequencing methods, enabling the analysis of tumor heterogeneity and clonal evolution at different levels of resolution. One of its advantages over existing algorithms is the ability to accommodate distinct copy number aberration (CNA) mechanisms that result from CIN and collectively transform a cell’s karyotype and fitness. CINner uses a number of numerical techniques to enhance the speed and efficiency of the simulations. It can generate tumors with sizes and karyotypes that match observations from DNA-sequencing data from real cancer samples. The framework allows for easy implementation of genomic events ranging in size from WGD to focal amplification/deletion and point mutations. The selection component of CINner is formulated as a function mapping a cell’s karyotype and single nucleotide variants (SNVs) to its fitness. At one extreme, cell fitness can be defined solely upon the aneuploidy pattern, which is appropriate for studying certain solid tumors with prevalent widespread CNAs [3–5]. At the other extreme, CINner can model cancers that are predominantly diploid and driven only by recurrent point mutations and focal indices targeting specific driver genes [6–8]. As most cancers can be characterized as driven mainly by recurrent mutations or CNAs, or a mixture of both [9], CINner is uniquely positioned to uncover evolutionary patterns in many tumors. Finally, cancer cells in CINner evolve according to a stochastic branching process model, constrained by the carrying capacity of the environment. The tumor growth pattern even in the same cancer type can vary between exponential and logistic with a decades-long steady-state level, with implications for genetic composition, disease progression and clonal extent [10]. The carrying capacity model therefore provides the flexibility to examine the effects of the tumor dynamics on its heterogeneity and fitness.

## RESULTS

### CINner models chromosomal instability during cancer evolution

In CINner, each cell is characterized by its copy number (CN) profile, or driver single nucleotide variant (SNV) profile, or both (**Fig. 1a**). As genomic regions are amplified or deleted as copy number aberrations (CNAs) occur, the SNVs residing in those regions are correspondingly multiplied or lost. CINner models cancer evolution as a branching process [11]. Cell lifespan is exponentially distributed with an input turn-over rate, similar to previous works [12,13]. At the end of its lifespan, the cell either divides or dies. This assumption is mathematically equivalent to other models such as [14], where cell division and death are simulated as two independent exponentially distributed processes [15]. The probability for a cell to divide depends on its fitness, determined by its CN and mutation profiles according to a selection model. The division probability is also calibrated so that the total cell population follows an established dynamic. After a cell division, daughter cells either have the same profiles as the mother cell, or harbor CNA or driver SNVs events resulting in new profiles.

**Figure 1.**
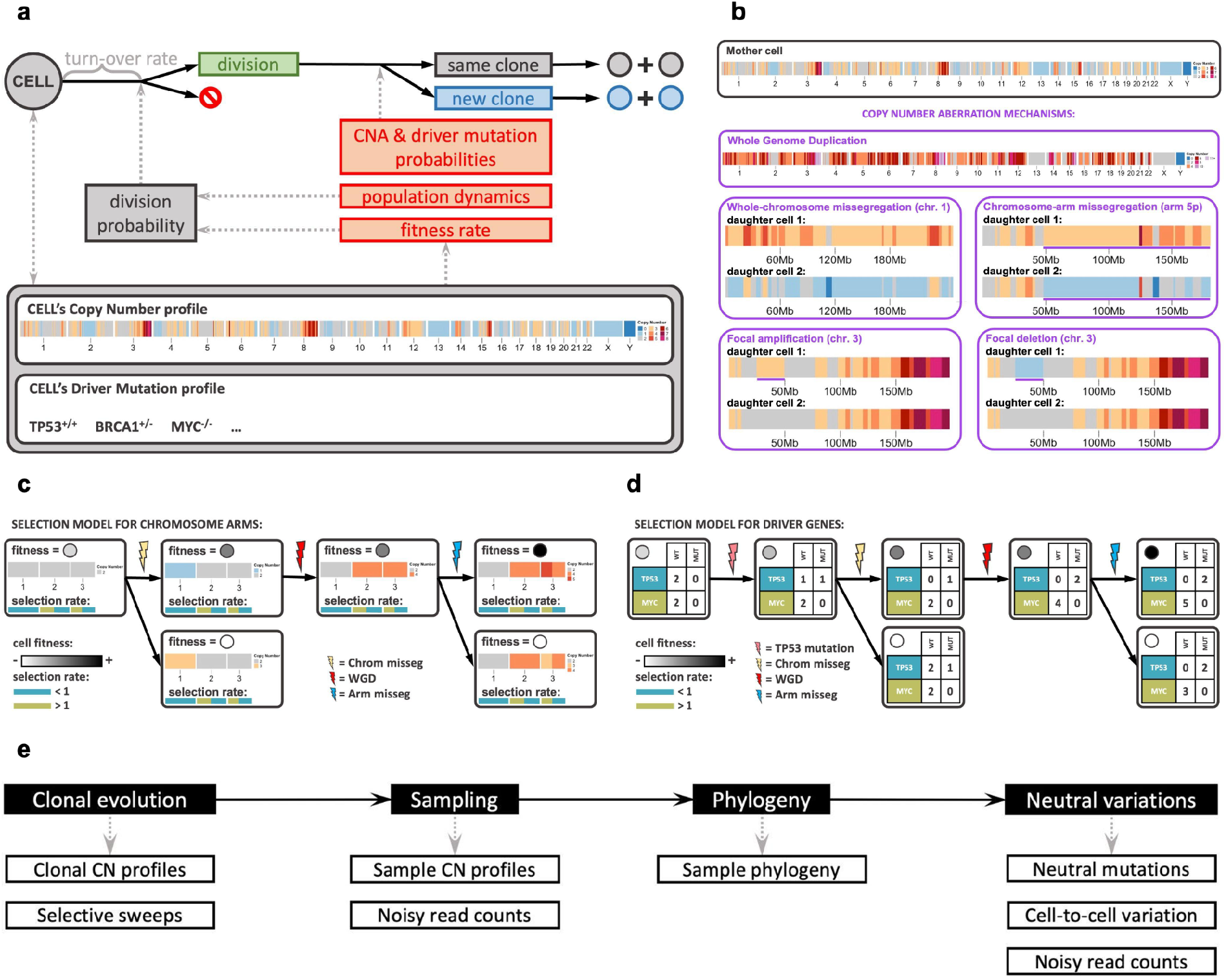
Overview of CINner’s mathematical model and simulation algorithm. **(a)** Each cell is characterized by a copy number profile and/or driver mutation profile, which define its fitness rate. The lifespan for all cells is exponentially distributed with the same turn-over rate. The probability of a cell to divide instead of dying takes into account its fitness rate and the total cell count, such that the total population follows known dynamics. If a cell divides, it can create new clones if a Copy Number Aberration (CNA) event occurs or a new driver mutation is acquired, otherwise the daughter cells belong to the same clone. **(b)** Multiple CNA mechanisms can occur during a cell division. Whole Genome Duplication results in one daughter cell with doubled genomic materials. Whole-chromosome missegregation misplaces one chromosome strand, chosen at random, in one daughter cell instead of the other (chr. 1 in the example). Chromosome-arm missegregation misplaces a random chromosome arm (5p in the example). Focal amplification and deletion choose a random region on a random chromosome arm, and either doubles or deletes the genomic materials there (25-50Mb region on chr. 3 in the example). Purple lines denote CN regions in daughter cells affected by CNAs. **(c-d)** Two selection models included in the model. Squares represent cells, profiles of which change according to CNAs and driver mutations. Circles in each cell represent its fitness (darker is fitter). **(c)** Selection model for chromosome arms. Arms dominated by Tumor Suppressor Genes (TSGs) have selection rates < 1, their losses are beneficial and gains are deleterious. The opposite holds true for arms with Oncogenes (OGs) and selection rates > 1. WGD does not change the arm balance and therefore the cell fitness rate remains constant. **(d)** Selection model for driver genes. Each driver gene has a selection rate for its wild-type (WT) and mutant (MUT) alleles. The balance of all driver gene allele counts and their selection rates defines a cell’s fitness rate. A cell is more advantageous if a TSG is either mutated or lost, or an OG is either mutated or gained. Here TP53 represents a TSG and MYC represents an OG. A third hybrid selection model is a combination of **(c)** and **(d)**. All selection models are further subject to viability checkpoints. If a cell violates thresholds on nullisomy extent, maximum bin CN, driver counts, etc. then its fitness rate is zero and the cell eventually dies. **(e)** Schematics of the simulation algorithm, divided into 4 main steps. Each step additionally outputs data requested by the user.

Previous mathematical models have mainly studied the evolution of SNVs during cancer development [16,17]. Some recent works have focused instead on analyzing the heterogeneity and convergence of tumor CN [18,19]. CINner is distinct from most cancer evolution models in its ability to incorporate both SNVs and CNAs during cancer evolution and study how they impact the selection landscape simultaneously. SimClone1000 [20] is another algorithm capable of generating synthetic tumor data with both genomic change classes. CINsim [21] is another method that allows modeling of CNAs in single cells and focuses on inferring rates of chromosome missegregation. However, unlike other methods, CINner can accommodate five distinct CNA mechanisms, each with distinct alteration patterns and varying impacts on cell fitness (**Fig. 1b**). Whole-genome duplication (WGD) results in one daughter cell with double the genomic material of the mother cell. Whole-chromosome missegregation misplaces a chromosome strand among the two daughter cells. In contrast, only a strand arm is misplaced in chromosome-arm missegregation. Finally, focal amplification and deletion target a random region in a strand arm, and either doubles the genomic material there or deletes it in a daughter cell.

Three selection models are included. The first model characterizes the selection of chromosome arms (**Fig. 1c**). If the selection parameter *s* of a chromosome arm is larger than 1, gaining the arm increases cell fitness and losing the arm is deleterious. The opposite holds if the arm selection parameter is less than 1. The magnitude of |1 − *s*| defines the selective (dis)advantage of such gains and losses. The selection parameter serves as an indicator for the balance of tumor suppressor genes (TSGs) and oncogenes (OGs), as arms with high OG counts are commonly amplified and arms with many TSGs frequently get lost in cancer [22]. The model for selection of driver mutations (**Fig. 1d**) seeks to portray the selection of TSGs and OGs directly. In this model, the selection parameters for the wild-type (WT) and mutant (MUT) alleles of a gene, are defined according to whether the gene functions as a TSG or an OG in that specific cancer type. The formulation dictates that a cell’s fitness increases if a TSG is mutated or lost, or an OG is mutated or gained. Thus, the model is based on the “one-hit” hypothesis [23], where each additional hit to a gene renders the cell more advantageous. The third model is a combination of these two models, describing cancer as driven both by small events targeting driver genes and large CNAs changing gene balance across the genome. As all three models are defined upon the gene balance in a cell, which is retained after WGD, a WGD cell has the same fitness as its parental cell. Furthermore, each selection model is subject to viability checkpoints, which eliminate cells that exceed defined thresholds on driver mutation count, bin-level CN, average ploidy, or extent of nullisomy. **Supplementary Note 1** describes the mathematical model in detail.

CINner is developed to efficiently simulate observed SNVs and CNAs in a tumor sample (**Fig. 1e**). To optimize for computing memory and runtime, the genome is divided into bins of a fixed size, and the allele-specific bin-level copy number profile of each cell is tracked throughout tumor progression. Each new mutation is assigned a genomic location, and gets multiplied or deleted if the site is affected by later CNAs. Two observations are utilized to increase the efficiency of CINner. First, cells with the same phylogenetic origin share the same CN and mutational profiles, therefore they evolve similarly throughout time. Second, the information relevant for downstream analysis is restricted to only the sampled cells. Therefore, it is not necessary to simulate single cells in the whole population individually, and instead we focus on clones, defined as groups of cells that have identical CN and mutational characteristics. The first step of CINner consists of simulating the evolution of clones in forward time. New clones are generated when CNAs or driver mutations occur, and the clone sizes change through time according to the branching process governing cell division and death. We use the tau-leaping algorithm [24] for efficiency, as the exact Gillespie algorithm [15] is time-consuming for cell populations of the typical size of tumors. In the second step, CINner samples cells from predefined time points. Next, it constructs the phylogeny for the sampled cells by using the “down-up-down” simulation technique [25]. The sampled cell phylogeny is generated as a coalescent (cf. [26]), informed by the recorded clone-specific cell division counts throughout time from step 1. This procedure is more efficient than directly simulating the branching process for the total cell population then extracting the phylogeny only for the sampled cells, as the sample size is typically of magnitudes smaller than the total cell population. Finally, cell-to-cell variations due to neutral CNAs and passenger mutations are simulated on top of the phylogeny tree and trickle down to the sample observations. CINner is explained in more detail in **Supplementary Note 2**.

### Selection parameters calibrated for chromosome arms predict gene imbalance and prevalence of whole-genome duplication

We develop a parameter estimation program for the chromosome arm selection model (**Fig. 1c**), which employs the Approximate Bayesian Computation random forest (ABC-rf) method [27] (**Supplementary Note 3**). We find that simultaneous parameter inference for both selection parameters and CNA probabilities in bulk DNA-sequencing data results in nonidentifiability issues. Previous works have observed around 1 – 9 × 10^−3^ missegregations per division in cancer cell lines [28–30]. However, in CINner this figure can be explained by either (i) high CNA probabilities coupled with selection parameters close to 1, or (ii) low CNA probabilities and selection parameters farther from 1. We will examine this in more detail in the next section.

In this section, we choose to study the selection parameters for each cancer type, and whether they indicate the tissue-specific selective pressure. Therefore, we fix the whole-chromosome missegregation probability at a comparatively low rate of 5 × 10^−5^ for all cancer types, so it is easier to analyze the inferred selection parameters. Given a CN data cohort, we find 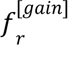 and 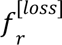, the frequencies of gain and loss for each chromosome arm *r*. The ABC method then finds the posterior probability distributions for the arm selection parameters and the chromosome-arm missegregation probability that explain the observed gain and loss frequencies. We obtain a point estimate for each parameter by using its maximum a posteriori probability (MAP) estimate, which is the mode of the parameter’s posterior distribution [31]. Finally, we generate CINner simulations with this estimated parameter set for direct comparison against the CNA data. The estimated chromosome-arm missegregations appear to be similar across different tumor types, possibly as a result of the nonidentifiability (**Supplementary Fig. 1-Supplementary Fig. 17**). Therefore, we focus the analysis on the inferred selection parameters.

We first employ the parameter fitting routine to study distinct cancer types with available data in PCAWG [32]. Samples with whole-genome duplication (WGD) have been shown to exhibit significantly different selection forces from non-WGD samples of the same cancer type, especially with respect to chromosome arm gains and losses [33]. Therefore, we limit the data to only non-WGD samples in each PCAWG data type for parameter calibration. We apply the parameter inference to 17 cancer types with *n* ≥ 10 non-WGD samples. In the CINner framework, gains of a chromosome arm *r* with selection parameter *s*_r_ ≫ 1 are selective, therefore DNA samples exhibit high 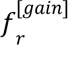. Conversely, chromosome arms with *s_r_* ≪ 1 exhibit high 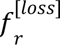. On the other hand, arms with low 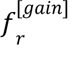 and 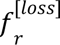 and do not frequently get altered copy number, so we assume that these arms are neutral. Therefore, we limit the inference only to chromosome arms *r* with 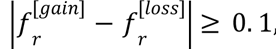 to mitigate the effects of neutral evolutionary noise.

**Fig. 2a** presents the fitting results for glioblastoma samples (CNS-GBM). This cancer type has relatively few frequent CNAs, except for the combination of chromosome 7 gain and chromosome 10 loss [34], and gains of chromosomes 19 and 20 at lower frequencies [35]. Compared to CNS-GBM, ovarian adenocarcinoma (Ovary-AdenoCA) contains extensive CNAs shaped by multiple mutational processes [36], especially genomic loss-of-function events in *BRCA1* and *BRCA2* genes [37] (**Fig. 2b**). Finally, breast adenocarcinoma (Breast-AdenoCA) is also associated with high aneuploidy [3] (**Fig. 2c**). For each cancer type, the gain/loss frequencies produced from the simulator with fitted selection parameters closely resemble the genomic CNA landscape from PCAWG. Additionally, the selection parameters for individual chromosome arms correlate strongly with their amplification or deletion proportions (**Fig. 2d-f**). Similarly, the inferred chromosome arm selection parameters for the other 14 cancer types lead to similar CNA landscapes to those from PCAWG (**Supplementary Fig. 1-Supplementary Fig. 17**). Overall, this demonstrates the model’s ability to uncover specific selection forces characteristic of particular cancers, regardless of the extent of aneuploidy or bias toward genomic gains or losses.

**Figure 2.**
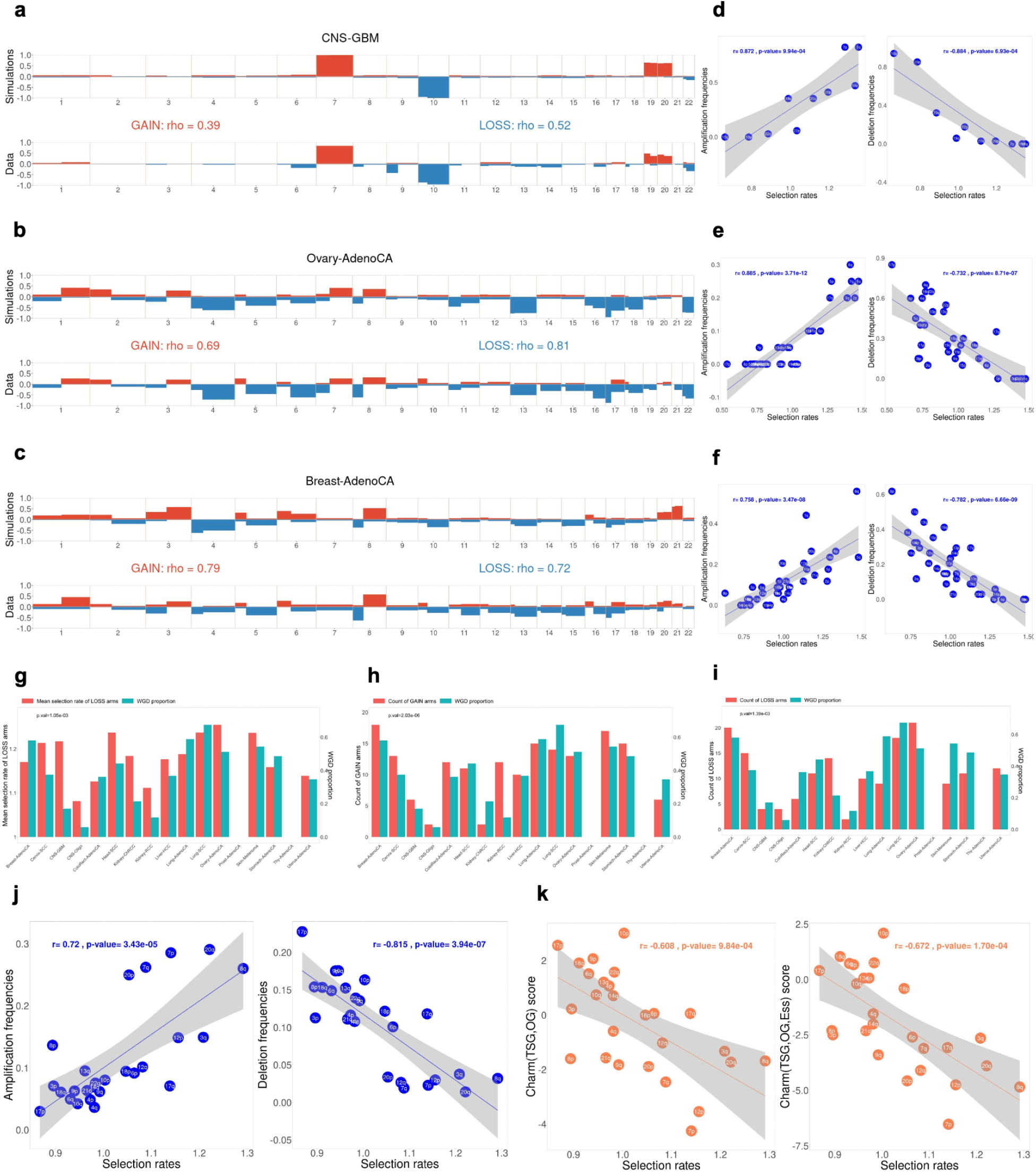
Results from fitting the chromosome arm selection model to CN data from PCAWG and TCGA. **(a-c)** Comparison between gain/loss frequencies from the fitted model (top) and non-WGD samples in PCAWG (bottom) for the cancer types diffuse glioma in central nervous system **(a;** CNS-GBM), ovary adenocarcinoma **(b;** Ovary-AdenoCA) and breast adenocarcinoma **(c;** Breast-AdenoCA). Spearman’s correlation coefficient rho between frequencies of gains (or losses) among all bins in PCAWG and simulations. **(d-f)** Correlation between inferred chromosome arm selection rates and amplification/deletion frequencies for individual chromosome arms in CNS-GBM **(d)**, Ovary-AdenoCA **(e)** and Breast-AdenoCA **(f)**. In CINner, gains of arms with selection rates > 1 are advantageous, similarly as losses of arms with selection rates < 1. Linear regressions and p-values from Pearson correlation. **(g-i)** Correlation between WGD proportion and mean selection rates of LOSS arms **(g)**, and counts of GAIN arms **(h)** and LOSS arms **(i)** in each PCAWG cancer type. p-values from Spearman correlation between WGD proportion and each variable. **(j, k)** Fitted selection rates versus TCGA pan-cancer chromosome arm gain/loss frequencies **(j)** and gene balance scores **(k)** (Davoli et al. 2013). The score Charm(TSG-OG) considers the gene imbalance between TSGs and OGs, and Charm(TSG-OG-Ess) additionally examines Essential genes. Linear regressions and p-values from Pearson correlation.

We then examine whether the estimated parameters are indicative of cancer properties, specifically the selection for whole-genome duplication (WGD). It occurs in about 30% of tumors and is associated with a poor prognosis, suggesting that it plays a crucial role in cancer development [38]. WGD is also linked with extensive and profound changes in the selective landscape, including heightened chromosomal instability [39,40], increased preference for losses over gains [41], and changes in co-occurrence and mutual exclusivity in aneuploidy patterns [33]. Because of WGD’s typical occurrence in initial stages of tumorigenesis and the many genomic changes it causes up to diagnosis [42], it is difficult to infer from DNA-sequencing data the causes for selection of WGD in early cancer development. We investigate whether the chromosome arm selection parameters inferred from CINner can predict tissue-specific WGD prevalence in PCAWG. We also explore which features correlate strongly with WGD proportion, which would imply contribution to increased fitness for WGD cells over the non-WGD population. For a given cancer type, we classify chromosome arms *r* with 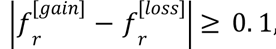 with inferred selection parameter *s*_*r*_> 1 as GAIN arms, and those with *s*_*r*_< 1 as LOSS arms. Each cancer type is then characterized by the counts of GAIN and LOSS arms, together with their respective mean selection parameters. It should be noted that in CINner, *s* > 1 indicates selection for gains and 0 < *s* < 1 implies losses are selected. Therefore, to normalize the selection parameters in the means, we use 1/*s* for LOSS arms and keep the inferred *s* for the GAIN arms.

One hypothesis for the prevalence of WGD in cancer is that WGD provides redundant genes to buffer the deleterious effects of nullisomy [40]. The risk of nullisomy increases if the CNA probabilities are high or if there exist LOSS arms with selection parameters *s* ≪ 1. Because our missegregation probabilities are similar across tissue types due to nonidentifiability, the hypothesis predicts that WGD is more frequently observed in cancers with highly selective LOSS arms. The correlation between WGD prevalence and mean selection parameters in LOSS arms inferred from CINner across cancer groups confirms this (**Fig. 2g**). Our results therefore are in agreement with the assumption that WGD helps cancer cells mitigate the risk of nullisomy from repeated losses in specific genomic regions [41,43]. On the other hand, the counts of GAIN and LOSS arms indicate the proportion of the genome that is under selection for CNAs. The correlation between WGD proportion and the counts of either GAIN or LOSS arms (**Fig. 2h, i**) is compatible with evidence that WGD is associated with chromosomal instability in cancer [40]. In conclusion, we have shown that the selective landscapes uncovered by CINner can predict tissue-specific WGD prevalence, indicating that the inferred selection parameters are biologically meaningful. Moreover, the cancer types with either (i) many GAIN and LOSS chromosome arms, or (ii) some LOSS arms with high selection parameters, are more likely to harbor WGD, indicating that selection for WGD in cancer development is driven by its role in helping tumors avoid nullisomy and tolerate aneuploidy.

Finally, we investigate if our classification of chromosome arms as GAIN or LOSS, and their selection parameters calibrated by the fitting routine, can reveal the genetic imbalances within the arms. We calibrate the model on frequencies of chromosome arm gains and losses from the pan-cancer data in TCGA [22] (**Supplementary Fig. 18**). Similar to the fitting results for PCAWG cancer types, there is a strong correlation between estimated arm selection parameters and the frequencies of either amplification or deletion (**Fig. 2j**). We then compare the fitted selection parameters to chromosome arm scores in [22]. For a given chromosome arm, the score Charm(TSG,OG) accounts for the count and potency of tumor suppressor genes (TSGs) and oncogenes (OGs). The score is higher for arms with higher count or increased potency of TSGs, and lower for arms more abundant with OGs. The second score Charm(TSG,OG,Ess) additionally considers essential genes (Ess), in the same manner as OGs. The selection parameters derived from our model correlate well with both scores (**Fig. 2k**), and the negative correlation reflects the selection model (**Fig. 1c**). Chromosome arms with selection parameters *s* >> 1 are under intense selective pressure to get amplified, indicating that they harbor many important OGs, hence low Charm(TSG,OG) or Charm(TSG,OG,Ess), and the opposite holds for arms with *s* << 1. The correlation is stronger for Charm(TSG,OG,Ess) than Charm(TSG,OG), possibly signaling the relevance of essential genes in shaping the selective landscape during cancer evolution. Overall, we have shown that the chromosome arm selection parameters uncovered in our model are biologically significant, as they reflect the bias in distribution and potency toward either tumor suppressor genes or oncogenes and essential genes. Therefore, the model can play a role in estimating the driver gene landscape in specific cancer types. Driver genes are largely identified through their mutation frequencies, therefore the cohort size limits the sensitivity to which rare driver genes can be detected [44]. However, the selection parameters fitted in our model are an estimate of the combined effects of genes located on the same chromosome arm, including both commonly altered genes and those with minor contributions to tumor growth. Importantly, the selection parameter fitting routine only requires a small cohort of cancer samples.

### Impact of selection, copy number aberration mechanisms and growth dynamics in the chromosome arm selection model

CINner provides a framework to study different families of models and analyze the impact of model parameters on observable statistics of individual cancer samples. We have shown that specific cancer types exhibit a wide discrepancy in chromosome arm driver count, the potency of these arms, and the distribution of GAIN and LOSS arms among them. We now examine the signals in the tumor samples resulting from this discrepancy. In this analysis, the selection parameters calibrated for the TCGA pan-cancer dataset (**Fig. 2i-j**, **Supplementary Fig. 18**) are denoted as scale x1. We then study the sample statistics when GAIN or LOSS chromosome arms increase their selection parameters (**Fig. 3a-d**). As the selection parameters of GAIN arms increase, chromosome amplifications are more advantageous. Cancer cells that have amplified GAIN arms are more selective and therefore more likely to expand and get fixed in the population, otherwise they become obliterated by other subclones. The increased preference for gains over losses hence leads to higher average ploidy in the sample (**Fig. 3b**). Although the samples contain more clonal gains, the counts of subclonal gains and losses decrease, because of shorter elapsed time from Most Recent Common Ancestor (MRCA) to when the sample is taken (**Fig. 3c**). Interestingly, the count of clonal losses also increases slightly, as deletions behave as hitchhikers to amplification drivers. Conversely, as LOSS chromosome arms are more selective, the clonal loss count increases significantly and the clonal gain count increases moderately, while the subclonal gain and loss counts decrease, resulting in lower average sample ploidy (**Fig. 3b, d**). In both cases, the heightened competition means that subclones either expand quickly or become extinct, therefore the tumor sample exhibits fewer subclones, lower Shannon diversity index, and more recent Most Recent Common Ancestor (MRCA) (**Fig. 3a**, **Supplementary Fig. 19**).

**Figure 3.**
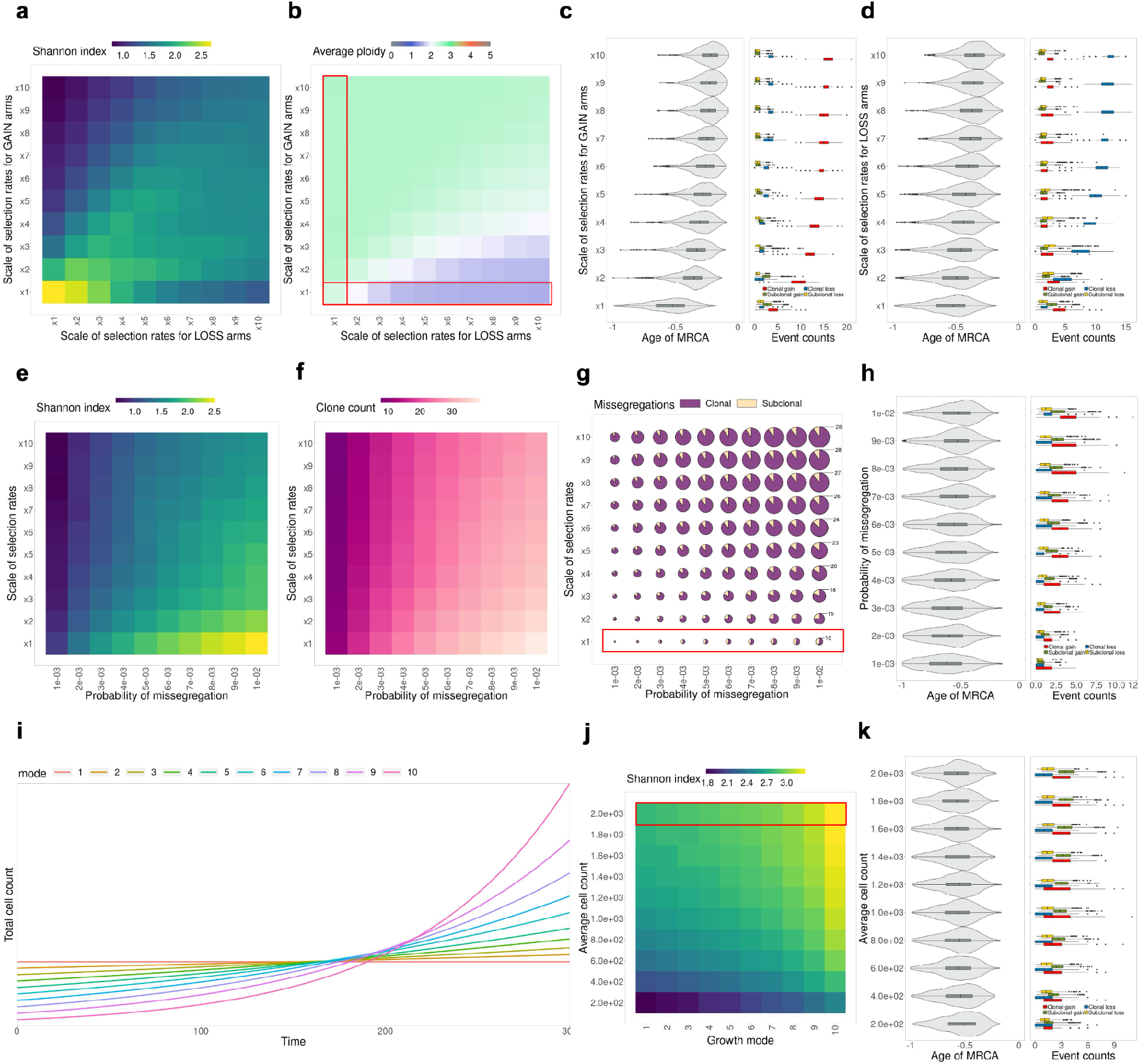
Analysis of parameters in the chromosome arm selection model. **(a-b)** Effects of selection rates for GAIN and LOSS chromosome arms on average Shannon diversity index **(a)** and average ploidy in each sample **(b)**. **(c-d)** MRCA age and average missegregation counts, grouped based on clonality (clonal/subclonal) and type (gain/loss), as selection rates for GAIN arms **(c)** or LOSS arms **(d)** increase (variables correspond to highlighted segments in **(b)**). MRCA age is computed as fraction over time back to when simulation starts. MRCA age = −1 if the sampled cell phylogeny tree starts branching at the beginning of the simulation. MRCA age → 0 as the MRCA is closer to sampling time. **(e-g)** Effects of probability of missegregation and chromosome-arm selection rates on average Shannon diversity index **(e)**, clone count **(f)** and average count of clonal and subclonal missegregations **(g)** (size of circles indicates the total missegregation counts). **(h)** MRCA age and missegregation counts as probability of missegregation increases (variables correspond to highlighted segment in **(g)**). **(i)** Different patterns of growth mode, ranging from constant (mode 1) to exponential with high growth rate (mode 10). **(j)** Effect of growth mode and average cell count on average Shannon diversity index. **(k)** MRCA age and missegregation counts as average cell count increases (variables correspond to highlighted segments in **(j)**). 1,000 simulations are created for every parameter combination.

Another region of interest is the effects associated with variable CNA probabilities on sample statistics. In particular, we study how different probabilities of missegregation impact cells’ fitness and tumor clonality (**Fig. 3e-h, Supplementary Fig. 20**). Although increasing selection parameters lead to heightened competition and therefore fewer subclones, a higher missegregation probability increases subclonal diversity and results in a larger Shannon diversity index (**Fig. 3e, f**). Because of the enhanced diversity, subclones share more clonal gains and losses, and they also harbor more subclonal missegregations (**Fig. 3h**). Because there is no change in the selection landscape, the MRCA age does not change appreciably. As a consequence, the ratio of clonal to subclonal gain and loss counts remains constant as probability of missegregation varies (**Fig. 3g**). This contrasts with increasing selection parameters, which likewise increases total missegregation count in the sample but with a higher bias toward clonal events. An important aspect to consider is that the total CNA count is the primary measure of CIN that can be obtained from bulk DNA-sequencing samples. However, as demonstrated in this study for missegregations, these counts can be explained by a spectrum of parameters in CINner, ranging from high CNA probabilities with low selective pressure (bottom right corner in **Fig. 3g**) to high selection parameters coupled with low CNA occurrences (top left corner). As discussed in the previous section, it is therefore challenging to estimate both CNA probabilities and selection parameters with bulk DNA data.

Finally, we investigate how different growth patterns impact the cancer sample statistics, taking advantage of the model’s ability to incorporate expected total cell count dynamics as input (**Fig. 3i-l**). The cell turn-over rate is unchanged under different tumor dynamics, hence the distribution of cell lifespan is constant. The sample size is also fixed at 1,000 cells for each parameter set. Despite this, as the population increases in size, the sampled cells both share more clonal missegregations and accrue more subclonal events (**Fig. 3k**), leading to higher Shannon diversity index (**Fig. 3j**). We also study ten different tumor growth models, ranging from constant to exponential with increasing growth rates (**Fig. 3i**). In tumors growing at a low rate, the competition for carrying capacity is more intense. In contrast, higher exponential growth rate represents faster growing tumors, in which cells do not have to compete as much for space. As a result, even subclones with low fitness can expand, leading to higher clone count and Shannon diversity index (**Fig. 3j, Supplementary Fig. 21**).

In conclusion, the different components of the model, ranging from CNA mechanisms to tumor dynamics to selection model, have distinct effects on common cancer sample statistics. These signals are important to analyze, as they directly affect the process of model calibration. When using CINner to estimate parameters for specific datasets, it is crucial to find values for model constants that conform to the corresponding tissue type and tumor growth.

### The role of whole-genome duplication in promoting chromosomal instability

In previous sections, we estimated cancer type-specific selection parameters and missegregation probabilities in diploid cell populations, in the absence of whole-genome duplication (WGD) (**Fig. 2**). WGD is a common and early event in many cancers [6], and is associated with an altered selection landscape [33] and heightened chromosomal instability (CIN) [45,46]. As has been observed in our parameter study, increasing CNA probabilities leads to higher heterogeneity, and the increased clonal competition results in greater tumor fitness (**Fig. 3e-h**). In this section, we utilize CINner to measure the CIN level associated with WGD in a cancer-specific context.

The WGD proportion varies substantially among different cancers (**Fig. 2g-h**). Moreover, the fraction of genome altered (FGA) among WGD and non-WGD samples in PCAWG further varies between cancer types (**Fig. 4a**). For instance, the genomes of WGD squamous cell lung carcinoma (Lung-SCC) are significantly altered, but not at a substantially higher level than non-WGD tumors. In contrast, kidney renal cell carcinoma (Kidney-RCC) has few genomic alterations on a non-WGD background, however the WGD tumor genomes exhibit a high CIN level. We already captured the genome alteration landscape in non-WGD cancers (**Fig. 2**). Therefore, in order to study the WGD-associated CIN, we characterize each PCAWG cancer type by two statistics: WGD proportion, and WGD FGA difference (defined as the mean difference in FGA between WGD and non-WGD tumors).

**Figure 4.**
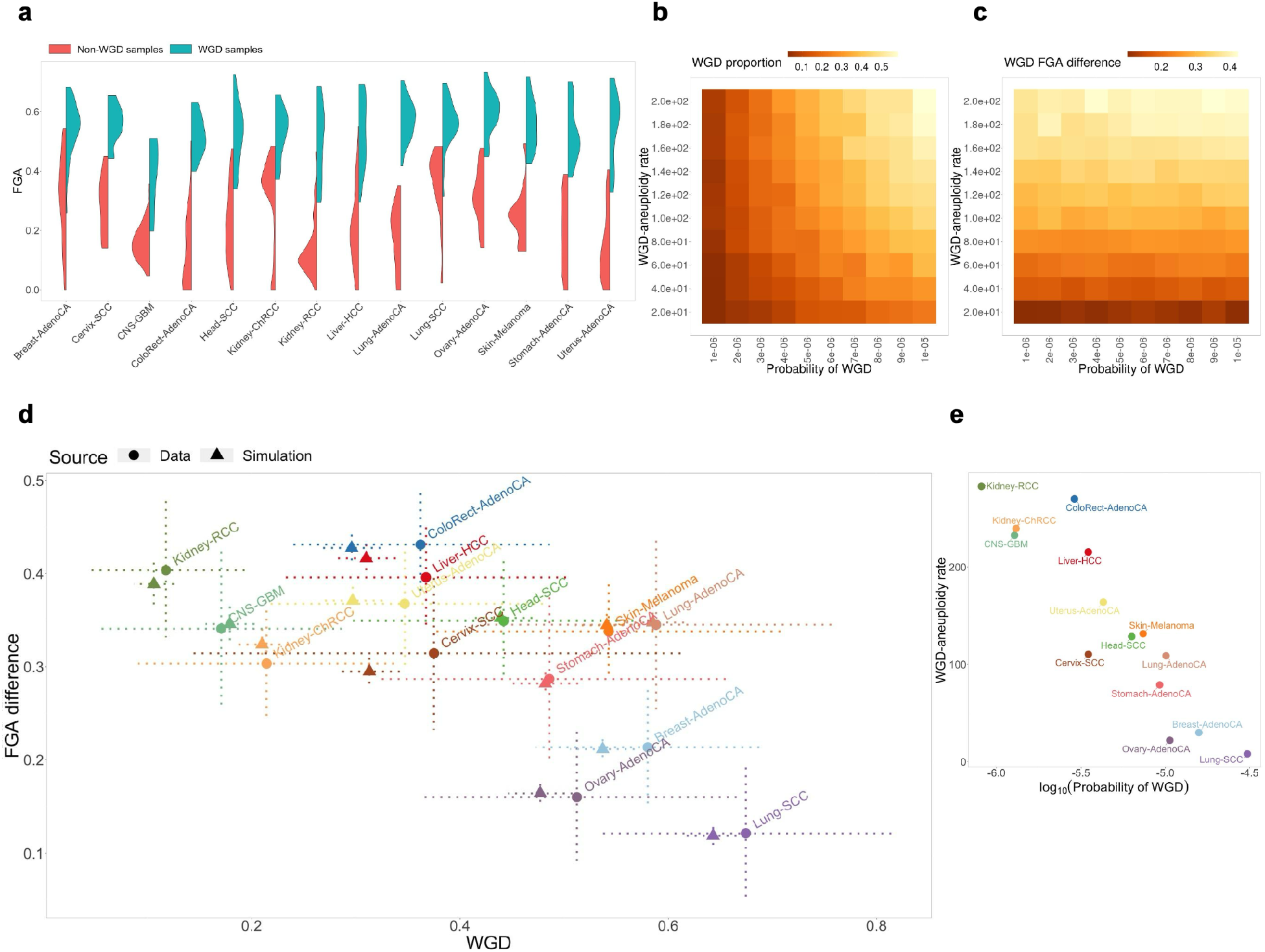
Results from fitting WGD parameters to CN data from PCAWG. **(a)** Distribution of Fraction of Genome Altered (FGA) in PCAWG by cancer type and WGD status. **(b-c)** Impact of varying WGD probability and WGD-aneuploidy rate on WGD proportion **(b)** and WGD FGA difference **(c)** in simulated samples. WGD FGA difference is defined as mean(FGA|WGD) - mean(FGA|non-WGD). 1,000 simulations are created for every parameter combination. **(d)** Comparison between WGD proportion and WGD FGA difference from the fitted model (triangle) and PCAWG (circle). Dotted bars represent ranges of each statistic from 10,000 bootstrap samples. **(e)** Comparison between inferred WGD probability (in logscale) and WGD-aneuploidy rate for each cancer type.

We define two parameters in CINner: the probability of WGD in each cell division, and WGD-aneuploidy rate, defined as the ratio of missegregation probabilities between WGD cells and non-WGD cells. We then study the changes in WGD statistics as these parameters vary, using the selection parameters fitted for the pan-cancer TCGA model (**Fig. 2i, j, Supplementary Fig. 18**). As can be expected, increasing WGD probability leads to higher WGD proportion within the simulated samples (**Fig. 4b**), however, the FGA difference between WGD and non-WGD samples is unchanged (**Fig. 4c**). On the other hand, increasing WGD-aneuploidy rate raises the FGA difference, as WGD-cells accumulate missegregations at a higher rate. The higher heterogeneity within WGD cells also leads to emergence of karyotypes with higher fitness compared to non-WGD cells, ultimately resulting in higher WGD proportion (**Fig. 4b**), similar to our observations from varying missegregation probabilities (**Fig. 3e-h**).

We now infer the WGD probability and WGD-aneuploidy rate for distinct cancer types from the WGD proportion and FGA difference in each PCAWG cohort. For each cancer type, we assume the chromosome-arm selection parameters and missegregation probabilities inferred previously (**Fig. 2**), then infer the WGD probability and WGD-aneuploidy rate with ABC-rf. To avoid overfitting, we limit the inference to 14 cancer types with at least 10 non-WGD samples and WGD proportion > 10% in PCAWG. The posterior probabilities are largely unimodal (**Supplementary Fig. 22, Supplementary Fig. 23**), indicating low uncertainty in the ABC inference. We simulated the WGD proportion and FGA difference using the modes for each inferred parameter, and the statistics are consistent with each PCAWG cohort (**Fig. 4d**).

The comparison of inferred parameters between different cancer types reveals that the WGD probability per cell division and WGD-aneuploidy rate are negatively correlated (**Fig. 4e**). One possible explanation is that there is a limit to the level of aneuploidy that can be tolerated in cancer cells, even in the presence of WGD. In cancer types with high WGD probability, there is a larger time span from WGD to diagnosis, as the event would frequently occur early in tumorigenesis. This results in increased aneuploidy, but also a large number of extreme karyotypes that are unviable. Therefore, the observed WGD samples exhibit much lower FGA as compared to expectations from CINner. Indeed, the FGA in WGD samples are more uniform across cancer types compared to the non-WGD samples (**Fig. 4a**). Another explanation is that the increased FGA in WGD samples results from increased likelihood of multipolar divisions [47]. The resulting progeny exhibit highly aneuploid genomes, and most are nonviable due to nullisomy. However, it is possible that rare surviving cells are more selectively advantageous than diploid cells, and expand across the tumor. Under this assumption, the WGD-aneuploidy rate would be lower, as the WGD cells would already have markedly higher FGA after multipolar divisions.

We also compare the inferred probability of WGD against the average fraction of monosomy in diploid samples from PCAWG. Under the hypothesis that WGD helps cancer escape nullisomy, we would expect a higher WGD probability in cancer types that have higher monosomy fraction in diploid samples. The comparison is unclear (**Supplementary Fig. 23g, h**). One potential reason is that there are few samples for certain cancer types in PCAWG. The approach that we employ necessitates further subdividing the samples based on WGD status, which could render the statistics too prone to noise. However, the three cancer types with highest monosomy fraction, namely cervical squamous cell carcinoma (Cervix-SCC), breast and lung adenocarcinomas (Breast-AdenoCA, Lung-AdenoCA), have medium to high inferred WGD probability, compared to other cancer types. This might indicate that WGD indeed provides cancer a means to escape from nullisomy.

### Inferring selection parameters for driver genes in chromosomally stable tumors

Thus far we have examined the implementation of CINner in elucidating the selective roles of different CNA mechanisms, such as whole-chromosome and chromosome-arm missegregations and whole-genome duplication. Although these CNAs are frequently observed in cancer, most blood cancers and certain solid tumors are not associated with large-scale aneuploidies [9]. Here, we explore potential applications for CINner in analyzing such cancer types.

As a case study, we investigate the CLLE-ES cohort in PCAWG, consisting of chronic lymphocytic leukemia samples. The samples are largely diploid in general, therefore previously excluded in our chromosomal CNA study (**Fig. 2**). We hypothesize that the cancer type is driven mostly by mutations and focal gains and losses, suggesting that our selection model for driver genes is appropriate (**Fig. 1d**).

The list of driver genes is derived from tabulating all genes that are either mutated or impacted by CNAs in at least one CLLE-ES sample. Our selection model requires that each driver gene is assigned as either TSG or oncogene, under the assumption that losses of TSGs and gains of oncogenes are associated with increased fitness during tumorigenesis. Therefore, the driver genes are labeled as TSG or oncogene depending on whether the loss or gain frequency in the CLLE-ES cohort is higher, respectively. If the frequencies are equal, the driver gene role is taken from Cancer Gene Census [48]. We restrict the driver gene list to those that are listed in Cancer Gene Census, located on autosomes, and can be assigned a selective role.

We then model the lengths of focal amplification and deletion events separately, under the assumption that the ratio of a focal event length over the chromosome-arm length follows the Beta distribution (**Fig. 5a**). Because none of the driver genes in our list is affected by focal amplifications (**Fig. 5b**), we limit the parameter inference to driver mutation rate, focal deletion probability, and driver gene selection parameters. Similar to the CNA probability inference problem where the confounding effect of missegregation probabilities and chromosome-arm selection parameters results in nonidentifiability, here we fix the driver mutation rate and infer the other parameters relative to this value.

**Figure 5.**
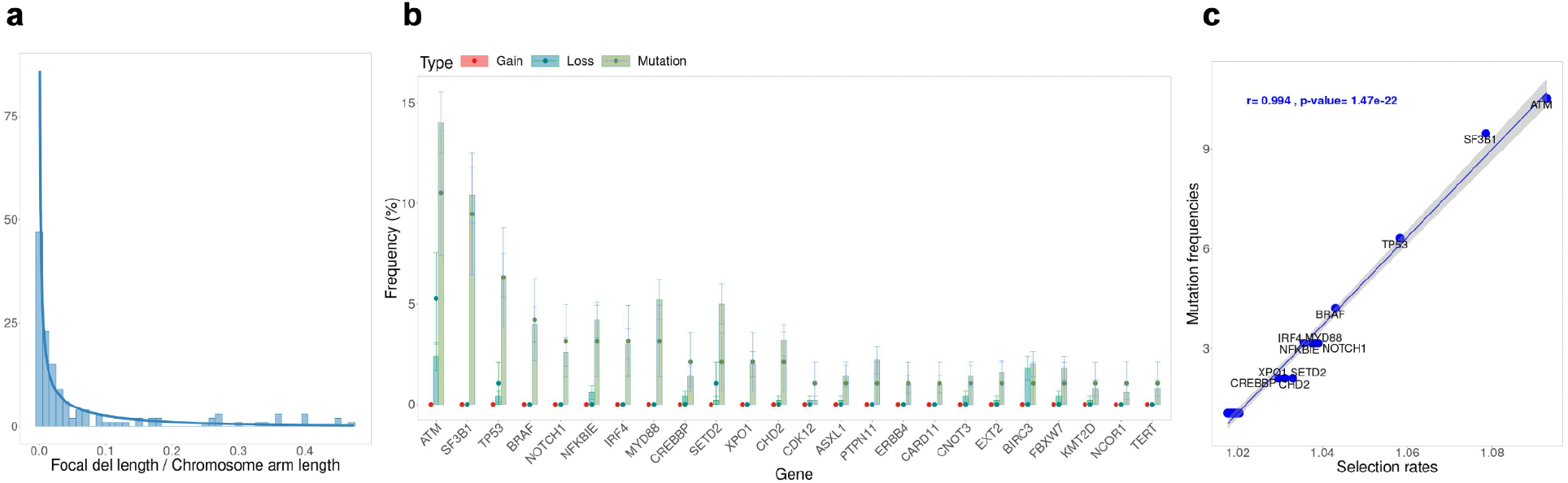
Results from fitting the driver gene selection model to the CLLE-ES cohort in PCAWG. **(a)** Ratios of focal deletion lengths over corresponding chromosome arm lengths are fitted with a Beta distribution. **(b)** Comparison between mutation/gain/loss frequencies among driver genes from the fitted model (bars) and PCAWG (circles). Error bars represent the standard deviations from 10,000 bootstrap samples. **(c)** Correlation between inferred selection rates and mutation frequencies for individual driver genes. Linear regressions and p-value from Pearson correlation.

All posterior distributions of the driver gene selection parameters are more concentrated than their prior distributions and are unimodal (**Supplementary Fig. 24**), confirming our ability to estimate the selection parameters effectively. Choosing the mode from each parameter’s posterior distribution, we compare the driver gene event frequencies from CINner against the PCAWG data (**Fig. 5b**). The mutation frequencies from CINner recover the signals from data. Moreover, the gene loss frequencies are largely in agreement with PCAWG observations. The inferred selection parameters exhibit a linear relationship with the mutation frequencies (**Fig. 5c**), similar to the correlation between CINner-inferred chromosome-arm selection parameters and frequencies of amplifications and deletions for cancer types driven by CNAs (**Fig. 2d-f**).

We note that currently the selection model is based on several simplifying assumptions. First, we assume haploinsufficiency for TSGs, i.e. cancer fitness increases with each additional allele inactivated via mutation or CN loss. For some driver genes, inactivation has been observed to require functional loss of both alleles (two-hit paradigm), and for yet other driver genes, the loss of one allele induces cancer but complete functional loss of the gene is toxic [23]. Second, our fitness formula is multiplicative, wherein each additional functional loss of a TSG or functional gain of an oncogene increases cell fitness by the same factor, dictated by the gene’s selection parameter. Third, in some cancer types, the fitness associated with a de novo gene alteration depends on the genomic landscape, indicating the importance of the order in which driver gene mutations and CNAs occur. This phenomenon is not included in our selection model. Although these assumptions greatly reduce the complexity of the parameter inference while still largely capturing the gene alteration patterns in CLLE-ES, applications in other data cohorts and cancer types might necessitate a more involved selection model.

## DISCUSSION

Cancer is characterized by a multistep development resulting in the reprogramming of key cellular components [49]. The genomic alterations that drive tumor heterogeneity and evolution range in extent and potency, from point mutations and small indels [50], to copy number aberrations and structural variants affecting one or multiple chromosomes simultaneously [43,51–53]. The occurrence rates of these distinct mutational processes and how they impact the selection landscape are highly context-dependent [39]. Loss or mutation of *TP53* and *BRCA1/2* leads to progressive increases in CNA rates and therefore tumor heterogeneity [54]. WGD likewise propagates chromosomal instability (CIN) [55], yet tolerance of WGD itself often also requires functional loss of *TP53* [43,56]. Different alleles of a TSG can be deactivated via mutation, deletion [57], or copy-neutral loss of heterozygosity [58]. However, the same mechanisms can constrain each other, as shown in a recent finding that 1q trisomy inhibits p53 signaling and accelerates tumor progression [59].

Because the effects of CNAs on cancer depend heavily on context, it is challenging to fully understand the development of chromosomal instability from DNA data statistics alone. Mathematical modeling can help distinguish between the effects of selection and neutral drift in CIN evolution, reveal how preference for specific karyotypes shapes cancer evolution, and forecast clonal dynamics [16]. Moreover, if both the occurrence rate and selection parameter of a CNA can be ascertained, we can infer the allele age and reconstruct the sample coalescence [60], which can offer valuable information in diagnosis and treatment selection.

To provide a comprehensive picture of how CIN impacts tumor progression, a model must possess two important features. First, it should account for the opposing forces of diversity and selection. Second, it should allow for the coexistence and interaction of different mutation and CNA mechanisms. Most models so far have only addressed some of these aspects. Early works studied chromosome copy number changes without selection, only assuming that nullisomic karyotypes were nonviable [61,62]. A recent model defined different phases of CIN in tumor growth, using breakpoint tally to measure subclone fitness [63]. Since copy number is not explicitly defined, this approach seems unsuitable for analyzing selection for optimal karyotypes. Another CIN study [64] employs CINSignatureGenomeSimulation [65], an algorithm to simulate the effects of different copy number signatures on the cancer genome, without direct modeling of selection. Other methods first generate a cell genealogy, then simulate CNAs along the branches [66,67]. However, the CNAs in this approach do not affect the phylogeny tree, therefore karyotype selection is not explicitly depicted. Some recent studies model the effects of selection and missegregation on subclonal copy numbers [18,68], also incorporating point mutations [13] or WGD [46,69]. Nevertheless, many of these models focus only on the average ploidy and do not consider chromosome-specific CN [13,46,68]. In contrast, two studies directly integrate chromosomal selection parameters and study the resulting CN trajectories [18,69]. They employ existing chromosomal selection parameters defined from counts and potencies of OGs and TSGs from pan-cancer studies [22], to uncover the prevalent karyotypic trajectories during tumorigenesis. This seems to be the most promising approach to study the role of heterogeneity and selection in cancer on the copy number level. However, a potential drawback is that identification of cancer driver genes is nontrivial and its sensitivity depends on the sample size as well as the genes’ mutation frequency, among other factors [70]. Therefore, although defining selection parameters based on known OGs and TSGs can give accurate results in a pan-cancer context, the approach seems to have limited applicability for studies of specific cancer types and datasets.

We present CINner, a model for simulating CNA mechanisms and selection in tumor evolution. It can accommodate various genomic events ranging in extent and impact, from point mutations to WGD. CINner uses several numerical techniques to reduce the memory and computing requirements of simulating whole genomes in large cancer populations. We use CINner to find chromosome-arm selection parameters from diploid PCAWG samples. The CN profiles simulated with the inferred parameters match the observed cancer-specific aneuploidy patterns. The estimated selection parameters predict WGD prevalence and correlate with driver gene count and potency in pan-cancer TCGA data. These signals indicate that the selection parameters inferred from CINner reflect the oncogenic effects of genes on specific genomic regions. Therefore, CINner can play an important role in modeling rare cancers, where driver gene identification is limited due to small sample size and low gene alteration frequencies. We also perform parameter analysis studies to quantify the effects of CNA probabilities, selection parameters, tissue cell count and tumor growth dynamics on CINner statistics. Finally, we apply CINner to cancers driven mainly by driver gene changes, such as CLLE-ES in PCAWG. In short, CINner is capable of modeling both small alterations impacting important genes and large-scale CNAs during tumor development.

An interesting finding from our parameter inference is that the WGD prevalence of a cancer type is connected to its chromosomal selection parameters in the diploid setting. This provides some insights into why WGD is a common early event in tumors, despite strong negative selection in normal tissues. In particular, WGD proportion correlates with higher selection parameters of TSG-acting chromosome arms (**Fig. 2g**). As deletions of these arms are strongly selective, the cancer cells gradually lose copies due to missegregations, thereby risking the toxic effect of nullisomy. WGD could help alleviate this risk by raising those chromosome copy numbers above 1. Another explanation is that, because of the increased ploidy level, tetraploid cells can have repeated losses of a chromosome, resulting in higher impact on the gene balance. The WGD proportion also correlates strongly with the count of chromosome arms acting as either TSGs or oncogenes (**Fig. 2h**). This could be because different cancer and tissue types have different levels of tolerance for aneuploidy and WGD. Alternatively, WGD has been shown to promote chromosomal instability [71]. The tetraploid cells, therefore, can explore the aneuploidy landscape and increase their fitness at a faster rate than diploid cells.

Although CINner has the power to study clonal dynamics at the single-cell level, the parameters were estimated by comparing pseudo-bulk simulations to bulk DNA-sequencing data. This is to take advantage of the large sample sizes available with PCAWG and other cancer studies, to reduce the risk of overfitting. However, as shown in our parameter studies (**Fig. 3g**), the chromosome gain and loss frequencies in bulk samples are similarly impacted by CNA probabilities and selection parameters. This makes it challenging to infer both CNA probabilities and selection parameters simultaneously. Therefore, in this work, we fix the missegregation probabilities and focus on finding cancer-specific selection parameters.

Recently, technological advances in single-cell DNA sequencing have led to better resolution in capturing genomic data, and have demonstrated that tumors exhibit different levels of heterogeneity and chromosomal instability [54,63,72,73]. Our parameter studies show that single-cell statistics have different trends under variable CNA probabilities and selection parameters in CINner (**Fig. 3e-h**). While higher missegregation probability results in increasing aneuploidy and sample diversity, higher selection parameters increase aneuploidy but decrease clone count, as a result of heightened subclonal competition. Therefore, as single-cell data increases in sample size and cell count, CINner can be implemented to estimate both the occurrence rate and fitness impact of different CNA mechanisms, circumventing the nonidentifiability issues in bulk data. In conclusion, we have shown that CINner offers a comprehensive framework to analyze the interplay between selection and distinct genomic alteration mechanisms. CINner can simulate individual cells and clones in a sample, making it adaptable for DNA data ranging from single-cell to bulk level. Its flexibility can accommodate data from different DNA-seq technologies, including targeted sequencing [74,75], and enable incorporation of new CNA mechanisms or point mutations [76]. We envision that with the advent of large genomic studies using both bulk and single-cell approaches, CINner will enable accurate parameterization of cancer evolution.

## CODE AVAILABILITY

CINner is available as an R package at https://github.com/dinhngockhanh/CINner.

## ACKNOWLEDGEMENTS

This work was funded in part by an Ovarian Cancer Research Alliance (OCRA) Ann Schreiber Mentored Investigator Award to IVG [650687]. ACW is supported by NCI Ruth L. Kirschstein National Research Service Award for Predoctoral Fellows F31-CA271673. RM was supported by the Computational Biology Summer Program at Memorial Sloan Kettering Cancer Center, funded by an R25 grant from the National Institutes of Health [5R25CA233208-04].

## SUPPLEMENTARY INFORMATION

**Supplementary Figure 1.**
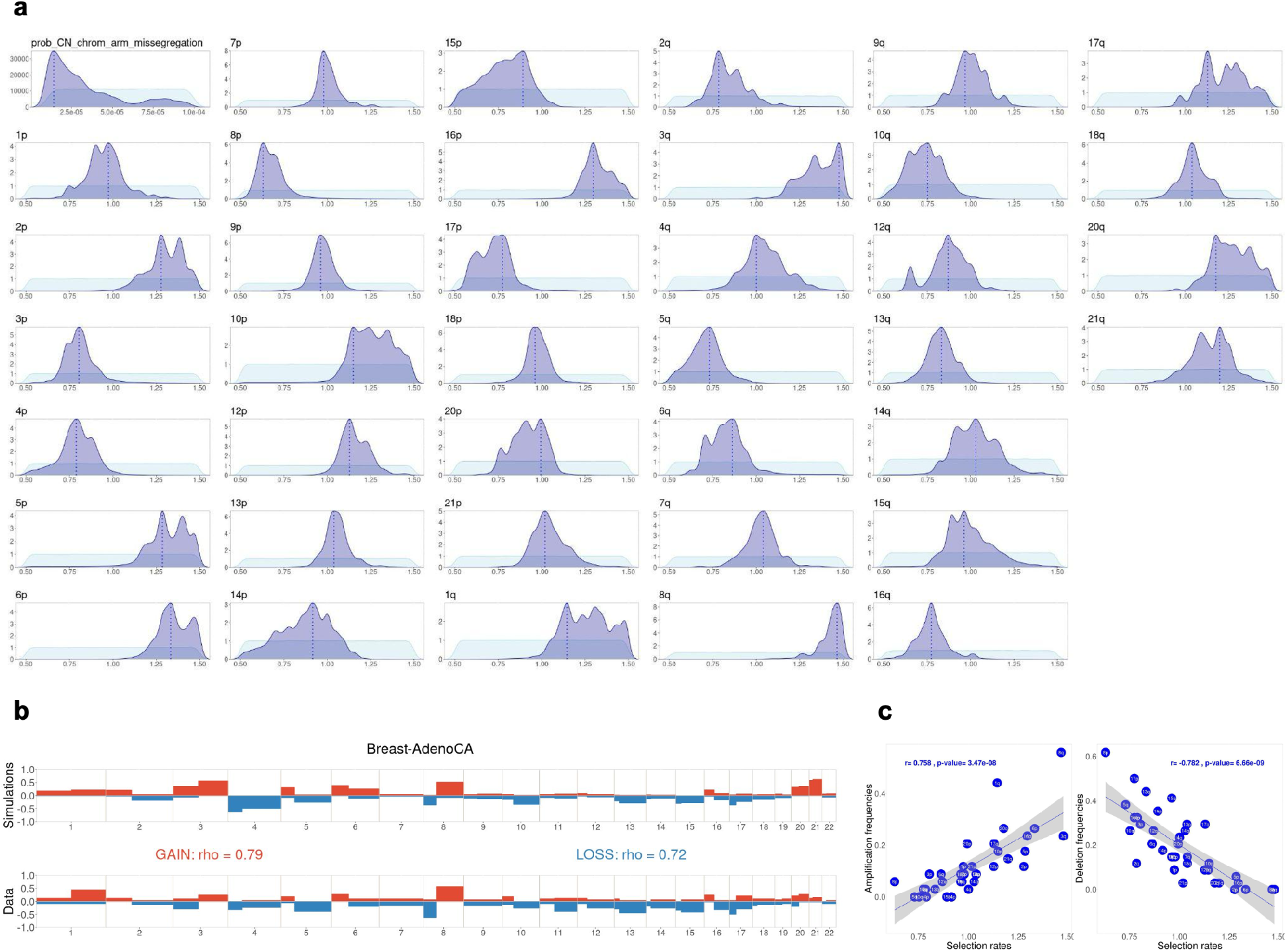
Inference of chromosome-arm selection rates in Breast-AdenoCA (PCAWG). **(a)** Prior distribution (light blue) and posterior distribution (dark blue) from inference with ABC random forest. Broken line represents the mode in the posterior distribution for each parameter. **(b)** Comparison between simulations with fitted parameter (top) and gain/loss frequencies at arm level from TCGA (bottom). The simulations are computed with the posterior modes from **(a)**. Spearman’s correlation coefficient rho between frequencies of gains (or losses) among each arm in PCAWG and simulations. **(c)** Correlation between inferred selection rates and amplification/deletion frequencies for individual chromosome arms. Linear regressions and p-values from Pearson correlation.

**Supplementary Figure 2.**
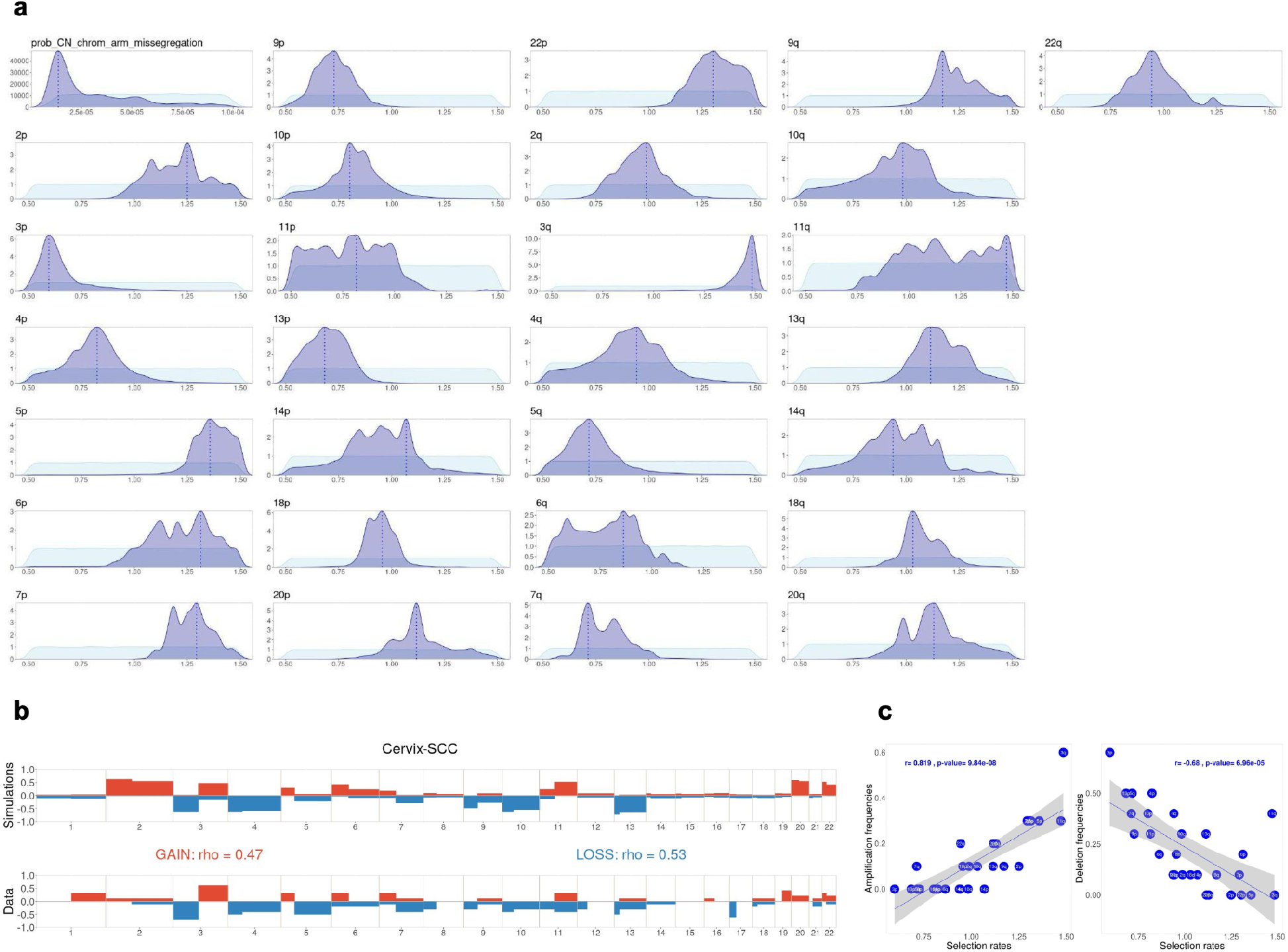
Inference of chromosome-arm selection rates in Cervix-SCC (PCAWG). **(a)** Prior distribution (light blue) and posterior distribution (dark blue) from inference with ABC random forest. Broken line represents the mode in the posterior distribution for each parameter. **(b)** Comparison between simulations with fitted parameter (top) and gain/loss frequencies at arm level from TCGA (bottom). The simulations are computed with the posterior modes from **(a)**. Spearman’s correlation coefficient rho between frequencies of gains (or losses) among each arm in PCAWG and simulations. **(c)** Correlation between inferred selection rates and amplification/deletion frequencies for individual chromosome arms. Linear regressions and p-values from Pearson correlation.

**Supplementary Figure 3.**
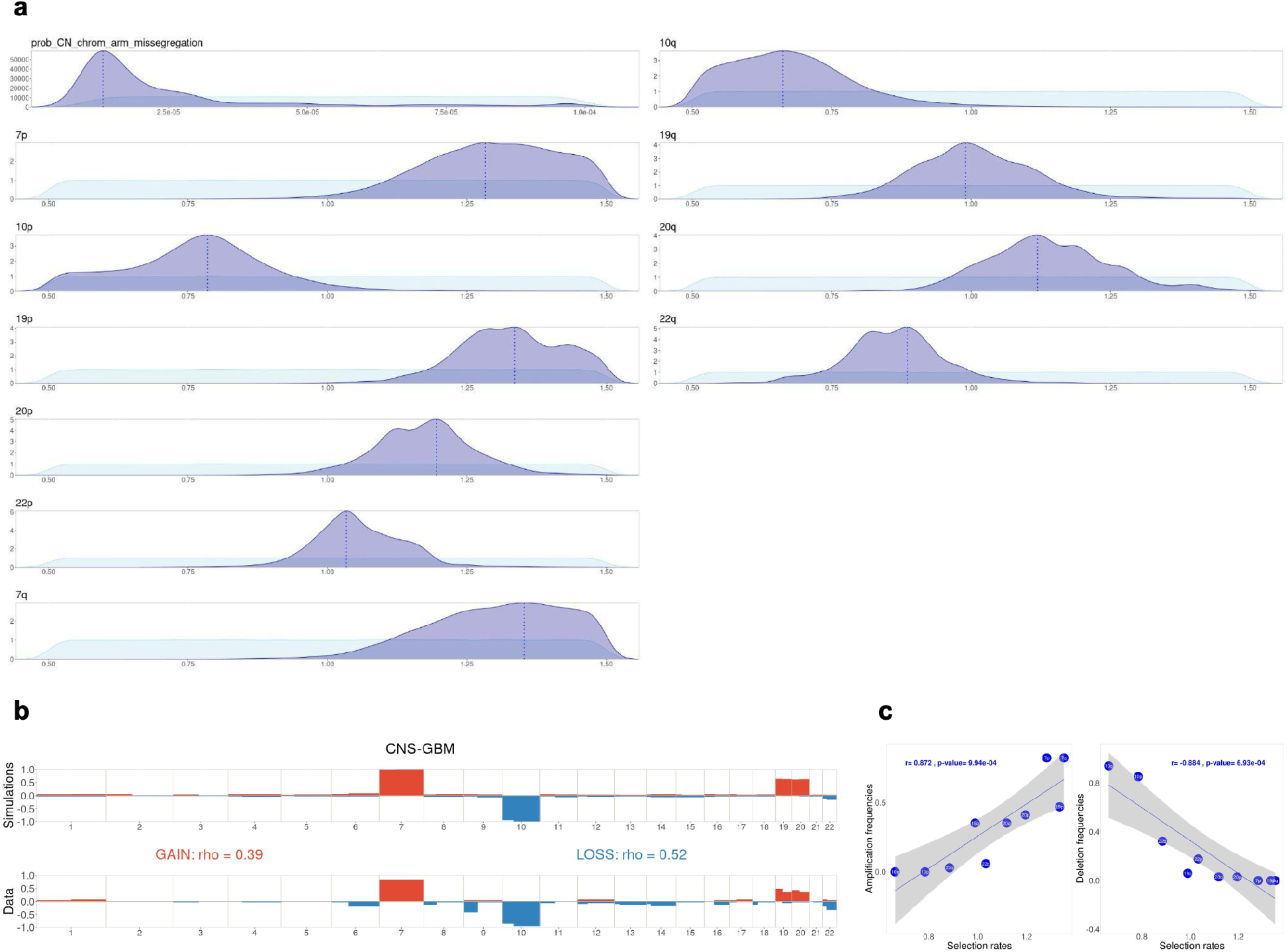
Inference of chromosome-arm selection rates in CNS-GBM (PCAWG). **(a)** Prior distribution (light blue) and posterior distribution (dark blue) from inference with ABC random forest. Broken line represents the mode in the posterior distribution for each parameter. **(b)** Comparison between simulations with fitted parameter (top) and gain/loss frequencies at arm level from TCGA (bottom). The simulations are computed with the posterior modes from **(a)**. Spearman’s correlation coefficient rho between frequencies of gains (or losses) among each arm in PCAWG and simulations. **(c)** Correlation between inferred selection rates and amplification/deletion frequencies for individual chromosome arms. Linear regressions and p-values from Pearson correlation.

**Supplementary Figure 4.**
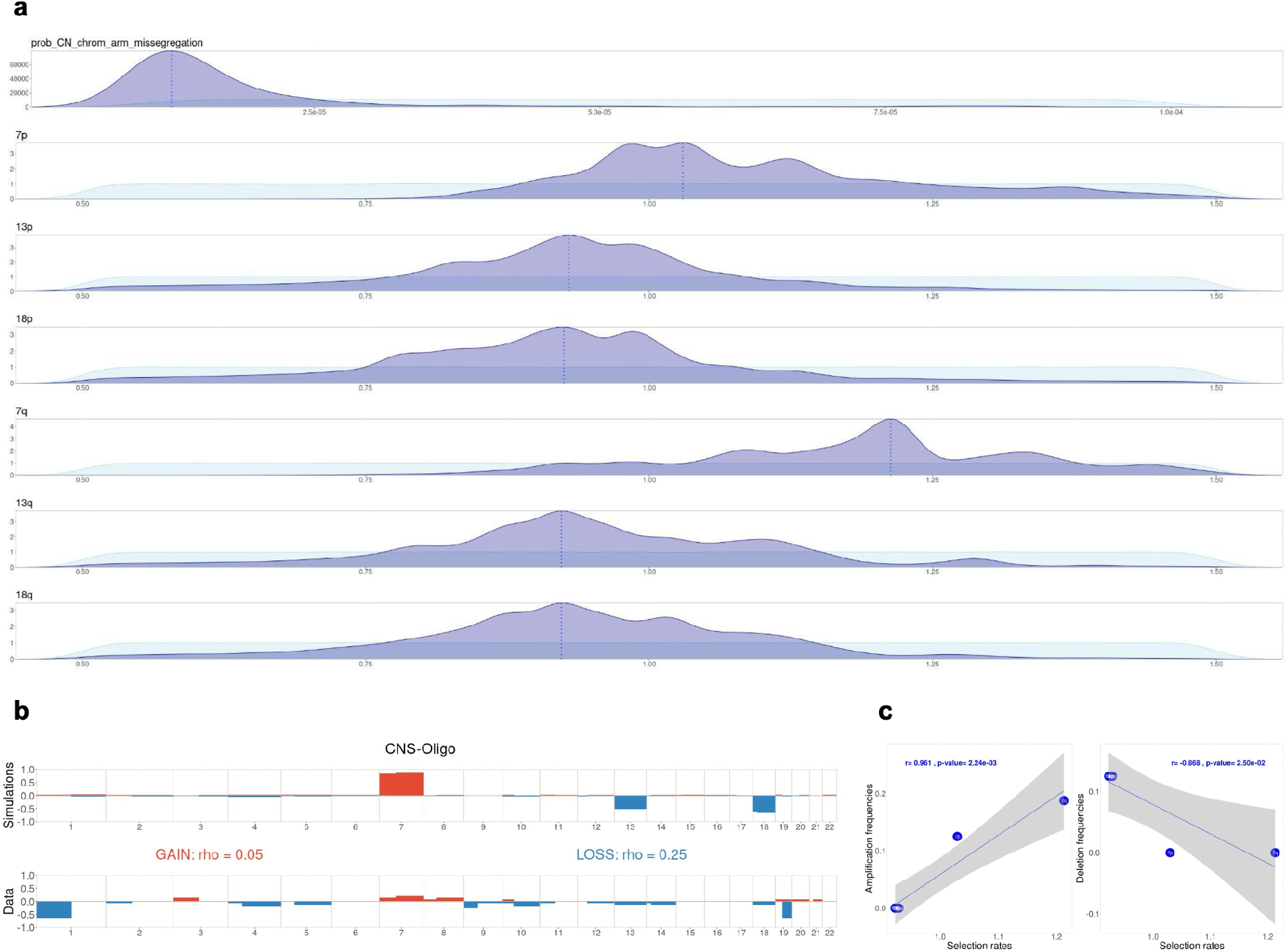
Inference of chromosome-arm selection rates in CNS-Oligo (PCAWG). **(a)** Prior distribution (light blue) and posterior distribution (dark blue) from inference with ABC random forest. Broken line represents the mode in the posterior distribution for each parameter. **(b)** Comparison between simulations with fitted parameter (top) and gain/loss frequencies at arm level from TCGA (bottom). The simulations are computed with the posterior modes from **(a)**. Spearman’s correlation coefficient rho between frequencies of gains (or losses) among each arm in PCAWG and simulations. **(c)** Correlation between inferred selection rates and amplification/deletion frequencies for individual chromosome arms. Linear regressions and p-values from Pearson correlation.

**Supplementary Figure 5.**
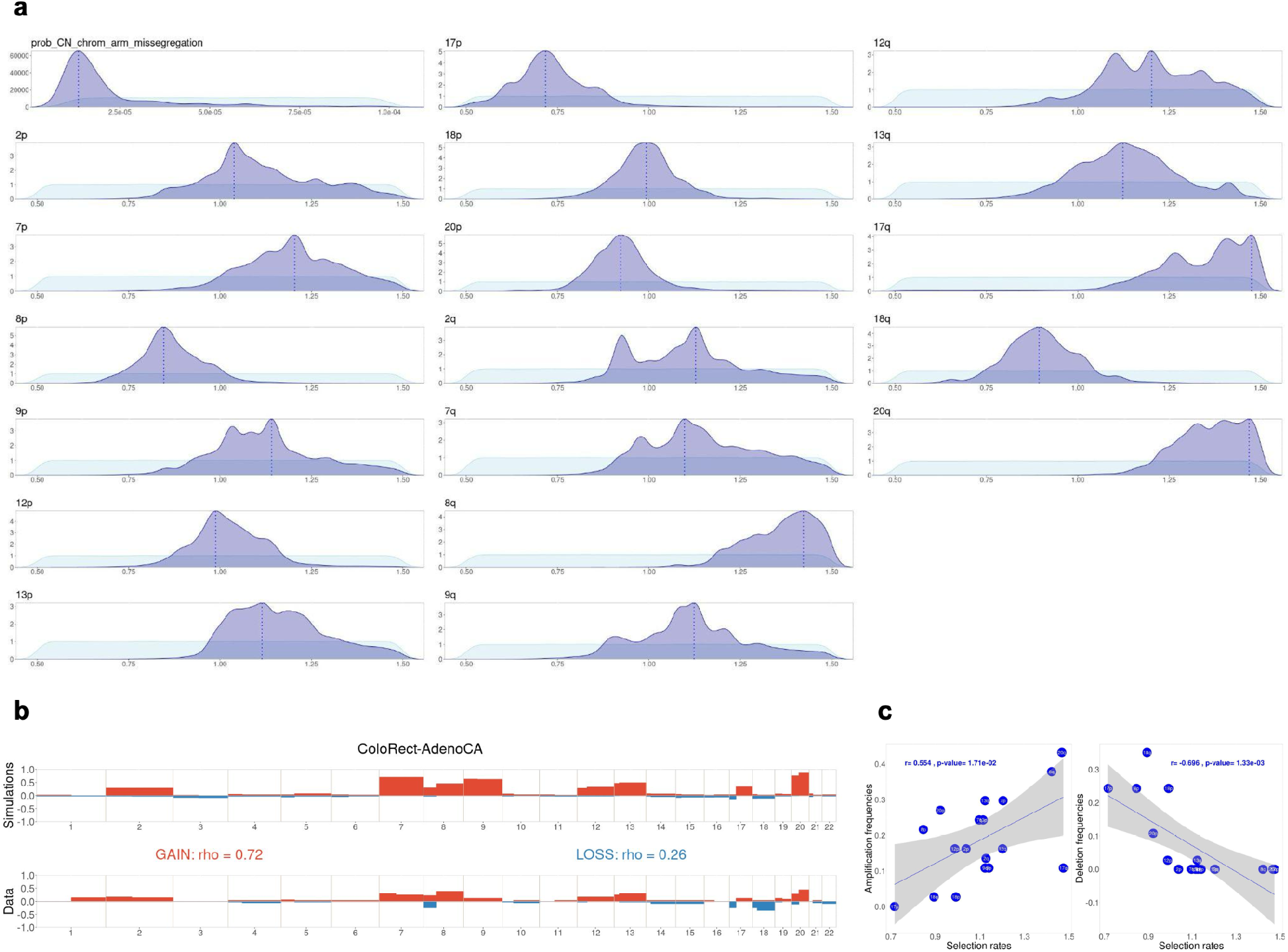
Inference of chromosome-arm selection rates in ColoRect-AdenoCA (PCAWG). **(a)** Prior distribution (light blue) and posterior distribution (dark blue) from inference with ABC random forest. Broken line represents the mode in the posterior distribution for each parameter. **(b)** Comparison between simulations with fitted parameter (top) and gain/loss frequencies at arm level from TCGA (bottom). The simulations are computed with the posterior modes from **(a)**. Spearman’s correlation coefficient rho between frequencies of gains (or losses) among each arm in PCAWG and simulations. **(c)** Correlation between inferred selection rates and amplification/deletion frequencies for individual chromosome arms. Linear regressions and p-values from Pearson correlation.

**Supplementary Figure 6.**
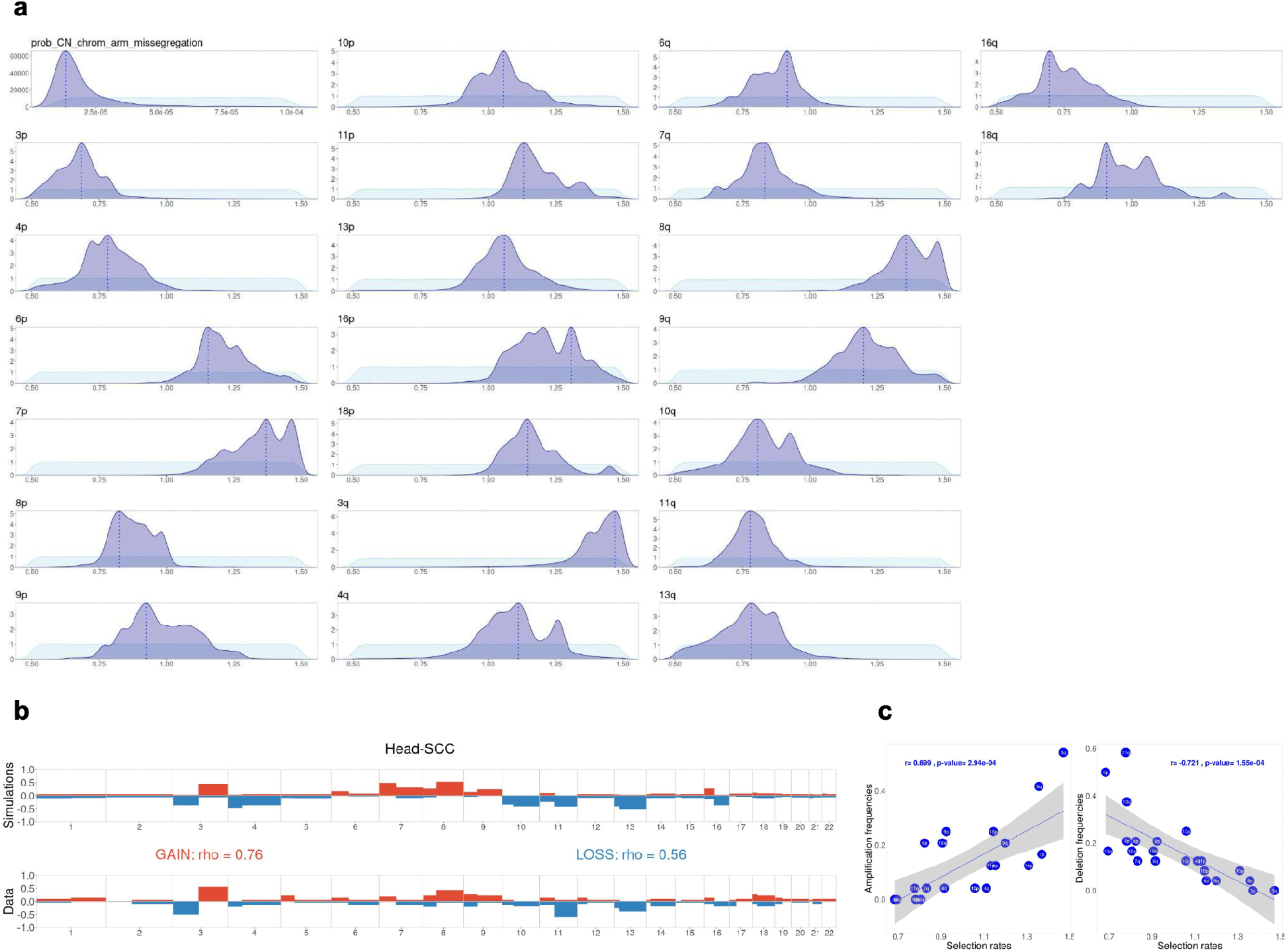
Inference of chromosome-arm selection rates in Head-SCC (PCAWG). **(a)** Prior distribution (light blue) and posterior distribution (dark blue) from inference with ABC random forest. Broken line represents the mode in the posterior distribution for each parameter. **(b)** Comparison between simulations with fitted parameter (top) and gain/loss frequencies at arm level from TCGA (bottom). The simulations are computed with the posterior modes from **(a)**. Spearman’s correlation coefficient rho between frequencies of gains (or losses) among each arm in PCAWG and simulations. **(c)** Correlation between inferred selection rates and amplification/deletion frequencies for individual chromosome arms. Linear regressions and p-values from Pearson correlation.

**Supplementary Figure 7.**
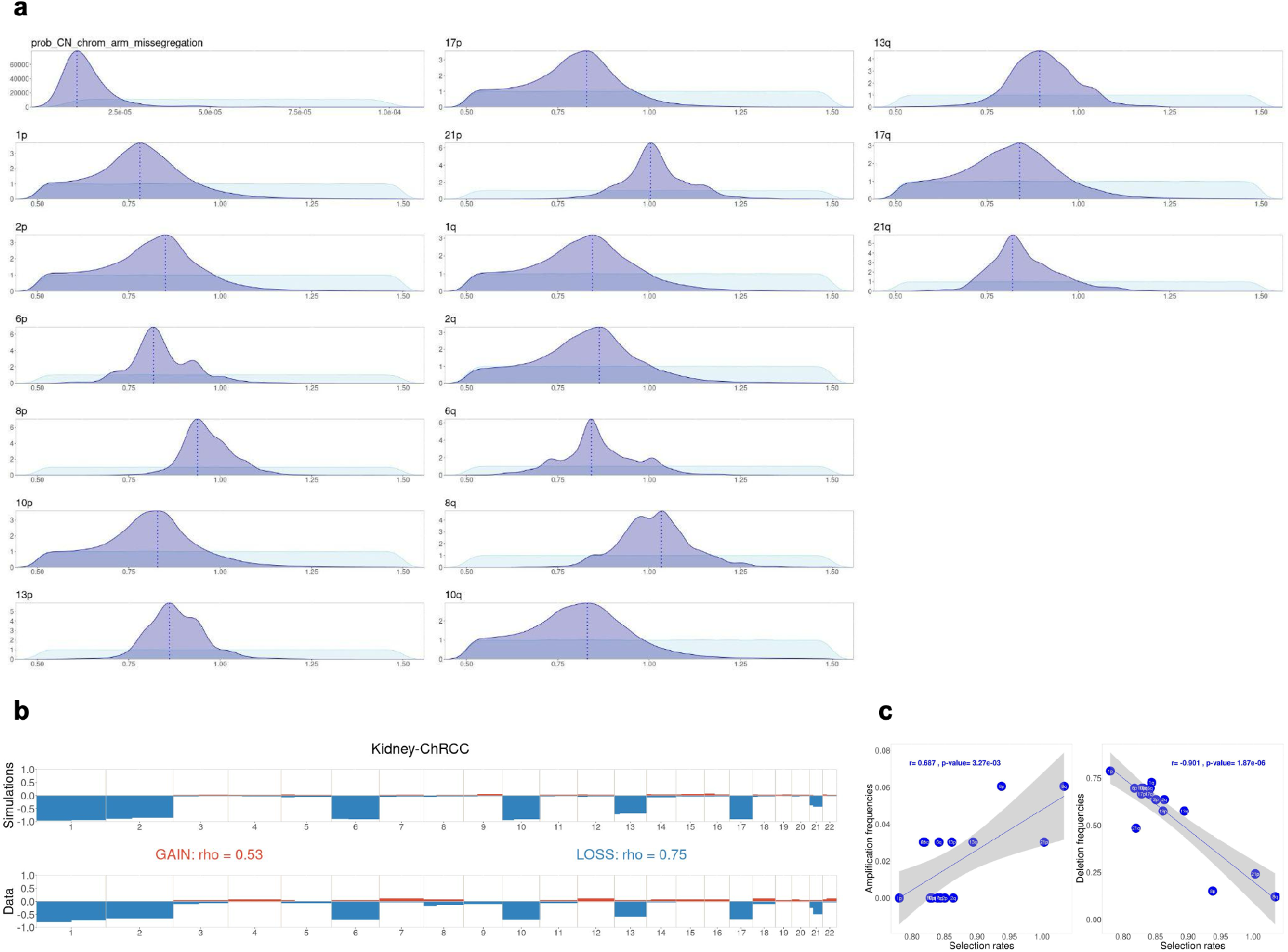
Inference of chromosome-arm selection rates in Kidney-ChRCC (PCAWG). **(a)** Prior distribution (light blue) and posterior distribution (dark blue) from inference with ABC random forest. Broken line represents the mode in the posterior distribution for each parameter. **(b)** Comparison between simulations with fitted parameter (top) and gain/loss frequencies at arm level from TCGA (bottom). The simulations are computed with the posterior modes from **(a)**. Spearman’s correlation coefficient rho between frequencies of gains (or losses) among each arm in PCAWG and simulations. **(c)** Correlation between inferred selection rates and amplification/deletion frequencies for individual chromosome arms. Linear regressions and p-values from Pearson correlation.

**Supplementary Figure 8.**
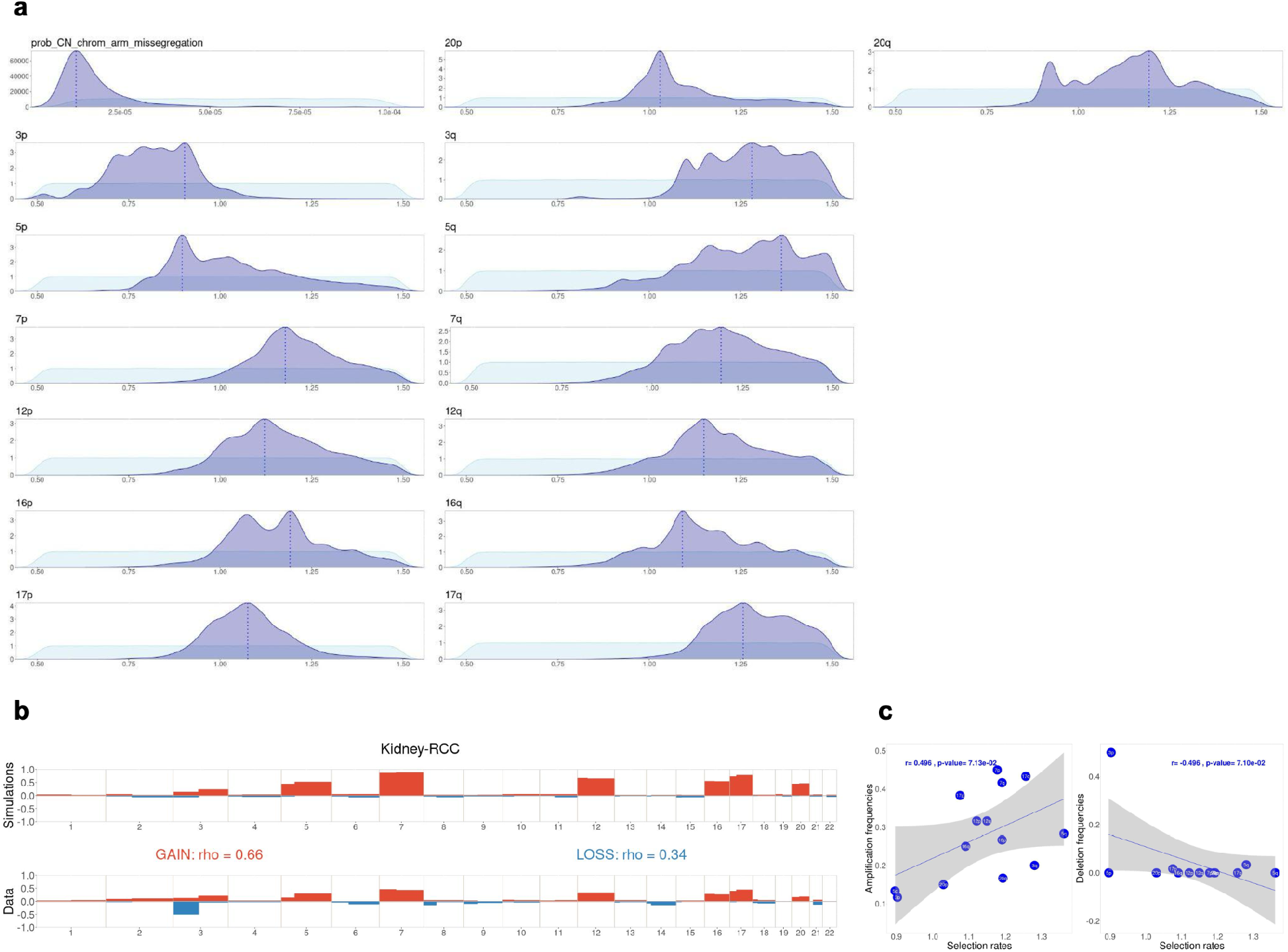
Inference of chromosome-arm selection rates in Kidney-RCC (PCAWG). **(a)** Prior distribution (light blue) and posterior distribution (dark blue) from inference with ABC random forest. Broken line represents the mode in the posterior distribution for each parameter. **(b)** Comparison between simulations with fitted parameter (top) and gain/loss frequencies at arm level from TCGA (bottom). The simulations are computed with the posterior modes from **(a)**. Spearman’s correlation coefficient rho between frequencies of gains (or losses) among each arm in PCAWG and simulations. **(c)** Correlation between inferred selection rates and amplification/deletion frequencies for individual chromosome arms. Linear regressions and p-values from Pearson correlation.

**Supplementary Figure 9.**
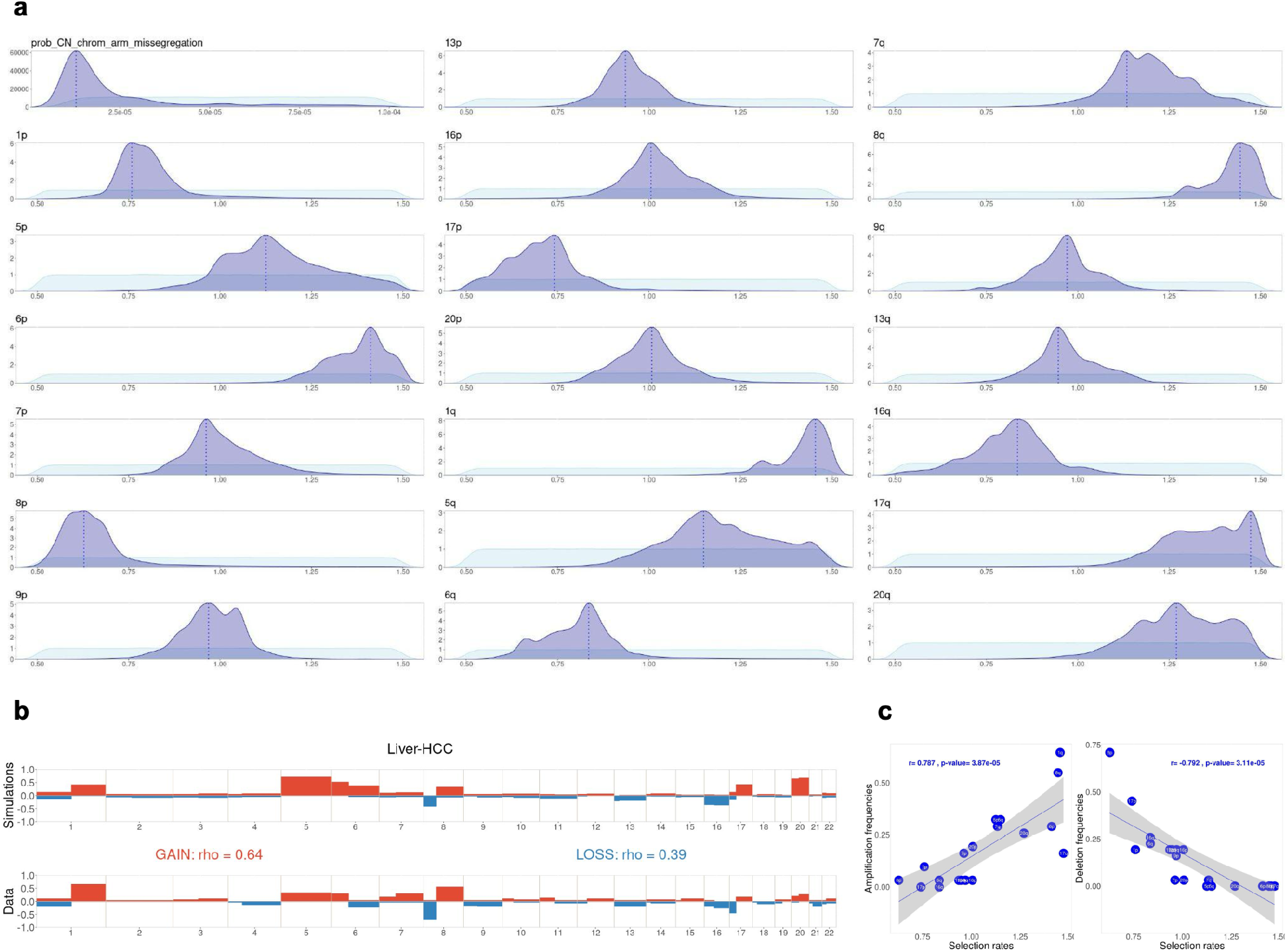
Inference of chromosome-arm selection rates in Liver-HCC (PCAWG). **(a)** Prior distribution (light blue) and posterior distribution (dark blue) from inference with ABC random forest. Broken line represents the mode in the posterior distribution for each parameter. **(b)** Comparison between simulations with fitted parameter (top) and gain/loss frequencies at arm level from TCGA (bottom). The simulations are computed with the posterior modes from **(a)**. Spearman’s correlation coefficient rho between frequencies of gains (or losses) among each arm in PCAWG and simulations. **(c)** Correlation between inferred selection rates and amplification/deletion frequencies for individual chromosome arms. Linear regressions and p-values from Pearson correlation.

**Supplementary Figure 10.**
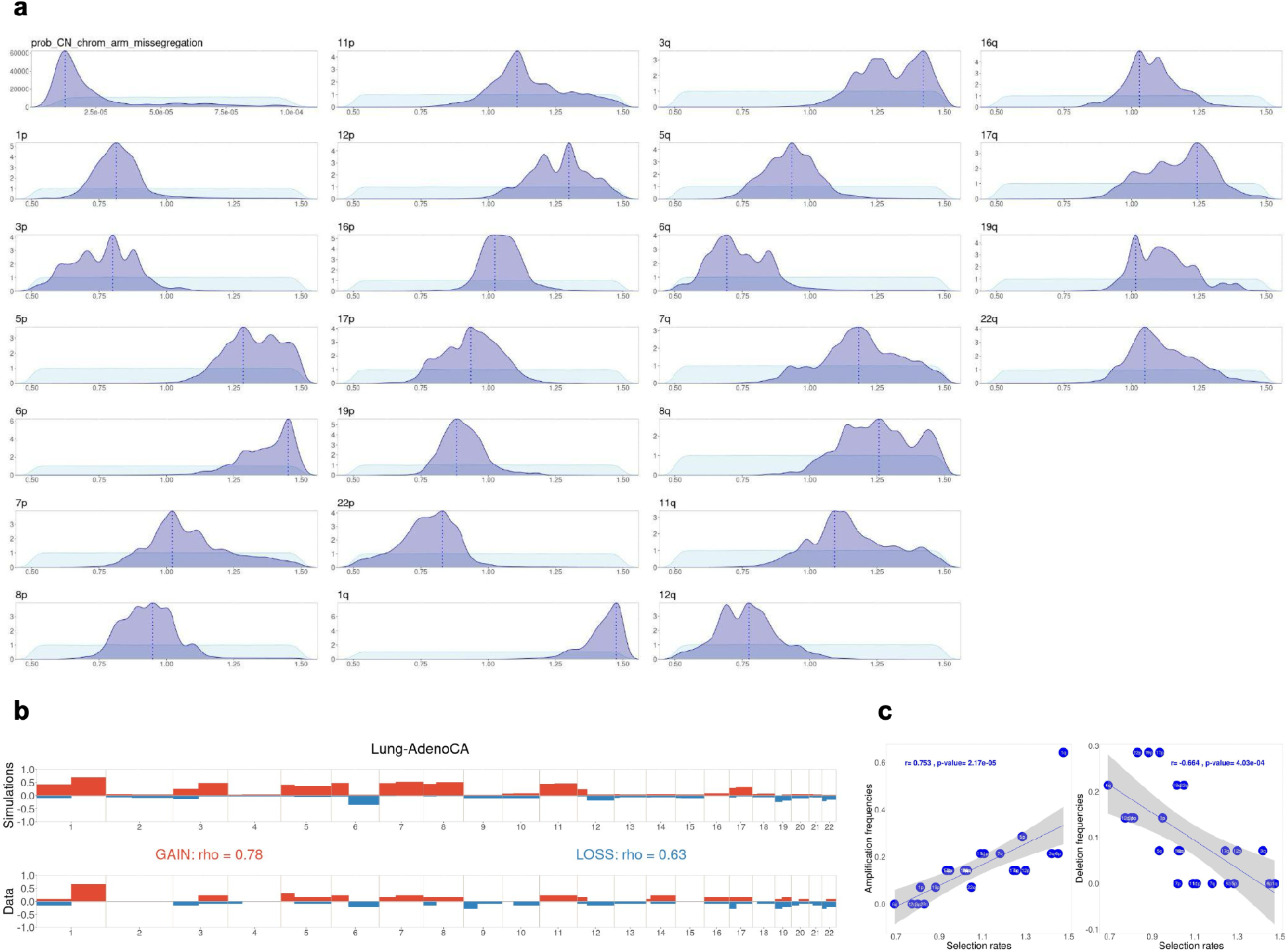
Inference of chromosome-arm selection rates in Lung-AdenoCA (PCAWG). **(a)** Prior distribution (light blue) and posterior distribution (dark blue) from inference with ABC random forest. Broken line represents the mode in the posterior distribution for each parameter. **(b)** Comparison between simulations with fitted parameter (top) and gain/loss frequencies at arm level from TCGA (bottom). The simulations are computed with the posterior modes from **(a)**. Spearman’s correlation coefficient rho between frequencies of gains (or losses) among each arm in PCAWG and simulations. **(c)** Correlation between inferred selection rates and amplification/deletion frequencies for individual chromosome arms. Linear regressions and p-values from Pearson correlation.

**Supplementary Figure 11.**
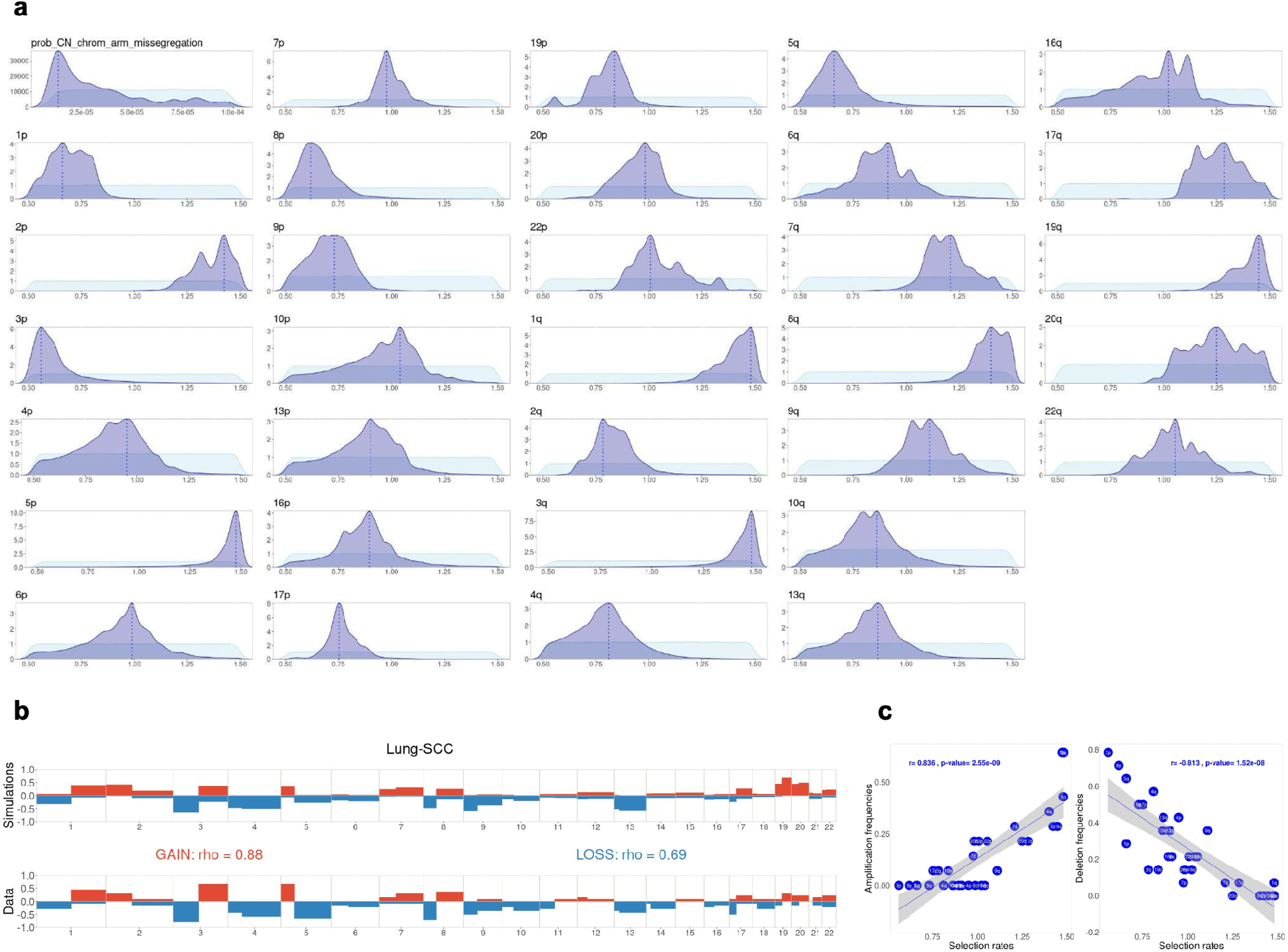
Inference of chromosome-arm selection rates in Lung-SCC (PCAWG). **(a)** Prior distribution (light blue) and posterior distribution (dark blue) from inference with ABC random forest. Broken line represents the mode in the posterior distribution for each parameter. **(b)** Comparison between simulations with fitted parameter (top) and gain/loss frequencies at arm level from TCGA (bottom). The simulations are computed with the posterior modes from **(a)**. Spearman’s correlation coefficient rho between frequencies of gains (or losses) among each arm in PCAWG and simulations. **(c)** Correlation between inferred selection rates and amplification/deletion frequencies for individual chromosome arms. Linear regressions and p-values from Pearson correlation.

**Supplementary Figure 12.**
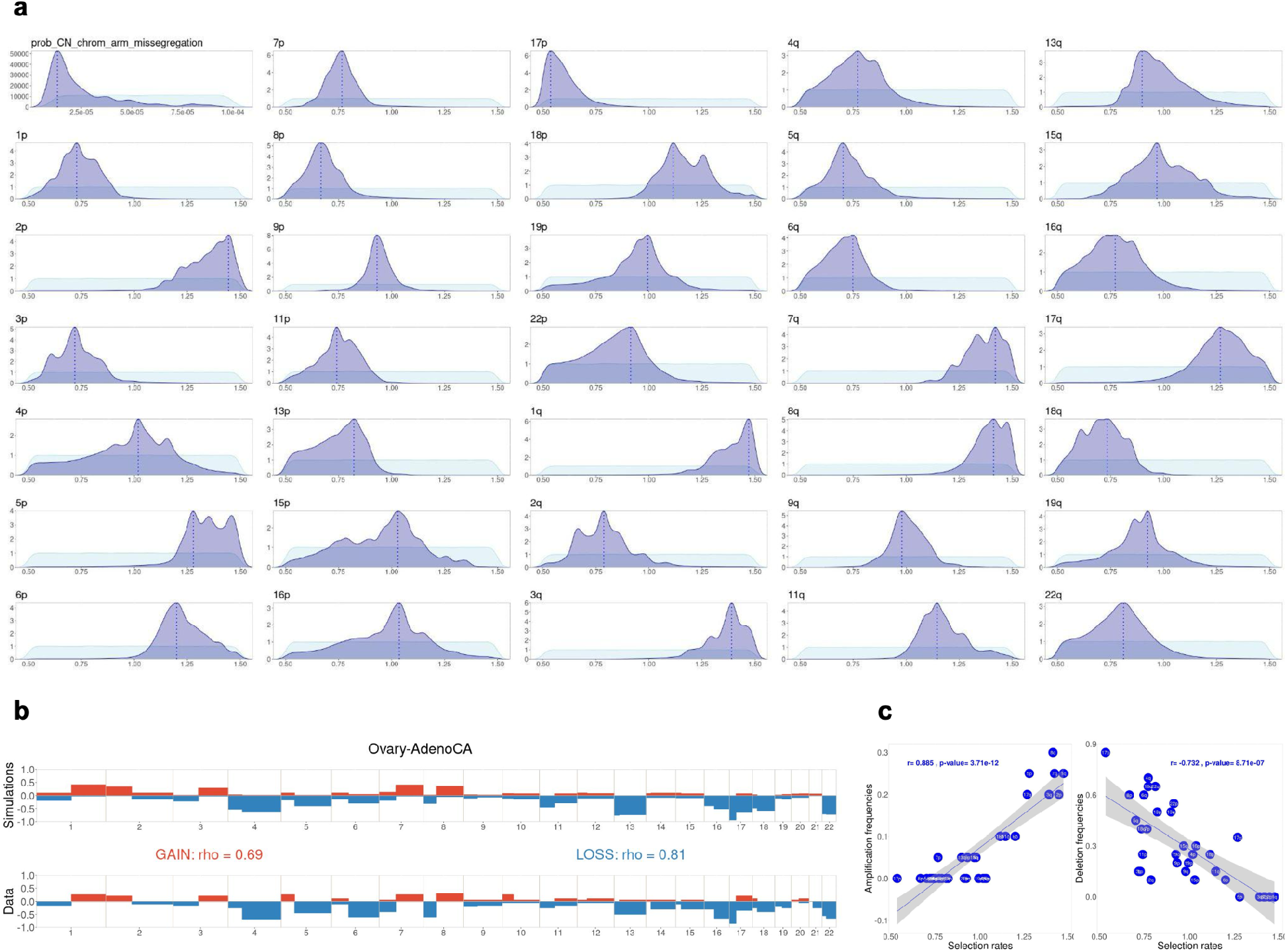
Inference of chromosome-arm selection rates in Ovary-AdenoCA (PCAWG). **(a)** Prior distribution (light blue) and posterior distribution (dark blue) from inference with ABC random forest. Broken line represents the mode in the posterior distribution for each parameter. **(b)** Comparison between simulations with fitted parameter (top) and gain/loss frequencies at arm level from TCGA (bottom). The simulations are computed with the posterior modes from **(a)**. Spearman’s correlation coefficient rho between frequencies of gains (or losses) among each arm in PCAWG and simulations. **(c)** Correlation between inferred selection rates and amplification/deletion frequencies for individual chromosome arms. Linear regressions and p-values from Pearson correlation.

**Supplementary Figure 13.**
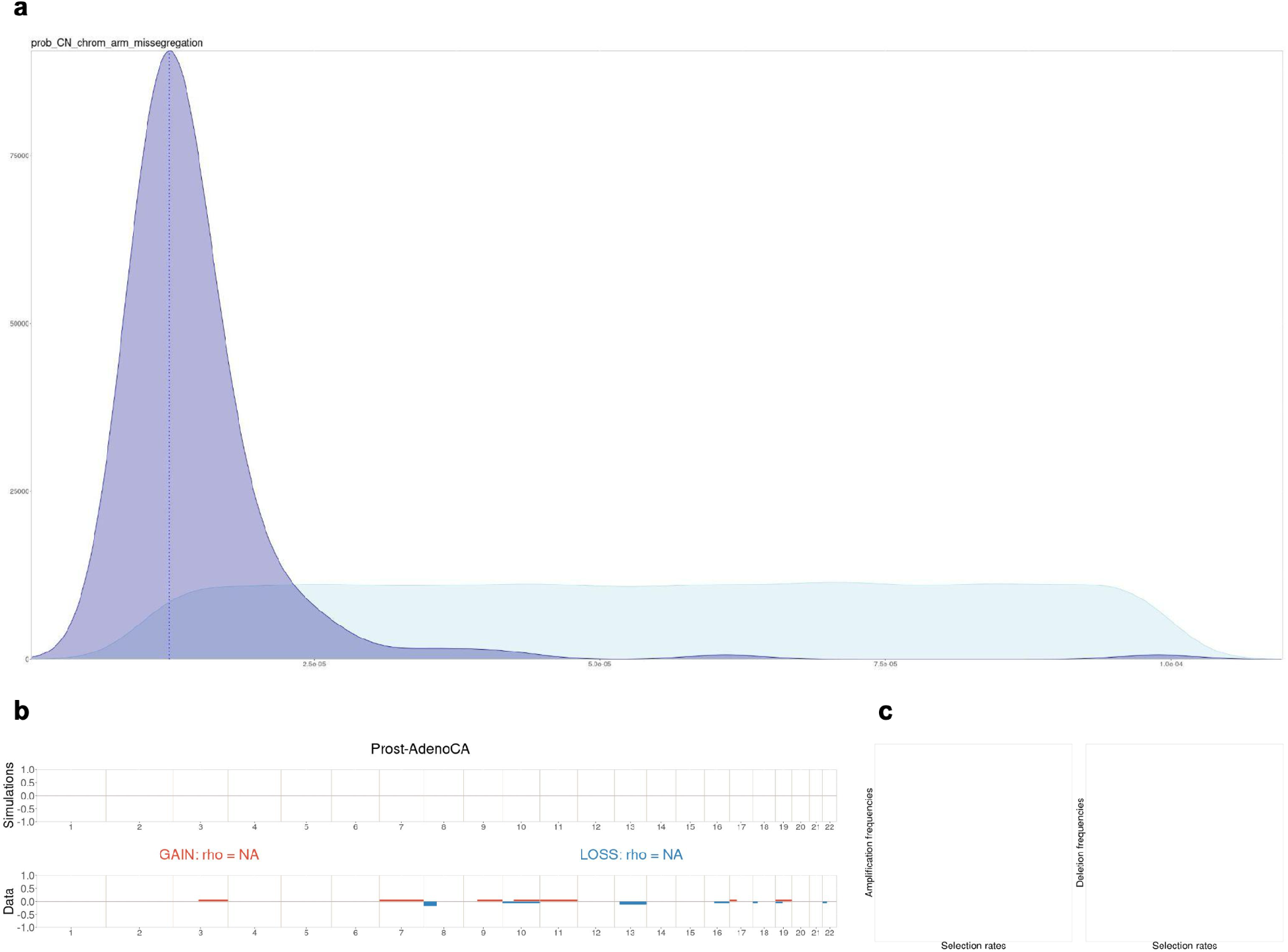
Inference of chromosome-arm selection rates in Prost-AdenoCA (PCAWG). **(a)** Prior distribution (light blue) and posterior distribution (dark blue) from inference with ABC random forest. Broken line represents the mode in the posterior distribution for each parameter. **(b)** Comparison between simulations with fitted parameter (top) and gain/loss frequencies at arm level from TCGA (bottom). The simulations are computed with the posterior modes from **(a)**. Spearman’s correlation coefficient rho between frequencies of gains (or losses) among each arm in PCAWG and simulations. **(c)** Correlation between inferred selection rates and amplification/deletion frequencies for individual chromosome arms. Linear regressions and p-values from Pearson correlation.

**Supplementary Figure 14.**
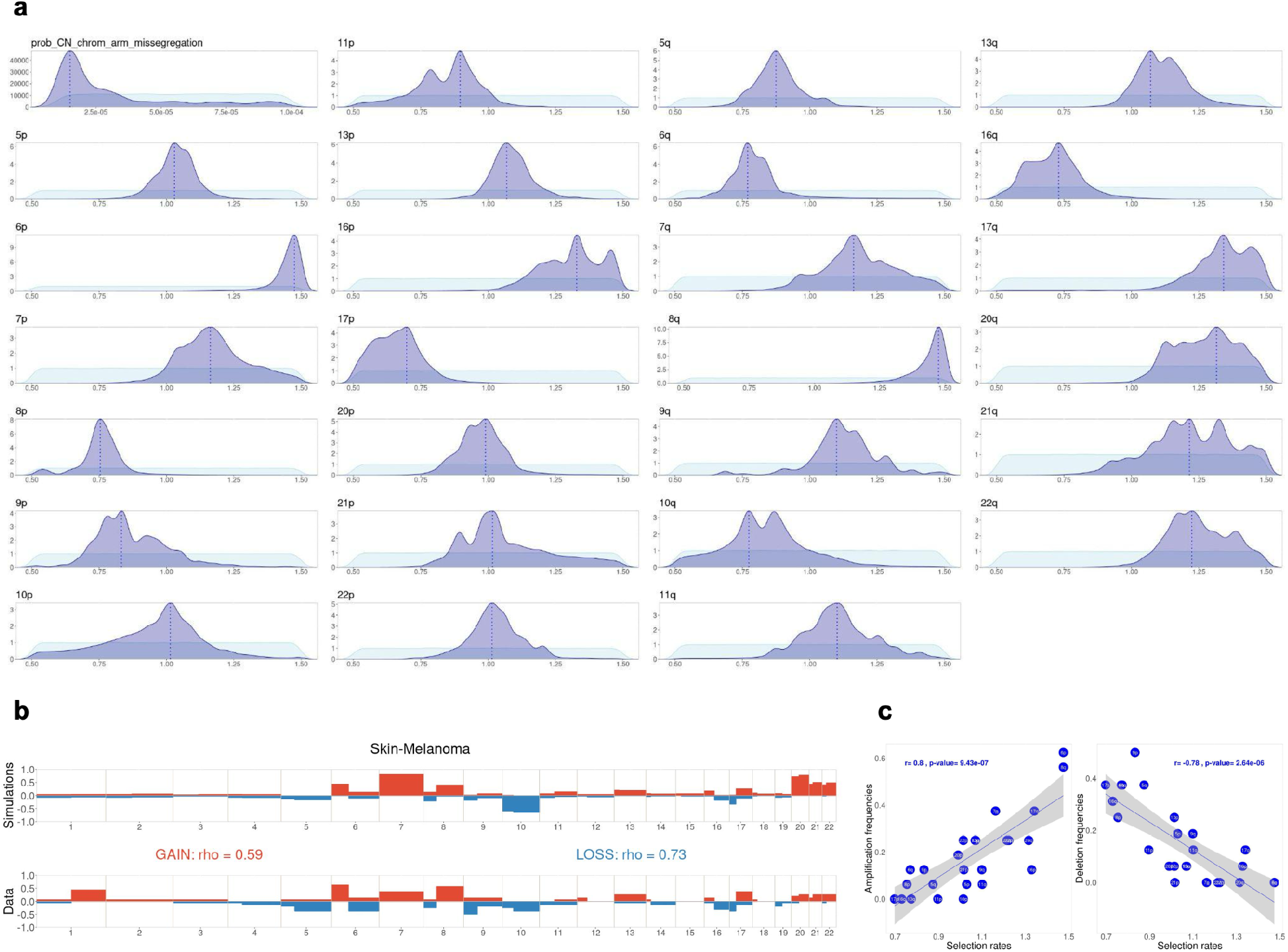
Inference of chromosome-arm selection rates in Skin-Melanoma (PCAWG). **(a)** Prior distribution (light blue) and posterior distribution (dark blue) from inference with ABC random forest. Broken line represents the mode in the posterior distribution for each parameter. **(b)** Comparison between simulations with fitted parameter (top) and gain/loss frequencies at arm level from TCGA (bottom). The simulations are computed with the posterior modes from **(a)**. Spearman’s correlation coefficient rho between frequencies of gains (or losses) among each arm in PCAWG and simulations. **(c)** Correlation between inferred selection rates and amplification/deletion frequencies for individual chromosome arms. Linear regressions and p-values from Pearson correlation.

**Supplementary Figure 15.**
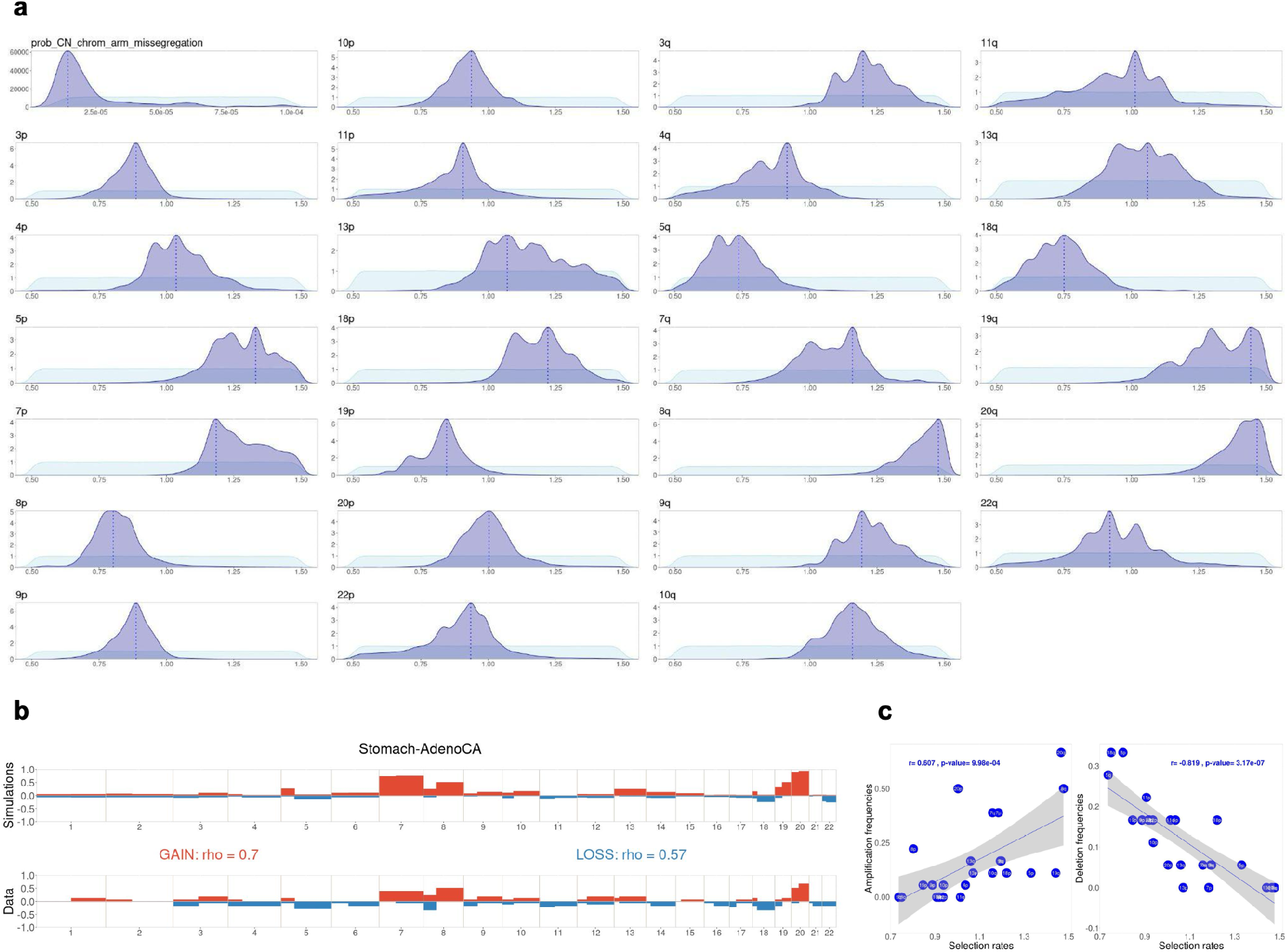
Inference of chromosome-arm selection rates in Stomach-AdenoCA (PCAWG). **(a)** Prior distribution (light blue) and posterior distribution (dark blue) from inference with ABC random forest. Broken line represents the mode in the posterior distribution for each parameter. **(b)** Comparison between simulations with fitted parameter (top) and gain/loss frequencies at arm level from TCGA (bottom). The simulations are computed with the posterior modes from **(a)**. Spearman’s correlation coefficient rho between frequencies of gains (or losses) among each arm in PCAWG and simulations. **(c)** Correlation between inferred selection rates and amplification/deletion frequencies for individual chromosome arms. Linear regressions and p-values from Pearson correlation.

**Supplementary Figure 16.**
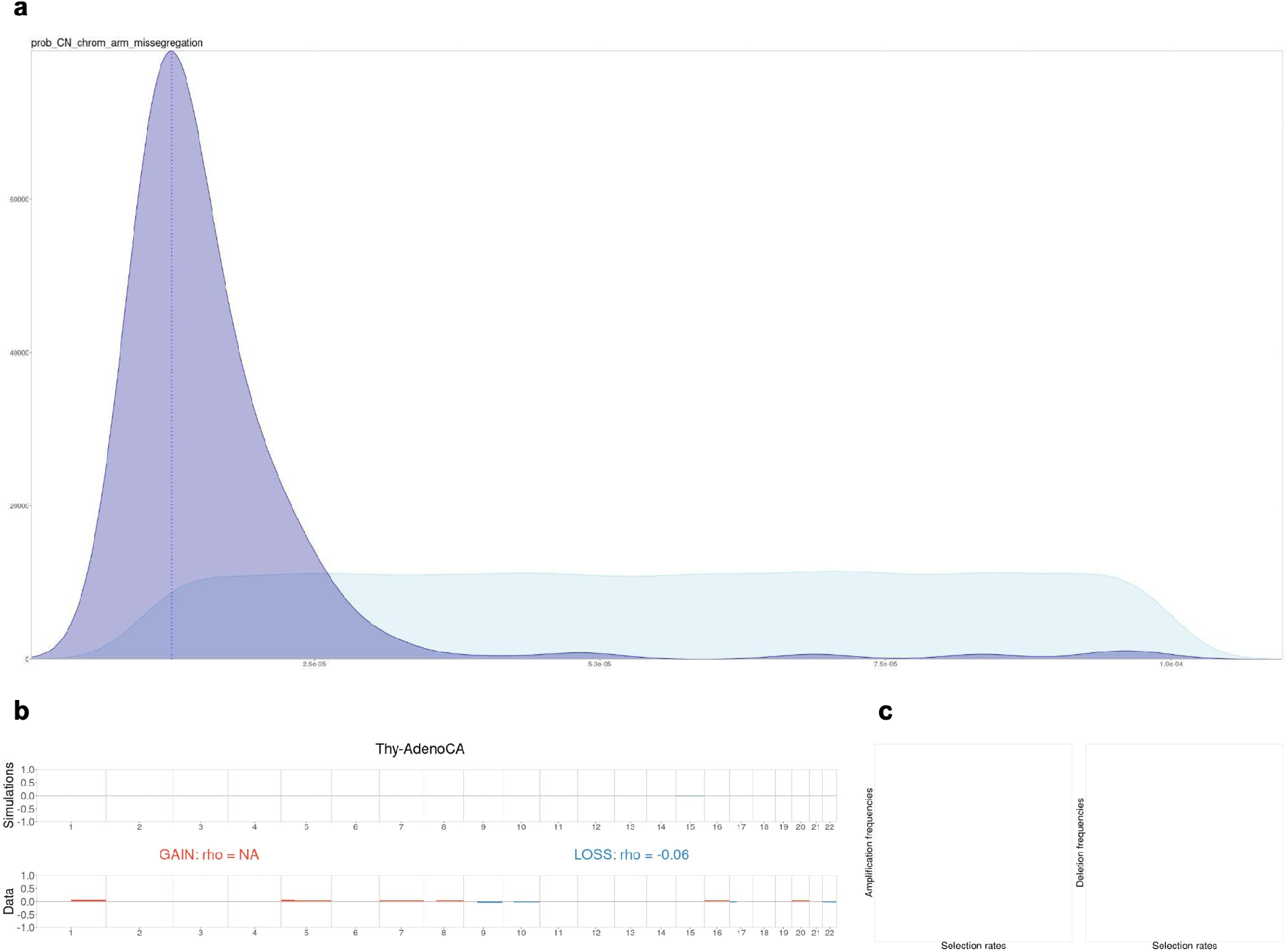
Inference of chromosome-arm selection rates in Thy-AdenoCA (PCAWG). **(a)** Prior distribution (light blue) and posterior distribution (dark blue) from inference with ABC random forest. Broken line represents the mode in the posterior distribution for each parameter. **(b)** Comparison between simulations with fitted parameter (top) and gain/loss frequencies at arm level from TCGA (bottom). The simulations are computed with the posterior modes from **(a)**. Spearman’s correlation coefficient rho between frequencies of gains (or losses) among each arm in PCAWG and simulations. **(c)** Correlation between inferred selection rates and amplification/deletion frequencies for individual chromosome arms. Linear regressions and p-values from Pearson correlation.

**Supplementary Figure 17.**
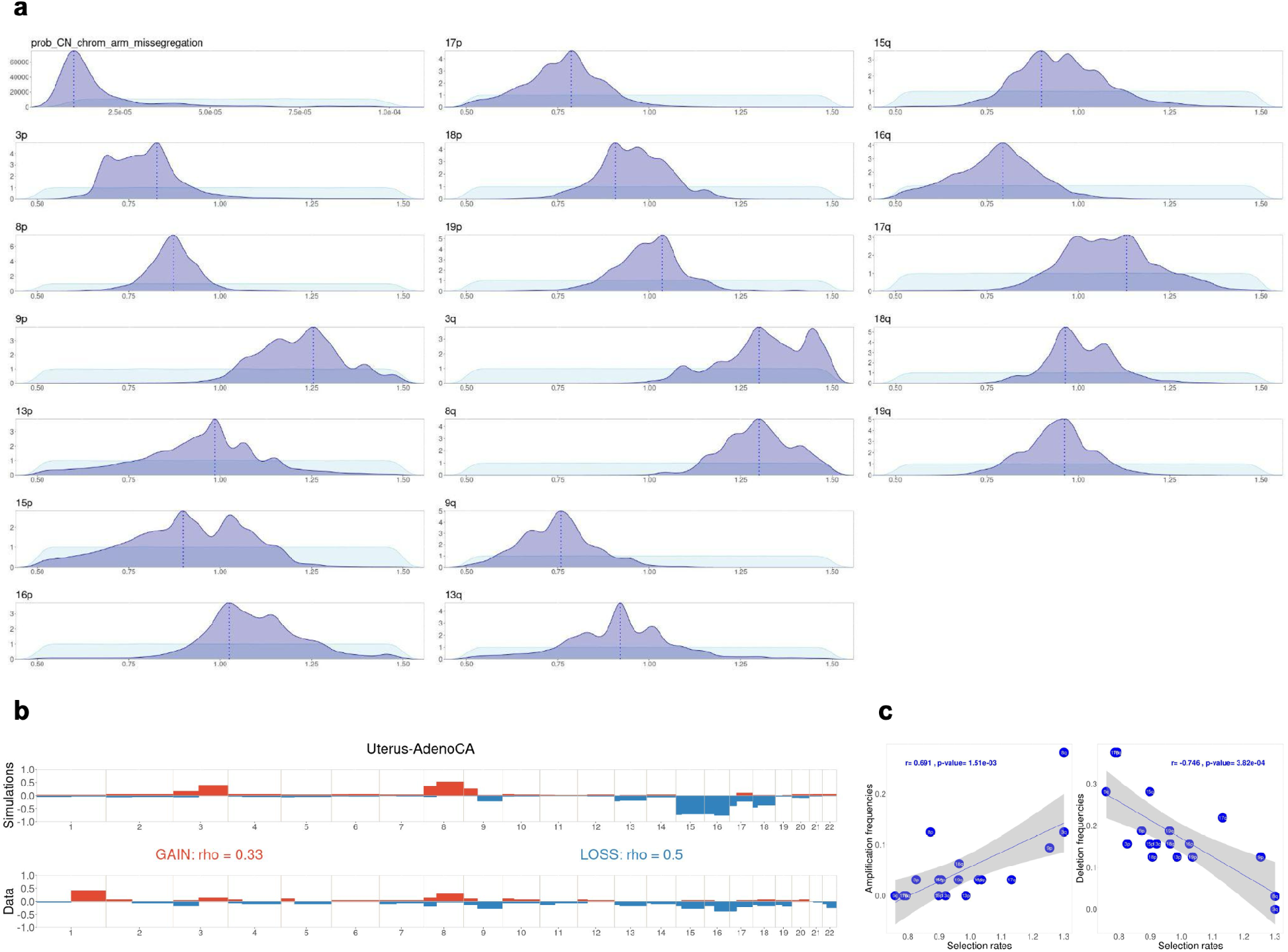
Inference of chromosome-arm selection rates in Uterus-AdenoCA (PCAWG). **(a)** Prior distribution (light blue) and posterior distribution (dark blue) from inference with ABC random forest. Broken line represents the mode in the posterior distribution for each parameter. **(b)** Comparison between simulations with fitted parameter (top) and gain/loss frequencies at arm level from TCGA (bottom). The simulations are computed with the posterior modes from **(a)**. Spearman’s correlation coefficient rho between frequencies of gains (or losses) among each arm in PCAWG and simulations. **(c)** Correlation between inferred selection rates and amplification/deletion frequencies for individual chromosome arms. Linear regressions and p-values from Pearson correlation.

**Supplementary Figure 18.**
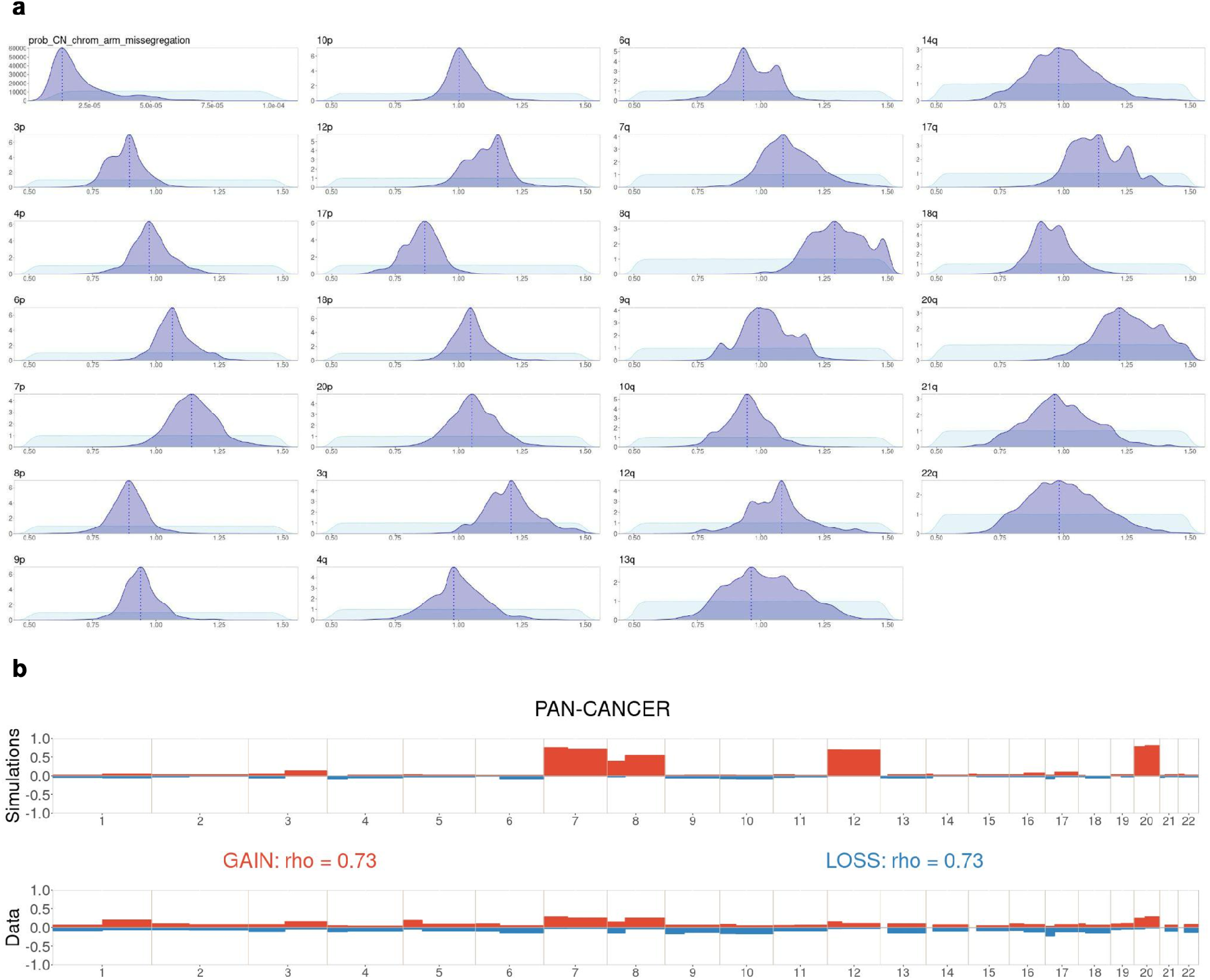
Inference of chromosome-arm selection rates from pan-cancer TCGA. **(a)** Prior distribution (light blue) and posterior distribution (dark blue) from inference with ABC random forest. Broken line represents the mode in the posterior distribution for each parameter. **(b)** Comparison between simulations with fitted parameter (top) and gain/loss frequencies at arm level from TCGA (bottom). The simulations are computed with the posterior modes from **(a)**. Spearman’s correlation coefficient rho between frequencies of gains (or losses) among each arm in TCGA and simulations.

**Supplementary Figure 19.**
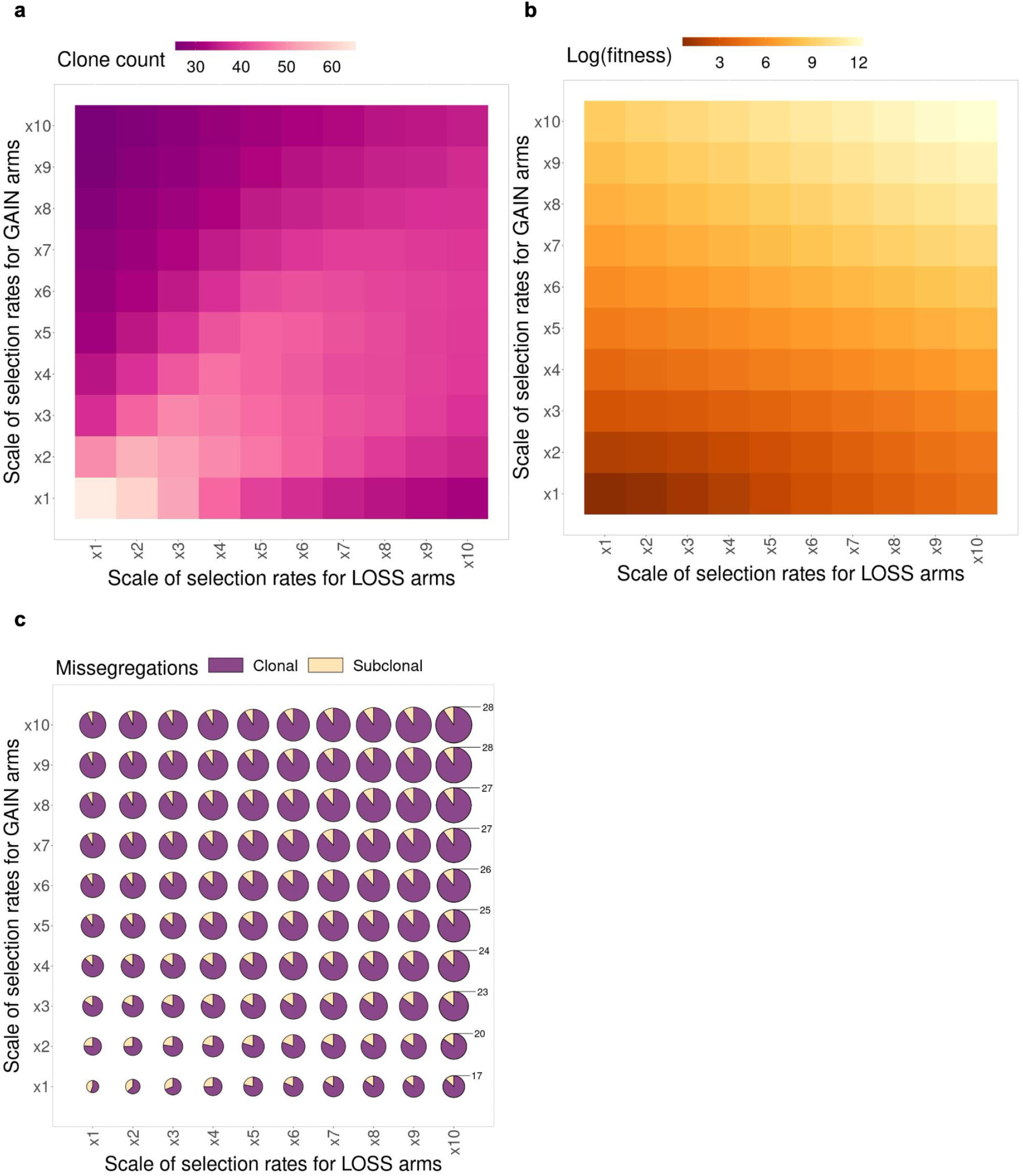
Analysis of selection rates for GAIN and LOSS chromosome arms. Impact of varying parameters on clone count **(a)**, average cell fitness **(b)**, and average count of clonal and subclonal missegregations **(c)** (size of circles indicates the total missegregation counts).

**Supplementary Figure 20.**
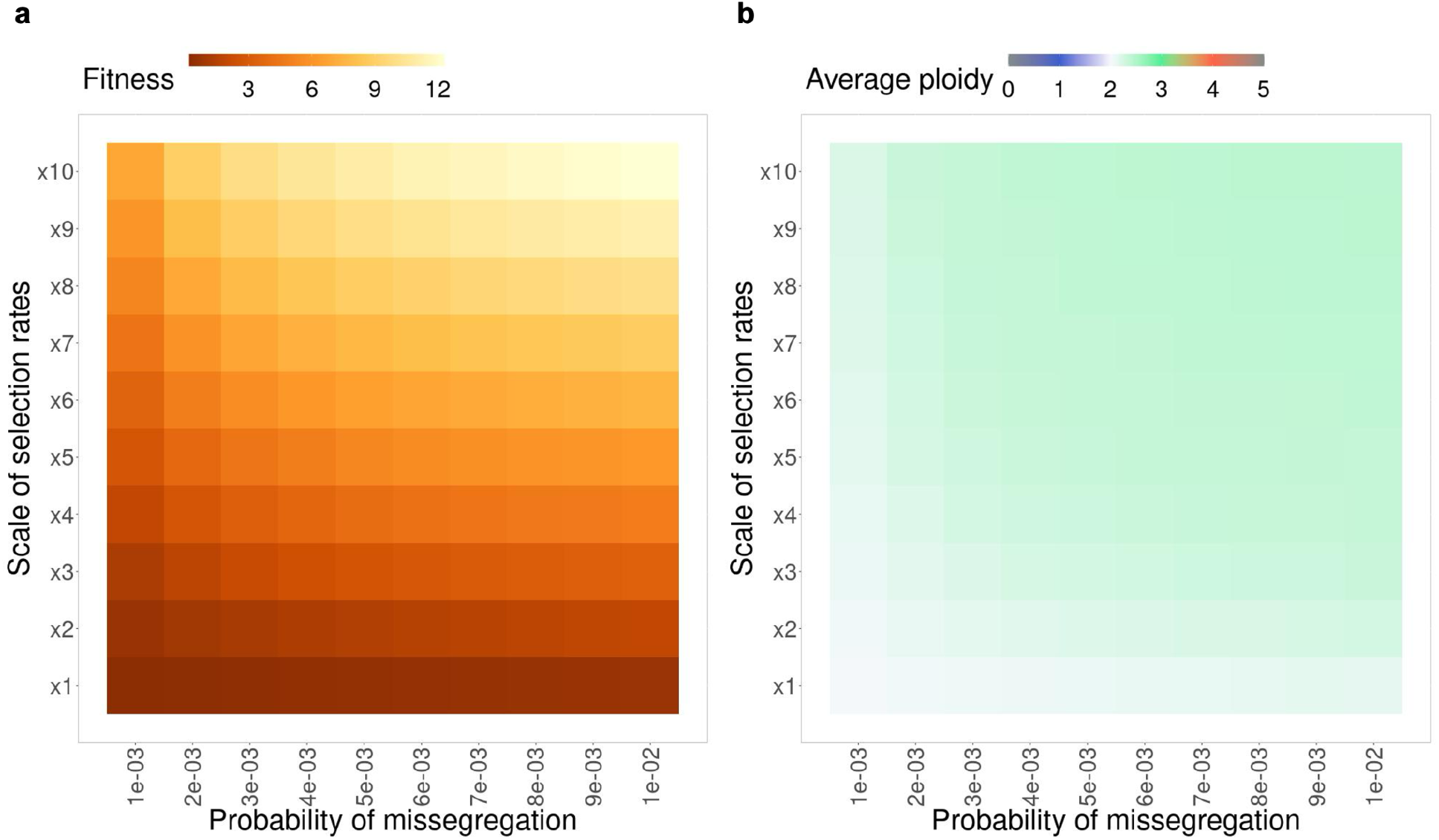
Analysis of probability of missegregation and chromosome-arm selection rates. Impact of varying parameters on average fitness **(a)**, and average ploidy in sample **(b)**.

**Supplementary Figure 21.**
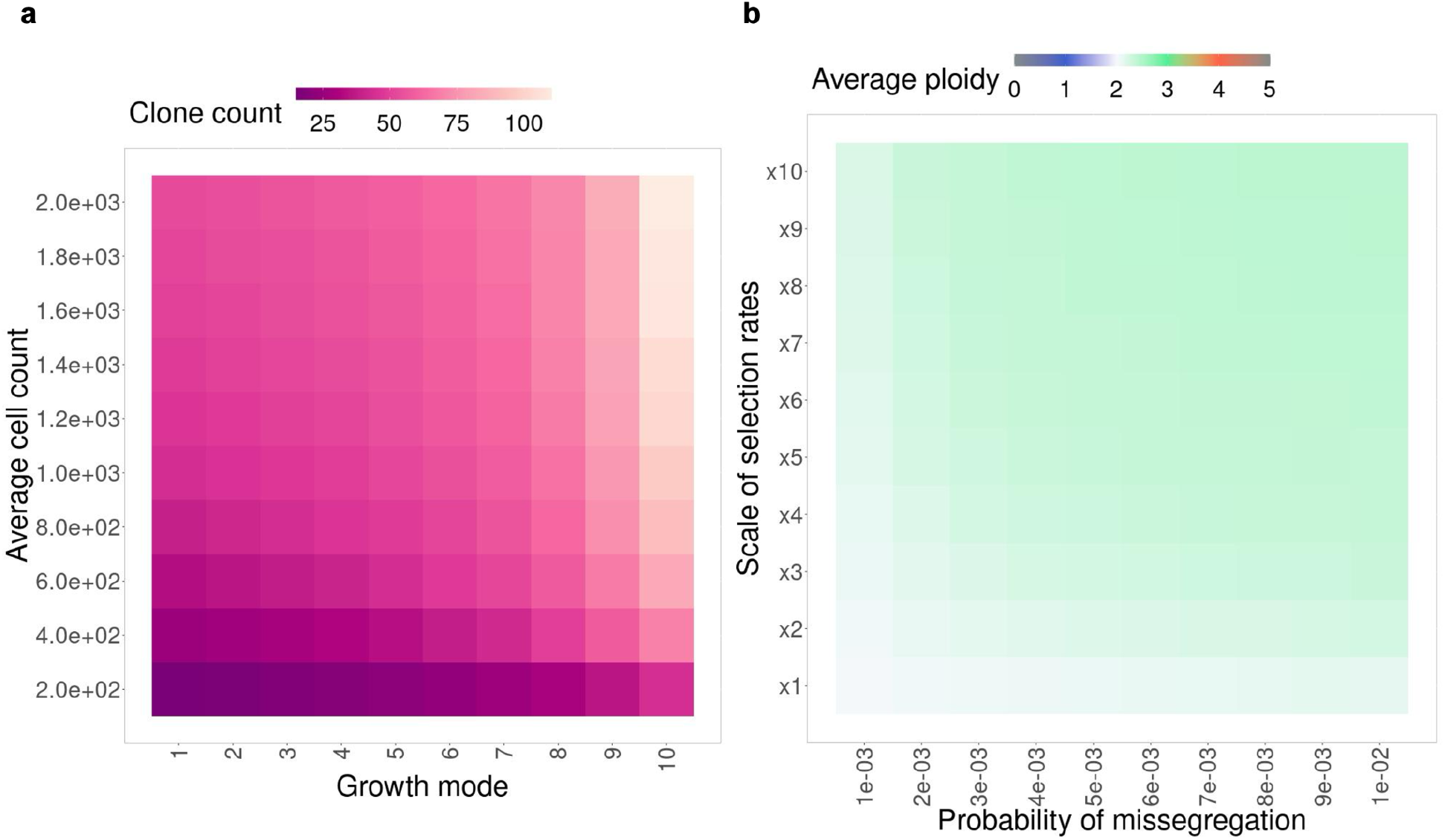
Analysis of growth rate and average cell count. Impact of varying parameters on clone count **(a)**, and average ploidy in sample **(b)**.

**Supplementary Figure 22.**
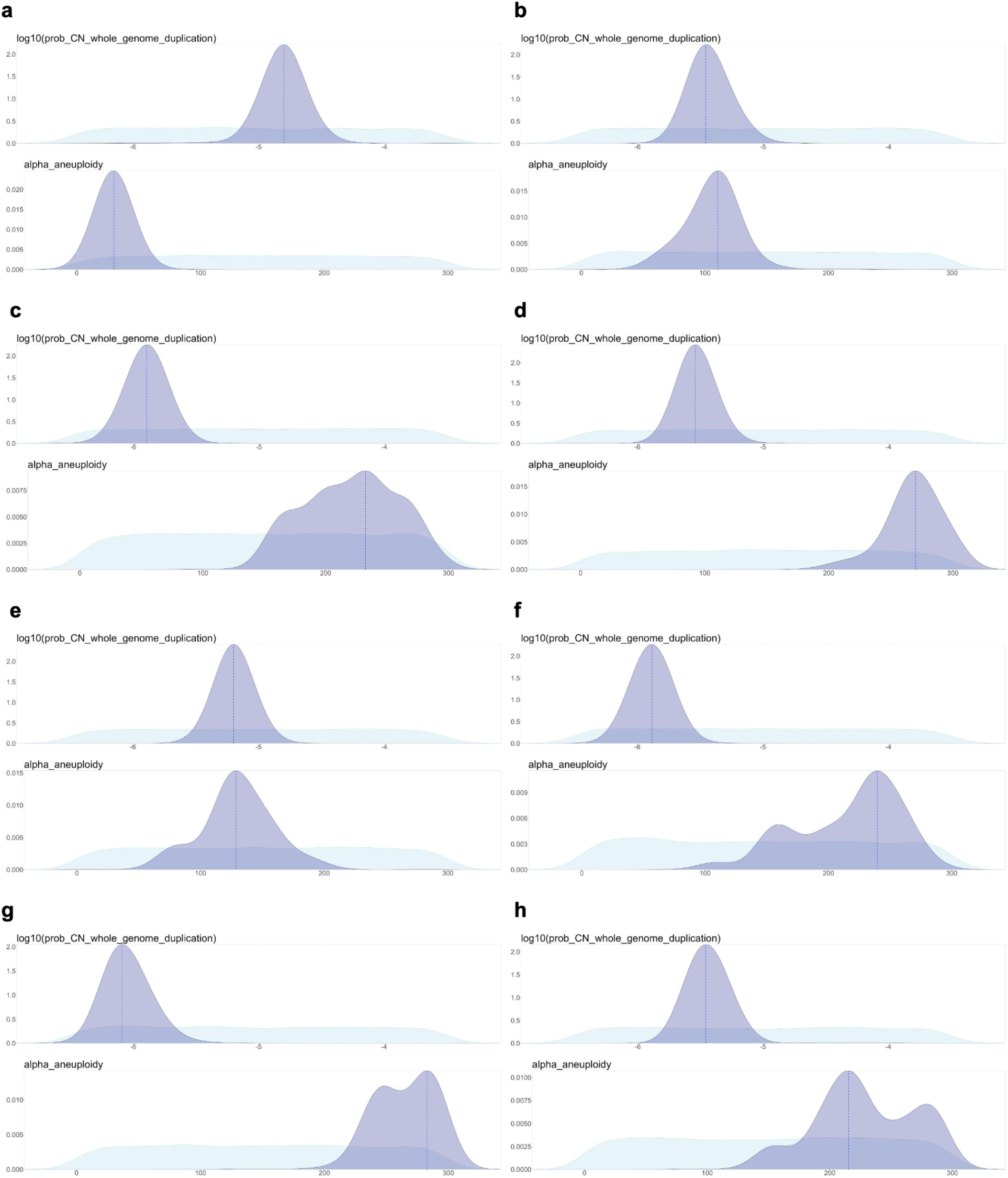
Inference of WGD probability and WGD-aneuploidy rate in individual PCAWG cancer types. Prior distribution (light blue) and posterior distribution (dark blue) from inference with ABC random forest, for Breast-AdenoCA **(a)**, Cervix-SCC **(b)**, CNS-GBM **(c)**, ColoRect-AdenoCA **(d)**, Head-SCC **(e)**, Kidney-ChRCC **(f)**, Kidney-RCC **(g)**, and Liver-HCC **(h)**. Broken line represents the mode in the posterior distribution for each parameter.

**Supplementary Figure 23.**
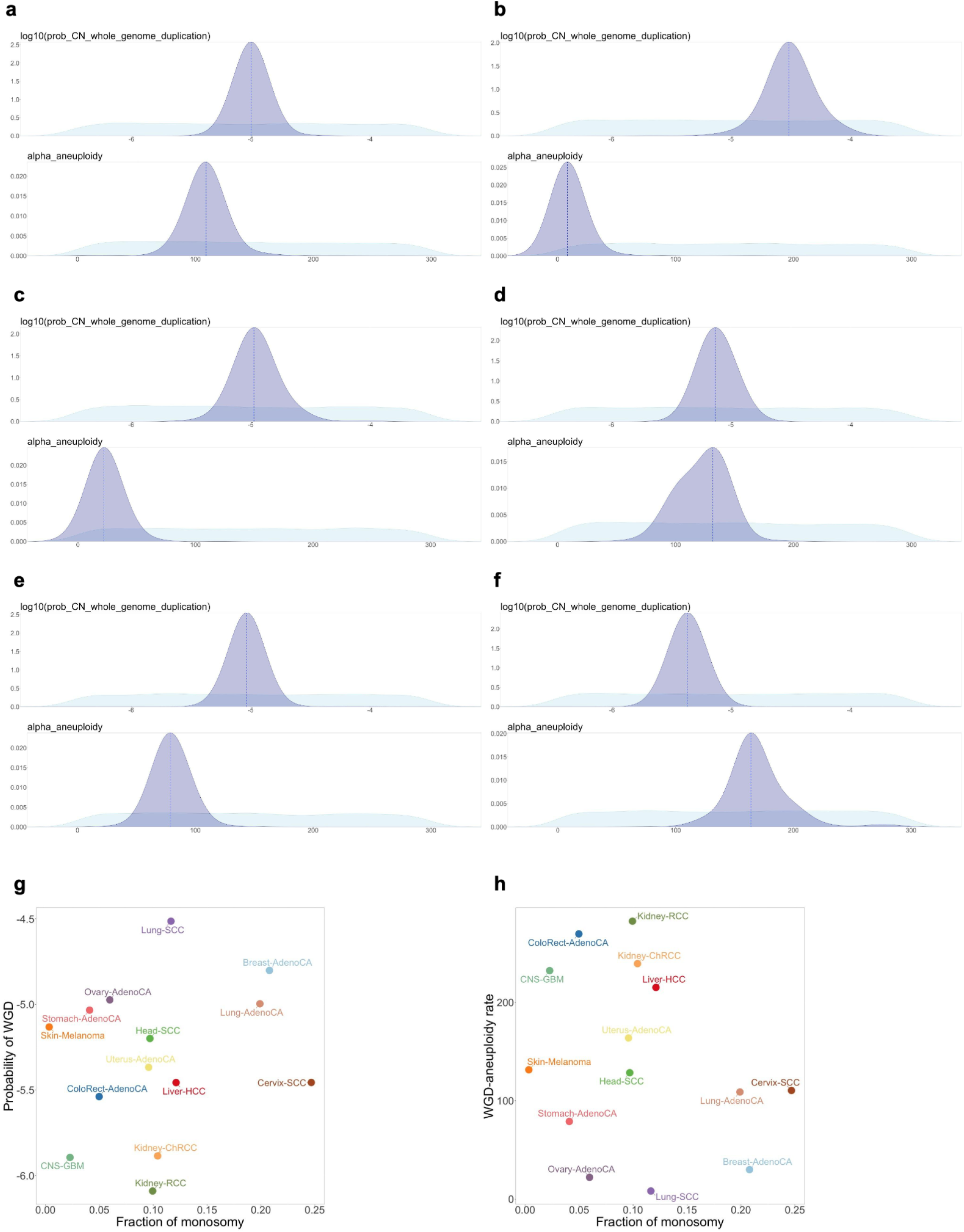
Inference of WGD probability and WGD-aneuploidy rate in individual PCAWG cancer types. **(a-f)** Prior distribution (light blue) and posterior distribution (dark blue) from inference with ABC random forest, for Lung-AdenoCA **(a)**, Lung-SCC **(b)**, Ovary-AdenoCA **(c)**, Skin-Melanoma **(d)**, Stomach-AdenoCA **(e)**, and Uterus-AdenoCA **(f)**. Broken line represents the mode in the posterior distribution for each parameter. **(g-h)** Comparisons between average genomic fraction of monosomy in non-WGD samples and inferred WGD probability **(g)** and WGD-aneuploidy rate **(h)** for each cancer type.

**Supplementary Figure 24.**
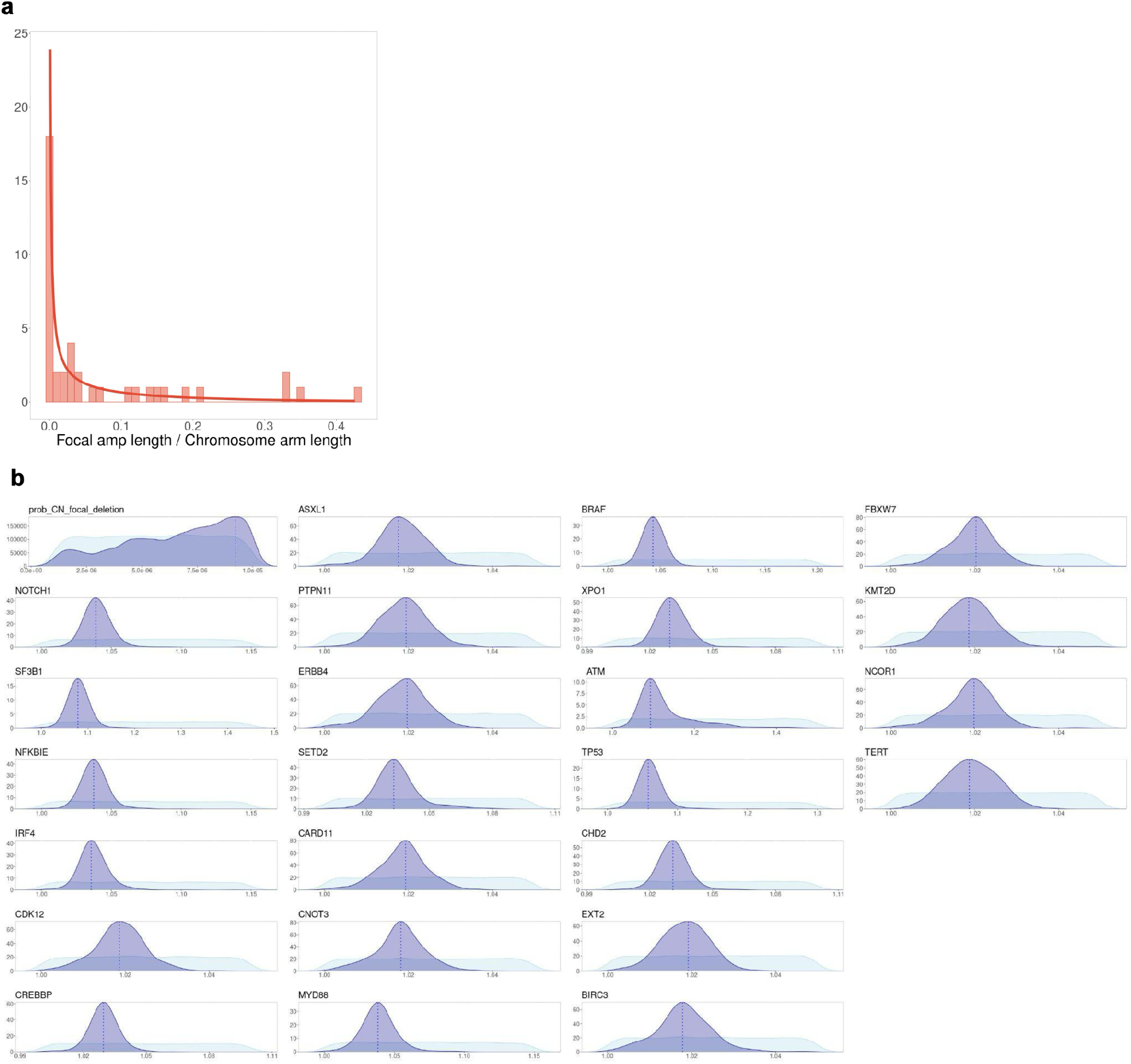
Inference of driver gene selection rates in CLLE-ES (PCAWG). **(a)** Ratios of focal amplification lengths over corresponding chromosome arm lengths are fitted with a Beta distribution. **(b)** Prior distribution (light blue) and posterior distribution (dark blue) from inference with ABC random forest. Broken line represents the mode in the posterior distribution for each parameter.

**Supplementary Note 1**. Mathematical model.

**Supplementary Note 2**. Simulation algorithm.

**Supplementary Note 3**. Inferring missegregation and chromosome-arm selection parameters.

**Supplementary Note 4**. Parameter studies of the chromosome arm selection model.

**Supplementary Note 5**. Inferring WGD parameters.

**Supplementary Note 6**. Inferring driver gene parameters.

