## Supplementary Notes for "CINner: modeling and simulation of chromosomal instability in cancer at single-cell resolution"

April 3, 2024

### 1 Mathematical model

#### 1.1 Characterization of a cell

Because of limitations in machine memory and practical sequencing resolutions, the genome is divided into bins of equal length and we assume each bin in each cell has one allele configuration. Let $\mathcal{N}$ be the chromosome count, $\mathcal{L}$ be the length of each bin and $\mathcal{M}$ be the number of bins spanning the whole genome. Let $\mathcal{M}_i$ be the bin count in chromosome $i$, whose centromere is in bin $\mathcal{C}_i$. We then have $\mathcal{M} = \sum \mathcal{M}_i$ and $\mathcal{L} \cdot \sum \mathcal{M}_i$ is the total length of the genome. For typical DLP+ data, $\mathcal{N} = 23$, $\mathcal{L} = 500,000$ bp and $\mathcal{M} = 6,206$. The copy number (CN) information of a cell is then characterized by:

- $\{J_i : i = 1, \ldots, \mathcal{N}\}$ is the global CN count vector, where $J_i$ is the number of strands of chromosome $i$. $J_i$ changes if the cell gains or loses chromosome strands, e.g. via Whole Genome Duplication or chromosome missegregations.
- $\{K_{i,j} : i = 1, \ldots, \mathcal{N}; j = 1, \ldots, J_i\}$ stores the local CN counts, where $K_{i,j} \in \mathbb{N}^{\mathcal{M}_i}$ is a vector of CN in each bin for strand $j$ of chromosome $i$. The entry $K_{i,j}(l)$ can increase or decrease if the cell undergoes amplifications or deletions affecting the bin $l$.

For a diploid cell in a female, $J_i = 2$ and $K_{i,j} = (1, \ldots, 1), \forall i, j$.

Given the CN set up, every mutation in the cell is assigned an address $(a_1, a_2, a_3, a_4)$, where

- $a_1 \in \{1, \ldots, \mathcal{N}\}$ denotes the chromosome,
- $a_2 \in \{1, \ldots, J_{a_1}\}$ denotes the strand,
- $a_3 \in \{1, \ldots, \mathcal{M}_i\}$ denotes the bin along the strand,
- $a_4 \in \{1, \ldots, K_{a_1,a_2}(a_3)\}$ denotes the CN unit in the bin.

A cell $k$ is then characterized by its CN profile $\left(\left\{J_i^{[k]}\right\}, \left\{K_{i,j}^{[k]}\right\}\right)$ and/or accumulated driver mutations, which define its fitness rate $s_k \in (0, \infty)$. The fitness rate $s_k$ is formulated in sections 1.5 and 1.6.

#### 1.2 Properties of a cell

Cells in the population follow a birth-death process, with two properties defining a given cell $k$:

- The lifespan of the cell is exponentially distributed with rate $\lambda_k$, after which it either divides or dies. We assume $\lambda_k = \lambda$ is constant and equal to the turn-over rate of the cells, estimated from the literature.
- The cell divides with probability $p_k^{\text{div}}$ and dies with probability $1 - p_k^{\text{div}}$. If the cell divides, the daughter cells might have a different CN profile or acquire new driver mutations, according to given probabilities (thereby changing their respective fitness rates). The mechanisms for CN changes and driver mutation acquisition are detailed in sections 1.3 and 1.4.

The division probability for a cell $k$ with fitness rate $s_k$ at time $t$ is

$$p_k^{\text{div}}(t) = g(t) \cdot f(s_k)$$

where $g(t)$ is a negative feedback loop ensuring that the current total cell count $P(t)$ follows the population dynamics $\bar{P}(t)$ observed from either the data or the literature:

$$g(t) = \frac{\bar{P}(t)}{\bar{P}(t) + P(t)}$$

and $f(s_k)$ models the selection for the fittest cells:

$$f(s_k) = \frac{s_k \cdot P(t)}{\sum_{k'=1,\ldots,P(t)} s_{k'}}$$

In the simulation, $\bar{P}(t)$ is interpolated using total population data points $\{t_i, \bar{P}_i\}$.

#### 1.3 Copy number aberration mechanisms

If a cell $k$ with CN profile $\left(\left\{J_i^{[k]}\right\},\left\{K_{i,j}^{[k]}\right\}\right)_{i=1,\dots,\mathcal{N},j=1,\dots,J_i^{[k]}}$ divides into cells $k_1$ and $k_2$, the CN profiles $\left(\left\{J_i^{[k_1]}\right\},\left\{K_{i,j}^{[k_1]}\right\}\right)$ and $\left(\left\{J_i^{[k_2]}\right\},\left\{K_{i,j}^{[k_2]}\right\}\right)$ of $k_1$ or $k_2$ can change if a Copy Number Aberration event (CNA) occurs. We consider the following CNA mechanisms:

- With probability $p_{WGD}$, a Whole Genome Duplication event occurs. In this case, there is only one daughter cell $k_1$, which has double the genomic materials in cell $k$. Therefore,

$$
\begin{aligned}
J_i^{[k_1]} &= 2 \cdot J_i^{[k]} \\
K_{i,j}^{[k_1]} &= K_{i,j}^{[k]} \text{ if } j \leq J_i^{[k]} \\
K_{i,j}^{[k_1]} &= K_{i,j-J_i^{[k]}}^{[k]} \text{ if } j > J_i^{[k]}
\end{aligned}
$$

  A mutation with address $(a_1, a_2, a_3, a_4)$ in cell $k$ translates to two mutations with addresses $(a_1, a_2, a_3, a_4)$ and $\left(a_1, a_2 + J_{a_1}^{[k]}, a_3, a_4\right)$ in cell $k_1$.

- With probability $p_{misseg}$, a whole-chromosome missegregation event occurs. A chromosome $i^*$ is uniformly sampled from $\{1, \dots, \mathcal{N}\}$ conditioned on $J_{i^*}^{[k]} > 0$, a strand $j^*$ is uniformly sampled from $\left\{1, \dots, J_{i^*}^{[k]}\right\}$, and either cell $k_1$ or $k_2$ is chosen to gain the chromosome with equal probabilities. Assuming that cell $k_1$ is chosen to gain, then

$$
\begin{aligned}
J_i^{[k_1]} &= J_i^{[k_2]} = J_i^{[k]} \text{ if } i \neq i^* \\
J_{i^*}^{[k_1]} &= J_{i^*}^{[k]} + 1 \\
J_{i^*}^{[k_2]} &= J_{i^*}^{[k]} - 1 \\
K_{i,j}^{[k_1]} &= K_{i,j}^{[k_2]} = K_{i,j}^{[k]} \text{ if } i \neq i^* \\
K_{i^*,j}^{[k_1]} &= K_{i^*,j}^{[k]} \text{ if } j \leq J_{i^*}^{[k]} \\
K_{i^*,J_{i^*}^{[k]}+1}^{[k_1]} &= K_{i^*,j^*}^{[k]} \\
K_{i^*,j}^{[k_2]} &= K_{i^*,j}^{[k]} \text{ if } j < j^* \\
K_{i^*,j}^{[k_2]} &= K_{i^*,j+1}^{[k]} \text{ if } j^* \leq j \leq J_{i^*}^{[k]} - 1
\end{aligned}
$$

For a mutation with address $(a_1, a_2, a_3, a_4)$ in cell $k$, it will remain the same in cells $k_1$ and $k_2$ if $a_1 \neq i^*$ or $a_2 \neq j^*$. Otherwise, it translates to two mutations with addresses $(a_1, a_2, a_3, a_4)$ and $\left(a_1, J_{i^*}^{[k]} + 1, a_3, a_4\right)$ in cell $k_1$ and is deleted in cell $k_2$.

- With probability $p_{arm-misseg}$, a chromosome-arm missegregation event occurs. We assume this arm exists as an extra chromosome strand in one daughter cell and is lost in the other daughter cell. A chromosome $i^*$ with total bin count $\mathcal{M}_{i^*}$ and centromere in bin $\mathcal{C}_{i^*}$ is uniformly sampled from $\{1, \ldots, \mathcal{N}\}$ conditioned on $J_{i^*}^{[k]} > 0$, a strand $j^*$ is uniformly sampled from $\left\{1, \ldots, J_{i^*}^{[k]}\right\}$, an arm $p$ or $q$ is selected for missegregation with equal probabilities, and either cell $k_1$ or $k_2$ is chosen to gain the chromosome arm with equal probabilities. If arm $p$ is missegregated, then the affected region runs from bin $l_a = 1$ to $l_b = \mathcal{C}_{i^*}$. If arm $q$ is missegregated, then $l_a = \mathcal{C}_{i^*} + 1$ and $l_b = \mathcal{M}_{i^*}$. Assuming that cell $k_1$ is chosen to gain, then

$$
\begin{aligned}
&J_i^{[k_1]} = J_i^{[k_2]} = J_i^{[k]} \text{ if } i \neq i^* \\
&J_{i^*}^{[k_1]} = J_{i^*}^{[k]} + 1 \\
&J_{i^*}^{[k_2]} = J_{i^*}^{[k]} \\
&K_{i,j}^{[k_1]} = K_{i,j}^{[k_2]} = K_{i,j}^{[k]} \text{ if } i \neq i^* \text{ or } j \neq j^* \\
&K_{i^*,j^*}^{[k_1]} = K_{i^*,j^*}^{[k]} \\
&K_{i^*,J_{i^*}^{[k]}+1}^{[k_1]}(l) = K_{i^*,j^*}^{[k]}(l) \text{ if } l_a \leq l \leq l_b \\
&K_{i^*,J_{i^*}^{[k]}+1}^{[k_1]}(l) = 0 \text{ if } l < l_a \text{ or } l > l_b \\
&K_{i^*,j^*}^{[k_2]}(l) = 0 \text{ if } l_a \leq l \leq l_b \\
&K_{i^*,j^*}^{[k_2]}(l) = K_{i^*,j^*}^{[k]}(l) \text{ if } l < l_a \text{ or } l > l_b
\end{aligned}
$$

  A mutation with address $(a_1, a_2, a_3, a_4)$ in cell $k$ remains the same in cells $k_1$ and $k_2$ if $a_1 \neq i^*$ or $a_2 \neq j^*$, or $a_3 < l_a$ or $a_3 > l_b$. Otherwise, it is multiplied to become two mutations with addresses $(a_1, a_2, a_3, a_4)$ and $\left(a_1, J_{a_1}^{[k]} + 1, a_3, a_4\right)$ in cell $k_1$ and is deleted in cell $k_2$.

- With probability $p_{foc-amp}$, a focal amplification event occurs in one of the daughter cells. A region in one arm of a chromosome strand is doubled and the extra genomic material stays on the same strand. A chromosome $i^*$ with total bin count $\mathcal{M}_{i^*}$ and centromere in bin $\mathcal{C}_{i^*}$ is uniformly sampled from $\{1,\ldots,\mathcal{N}\}$, a strand $j^*$ is uniformly sampled from $\left\{1,\ldots,J_{i^*}^{[k]}\right\}$, an arm $p$ (with start bin $\bar{l}_a=1$, end bin $\bar{l}_b=\mathcal{C}_{i^*}$ and length $\mathcal{M}_{arm}=\mathcal{C}_{i^*}$) or $q$ (with start bin $\bar{l}_a=\mathcal{C}_{i^*}+1$, end bin $\bar{l}_b=\mathcal{M}_{i^*}$ and length $\mathcal{M}_{arm}=\mathcal{M}_{i^*}-\mathcal{C}_{i^*}$) is selected for missegregation with equal probabilities, the focal amplification length $\mathcal{M}_{foc-amp}$ is sampled from $\mathcal{M}_{arm}\cdot\text{Beta}\left(\alpha_{foc-amp},\beta_{foc-amp}\right)$, the left border $l_a$ of the amplified region is uniformly sampled from $\left\{\bar{l}_a,\ldots,\mathcal{M}_{arm}-\mathcal{M}_{foc-amp}+1\right\}$, dictating the right border $l_b=l_a+\mathcal{M}_{foc-amp}-1$ of the amplified region, and either cell $k_1$ or $k_2$ is chosen to gain the focal amplification with equal probabilities. Assuming that cell $k_1$ is chosen to gain, then

$$
\begin{aligned}
&J_i^{[k_1]}=J_i^{[k_2]}=J_i^{[k]},\forall i\\
&K_{i,j}^{[k_1]}(l)=K_{i,j}^{[k]}(l)\text{ if }i\neq i^*\text{ or }j\neq j^*\text{ or }l<l_a\text{ or }l>l_b\\
&K_{i^*,j^*}^{[k_1]}(l)=2\cdot K_{i^*,j^*}^{[k]}(l)\text{ if }l_a\leq l\leq l_b\\
&K_{i,j}^{[k_2]}=K_{i,j}^{[k]},\forall i,j
\end{aligned}
$$

All mutations in cell $k$ remain the same in cell $k_2$, and similarly in cell $k_1$ for mutations with addresses $(a_1,a_2,a_3,a_4)$ if $a_1\neq i^*$ or $a_2\neq j^*$, or $a_3<l_a$ or $a_3>l_b$. Otherwise, they double to mutations with addresses $(a_1,a_2,a_3,a_4)$ and $\left(a_1,a_2,a_3,a_4+K_{a_1,a_2}^{[k]}(a_3)\right)$ in cell $k_1$.

- With probability $p_{foc-del}$, a focal deletion event occurs in one of the daughter cells. Chromosome $i^*$, strand $j^*$, and specific arm are sampled similarly to focal amplifications. The focal deletion length is sampled as $\mathcal{M}_{foc-del}\sim\mathcal{M}_{arm}\cdot\text{Beta}\left(\alpha_{foc-del},\beta_{foc-del}\right)$, and the borders $l_a$ and $l_b$ are then sampled accordingly, similar to focal amplification borders. If cell $k_1$ is chosen to gain, then

$$
\begin{aligned}
&J_i^{[k_1]}=J_i^{[k_2]}=J_i^{[k]},\forall i\\
&K_{i,j}^{[k_1]}(l)=K_{i,j}^{[k]}(l)\text{ if }i\neq i^*\text{ or }j\neq j^*\text{ or }l<l_a\text{ or }l>l_b\\
&K_{i^*,j^*}^{[k_1]}(l)=0\text{ if }l_a\leq l\leq l_b\\
&K_{i,j}^{[k_2]}=K_{i,j}^{[k]},\forall i,j
\end{aligned}
$$

All mutations in cell $k$ remain the same in cell $k_2$, and similarly in cell $k_1$ for mutations with addresses $(a_1, a_2, a_3, a_4)$ if $a_1 \neq i^*$ or $a_2 \neq j^*$, or $a_3 < l_a$ or $a_3 > l_b$. Otherwise, they are deleted in cell $k_1$.

#### 1.4 Mutation mechanisms

We assume that a list of $N_{dri-mut}$ driver genes for a particular cancer type is known. For every driver gene, the following information is known: (i) gene ID, (ii) role (either Tumor Suppressor Gene or Oncogene), and (iii) chromosome and bin location (forming $a_1$ and $a_3$ of the gene's address).

For every division of cell $k$ with genomic length $\mathcal{M}^{[k]} = \mathcal{L} \cdot \sum_{i,j} \|K_{i,j}^{[k]}\|_1$, a new driver mutation arises with probability $\mathcal{M}^{[k]} \cdot p_{dri-mut}$ and is inherited by daughter cells $k_1$ and $k_2$. The gene at bin $a_3$ on chromosome $a_1$ being mutated is randomly sampled from all driver genes satisfying that (i) its genomic location is not deleted, i.e. $K_{a_1,a_2}(a_3) \neq 0$ for some strand $a_2$, and (ii) the gene is not already mutated in cell $k$. Then the strand $a_2$ for the gene mutation is sampled from $1, \ldots, J_{a_1}^{[k]}$ (excluding $a_2$ where $K_{a_1,a_2}(a_3) = 0$) and the bin $a_4$ is sampled from $a, \ldots, K_{a_1,a_2}(a_3)$. The driver mutation profiles for cells $k_1$ and $k_2$ are then expanded with driver gene $(a_1, a_2, a_3, a_4)$.

The same process applies for passenger mutations, with rate $p_{pas-mut}$. These mutations are assumed to be neutral, i.e. they do not change a cell's fitness rate. However, upon arrival, they each get assigned a random address and can get multiplied or deleted by the CNA mechanisms in 1.3.

#### 1.5 Selection model

We now discuss the formulae for the fitness rate $s_k$ of a cell $k$. We explore three different models, of selection for chromosome arms, or drive mutations, or both. Each selection model is further subject to viability checkpoints in section 1.6.

##### 1.5.1 Selection model for chromosome arms

In this model, we assume a library of chromosome arms $\{1p, 1q, \ldots\}$, where each chromosome arm $r$ has selection rate $\lambda_r \in (0, \infty)$ and spans from bin $l_a^{[r]}$ to $l_b^{[b]}$. For arm $p$ of chromosome $i$, $l_a^{[r]} = 1$ and $l_b^{[b]} = \mathcal{C}_i$. For arm $q$, $l_a^{[r]} = \mathcal{C}_i + 1$ and $l_b^{[b]} = \mathcal{M}_i$.

Given a cell $k$ with CN profile $\left(\left\{J_i^{[k]}\right\},\left\{K_{i,j}^{[k]}\right\}\right)_{i=1,\ldots,\mathcal{N},j=1,\ldots,J_i^{[k]}}$, we find its ploidy:

$$c^{[k]} = \text{rounded } \frac{\sum_{i,j} K_{i,j}^{[k]}}{\mathcal{M}} \tag{1}$$

and the CN of each arm $r$ on chromosome $i_r$:

$$c_r^{[k]} = \text{rounded } \frac{\sum_{j;l=l_a^{[r]},\ldots,l_b^{[r]}} K_{i_r,j}^{[k]}(l)}{l_b^{[r]} - l_a^{[r]} + 1}$$

The cell's fitness is then

$$s_k = \prod \lambda_r^{c_r^{[k]}/c^{[k]}} \tag{2}$$

The formula 2 ensures that if $\lambda_r > 1$, gaining the arm $r$ gives a cell fitness advantage, while losing the arm makes it less advantageous. The opposite holds if $\lambda_r < 1$.

##### 1.5.2 Selection model for driver mutations

This selection model assumes a library of driver genes for a specific cancer type. Each driver gene $d$ is located at bin $a_3^{[d]}$ on chromosome $a_1^{[d]}$, and has selection rate $\lambda_d \in (0, \infty)$. Based on the gene's specific role in the cancer type, we define its wild-type allele's selection rate $\lambda_{d-WT}$ and mutant allele's selection rate $\lambda_{d-MUT}$.

If driver gene $d$ behaves as an Oncogene (ONC), then

$$\lambda_{d-WT} = \lambda_d$$
$$\lambda_{d-MUT} = \lambda_d^2$$

If driver gene $d$ behaves as a Tumor Suppressor Gene (TSG), then

$$\lambda_{d-WT} = 1/\lambda_d$$
$$\lambda_{d-MUT} = 1$$

Given a cell $k$ with CN profile $\left(\left\{J_i^{[k]}\right\},\left\{K_{i,j}^{[k]}\right\}\right)_{i=1,\ldots,\mathcal{N},j=1,\ldots,J_i^{[k]}}$ and driver mutation profile $\left(n_{d-MUT}^{[k]}; \left\{\left(a_1^{[d]}, a_2^{[z]}, a_3^{[d]}, a_4^{[z]}\right)\right\}_{z=1,\ldots,n_{d-MUT}^{[k]}}\right)_d$, where there are $n_{d-MUT}^{[k]}$ copies of gene $d$ mutants and each copy $z$ is on a different CN unit $a_4^{[z]}$ of strand $a_2^{[z]}$. We define the cell's ploidy with 1 and for each driver gene $d$ is located at bin $a_3^{[d]}$ on chromosome $a_1^{[d]}$, we find the total CN at the bin:

$$n_{d-TOT}^{[k]} = \sum_j K_{a_1^{[d]},j}^{[k]} \left( a_3^{[d]} \right)$$

which leads to the gene's wild-type copy count:

$$n_{d-WT}^{[k]} = n_{d-TOT}^{[k]} - n_{d-MUT}^{[k]}$$

Finally, the cell's fitness is

$$s_k = \prod \lambda_{d-WT}^{2 \cdot n_{d-WT}^{[k]}/c^{[k]}} \cdot \lambda_{d-MUT}^{2 \cdot n_{d-MUT}^{[k]}/c^{[k]}} \tag{3}$$

The fitness of a cell increases if a TSG is mutated or lost (both increase $s_k$ by $\lambda_d$). If an oncogene is mutated or gained, $s_k$ also increases by $\lambda_d$.

##### 1.5.3 Hybrid selection model for chromosome arms and driver mutations

This model combines the selection for chromosome arms in section 1.5.1 and the selection for driver mutations in section 1.5.2. Given the CN and driver mutation profiles of a cell $k$, let $s_k^{[CN]}$ be defined by 2 and $s_k^{[gene]}$ be defined by 3, then the cell's fitness is defined as

$$s_k = s_k^{[CN]} \cdot s_k^{[gene]} \tag{4}$$

#### 1.6 Viability checkpoints

We include conditions for cell viability. If a cell $k$ violates one of these conditions, then $s_k = 0$ and it dies:

- Average ploidy $\frac{\sum K_{i,j}^{[k]}}{\mathcal{M}} \leq \text{ploidy}_{\max}$
- Highest CN in a bin $\max_l \sum K_{i,j}^{[k]}(l) \leq \text{CN}_{\max}$
- Highest normalized CN in a bin $\max_l \frac{\sum K_{i,j}^{[k]}}{c^{[k]}}(l) \leq \text{CN}_{\max}^{\text{nor}}$
- Nullisomy bin count $\sum \mathbf{1}_{l:\sum K_{i,j}^{[k]}(l)=0} \leq \text{nullisomy}_{\max}$
- Count of distinct mutated drivers $\sum \mathbf{1}_{d:n_{d-MUT}>0} \leq \text{driver}_{\max}$
- Count of WGD $\leq \text{WGD}_{\max}$

#### 1.7 Model initialization and sampling

At initial time $t_0$, the simulation starts from $\mathcal{N}_0$ clones. Each clone $n_0$ consists of $N^{[n_0]}$ cells with the CN profile $\left(\left\{J_i^{[n_0]}\right\}, \left\{K_{i,j}^{[n_0]}\right\}\right)$ and driver mutation profile $\left(n_{d-MUT}^{[n_0]}; \left\{\left(a_1^{[d]}, a_2^{[z]}, a_3^{[d]}, a_4^{[z]}\right)\right\}_{z=1,\ldots,n_{d-MUT}^{n_0}}\right)$.

The cell population evolves according to the birth-death process described above, until time $t_f$. A sample of $N_{\text{sample}}$ cells is drawn from the total population at time $t_{\text{sample}}$, and the aim for the algorithm is to simulate characteristics of this cell sample.

### 2 Simulation algorithm

In this section, we detail the simulation algorithm. Several observations are employed to optimize the runtime for each simulation, which is necessary for either large cell populations or high number of simulations, both of which are usually the case in parameter fitting. The first observation is that cells having the same phylogenetic origin have the same CN and mutational profiles, therefore they have the same fitness rate and behave similarly. Secondly, the only information that is relevant to oncologists is restricted to within the sequenced sample, which is magnitudes smaller in cell count compared to the whole tumor. Therefore, instead of simulating individual cells, we define a clone to consist of cells with the same phylogenetic origin, and simulate the clones' populations first (section 2.2). Note that in the case of parallel or convergent evolution, it is possible for different clones to have the same profile, but the simulator would have records of the distinct clonal arrivals. Next, the algorithm samples cells to be sequenced (section 2.3) and simulates the phylogeny of these cells based on the clonal population changes (section 2.4). The simulation algorithm decides which step to stop at, based on whether further steps are necessary to produce the data requested by the user.

#### 2.1 Simulation initialization

The necessary variables for setting up the simulations are listed in Table 1.

Table 1: Input variables for simulations

| Variable | Definition | Reference |
|---|---|---|

| | | |
|---|---|---|
| $t_0$ | Simulation start time | 1.7 |
| $t_f$ | Simulation end time | 1.7 |
| $\{t_i, \bar{P}_i\}$ | Total population dynamics | 1.2 |
| $t_{\text{sample}}$ | Time of sample | 1.7 |
| $N_{\text{sample}}$ | Cell count of sample | 1.7 |
| $\lambda$ | Cell's turn-over rate | 1.2 |
| $\tau$ | Time step in clonal evolution | 2.2 |
| $\mathcal{L}$ | Bin length in bp | 1.1 |
| $p_{WGD}$ | Prob. of WGD | 1.3 |
| $p_{misseg}$ | Prob. of missegregation | 1.3 |
| $p_{arm-misseg}$ | Prob. of arm missegregation | 1.3 |
| $p_{foc-amp}$ | Prob. of focal amplification | 1.3 |
| $\alpha_{foc-amp}, \beta_{foc-amp}$ | Beta parameters of focal amplification | 1.3 |
| $p_{foc-del}$ | Prob. of focal deletion | 1.3 |
| $\alpha_{foc-del}, \beta_{foc-del}$ | Beta parameters of focal deletion | 1.3 |
| $p^{neu}_{WGD}$ | Prob. of neutral WGD | 2.5 |
| $p^{neu}_{misseg}$ | Prob. of neutral missegregation | 2.5 |
| $p^{neu}_{arm-misseg}$ | Prob. of neutral arm missegregation | 2.5 |
| $p^{neu}_{foc-amp}$ | Prob. of neutral focal amplification | 2.5 |
| $\alpha^{neu}_{foc-amp}, \beta^{neu}_{foc-amp}$ | Beta parameters of neutral focal amplification | 2.5 |
| $p^{neu}_{foc-del}$ | Prob. of neutral focal deletion | 2.5 |
| $\alpha^{neu}_{foc-del}, \beta^{neu}_{foc-del}$ | Beta parameters of neutral focal deletion | 2.5 |
| $p_{dri-mut}$ | Prob. of driver mutations per bp | 1.4 |
| $p_{pas-mut}$ | Prob. of passenger mutations per bp | 1.4 |
| $\{\mathcal{M}_i\}$ | Bin count of each chromosome | 1.1 |
| $\{\mathcal{C}_i\}$ | Bin location of each centromere | 1.1 |
| $\text{ploidy}_{\max}$ | Max ploidy in viable cell | 1.6 |
| $\text{CN}_{\max}$ | Max bin CN in viable cell | 1.6 |
| $\text{CN}^{\text{nor}}_{\max}$ | Max bin normalized CN in viable cell | 1.6 |
| $\text{WGD}_{\max}$ | Max WGD count in viable cell's history | 1.6 |
| $\text{nullisomy}_{\max}$ | Max nullisomy bin count in viable cell | 1.6 |
| $\text{driver}_{\max}$ | Max distinct driver count in viable cell | 1.6 |
| $\{\lambda_r\}$ | Selection rate of each arm | 1.5.1 |
| $\{\lambda_d\}$ | Selection rate of each driver gene | 1.5.2 |
| $\{N^{[n_0]}\}$ | Clonal cell counts at $t_0$ | 1.7 |
| $\left(\left\{J^{[n_0]}_i\right\}, \left\{K^{[n_0]}_{i,j}\right\}\right)$ | Clonal CN profiles at $t_0$ | 1.7 |
| $\left(n^{[n_0]}_{d-MUT}, \left\{\left(a^{[d]}_1, a^{[z]}_2, a^{[d]}_3, a^{[z]}_4\right)\right\}\right)$ | Clonal mutation profiles at $t_0$ | 1.7 |

#### 2.2 Clonal evolution

For convenience, let $\{p_q\}_{q=1,\ldots,\mathcal{Q}}$ be the positive probabilities of mechanisms ending in either new CN or mutational profiles. This includes $p_{dri-mut}$, $p_{WGD}$, $p_{misseg}, \cdots$

Assume that at time point $t$, the population consists of $\mathcal{N}_t$ clones. There are $N^{[n]}$ cells of each clone $n$, each with division probability $p^{div}_n(t)$. Since all cells have the same turn-over rate of $\lambda$, the next event in the population takes place at time $t + \tau_t$, where

$$\tau_t \sim \text{Exponential}\left(\lambda \cdot \sum_{n=1,\ldots,\mathcal{N}_t} N^{[n]}\right)$$

The cell $k$ to undergo the event is chosen uniformly from the $\sum N^{[n]}$ cells. If this cell belongs to clone $n$, it divides with probability $p_n^{div}(t)$ or dies with probability $1 - p_n^{div}(t)$. If division is chosen, each mechanism $q$ is sampled with probability $p_q$ to either keep the daughter cells in clone $n$, or create new clones for them, with updated profiles following directions in sections 1.3 and 1.4.

This Gillespie algorithm [3] is statistically exact. However, as total population $\sum N^{[n]}$ increases, the time step $\tau_t$ decreases and each simulation takes increasingly more steps to finish. The runtime for $\sum N^{[n]} \approx 10^6 - 10^9$ cells, as the case for most cancer types, becomes untenable for practical purposes.

We therefore employ the tau-leaping method [1], an approximation of the Gillespie algorithm. At time point $t$, we dictate the next time step to be $\tau$, then simulate the event counts in each clone. The total event count in clone $n$ within time step $[t, t + \tau)$ is

$$e_n \sim \text{Poisson}\left(\tau \cdot \lambda \cdot N^{[n]}\right) \tag{5}$$

Note that if the rate is too large (e.g., $\tau \cdot \lambda \cdot N^{[n]} > 1000$), Poisson random number generator can be time-consuming, in which case we replace 5 with

$$e_n \sim \text{Normal}\left(\tau \cdot \lambda \cdot N^{[n]}, \sqrt{\tau \cdot \lambda \cdot N^{[n]}}\right)$$

In either case, the total event count is conditioned to $e_n \leq N^{[n]}$, as the number of events cannot exceed the number of cells. The $e_n$ events are divided into cell divisions $e_b$ and cell deaths $e_d$:

$$(e_b, e_d) \sim \text{Multinomial}\left(e_n; p_b = p_n^{div}(t), p_d = 1 - p_n^{div}(t)\right)$$

and the cell divisions are further divided into those resulting in new profiles ($e_{b-new}$) and those remaining in clone $n$ ($e_{b-old}$):

$$(e_{b-new}, e_{b-old}) \sim \text{Multinomial}\left(e_b; p_{b-new} = p_{new}, p_d = 1 - p_{new}\right)$$

where $p_{new} = 1 - \prod_{q=1,\ldots,\mathcal{Q}}(1 - p_q)$ is the probability that any given cell division leads to creation of new cell profiles and therefore new clones. Once $(e_{b-new}, e_{b-old}, e_d)$ is known, the clones are updated as

$$N^{[n]} \leftarrow N^{[n]} - e_d - e_{b-new} + e_{b-old}$$
$$\text{Clones } n' = \mathcal{N}_t + 1, \ldots, \mathcal{N}_t + e_{b-new} \text{ are initialized}$$
$$\mathcal{N}_t \leftarrow \mathcal{N}_t + e_{b-new}$$

The most consequential variable in this step is the choice of $\tau$. The tau-leaping algorithm converges to the exact Gillespie algorithm as $\tau \to 0$, but the simulation runtime increases accordingly. Large $\tau$ leads to fast simulations but the results might diverge from statistical expectation, and drawing $e_n$ with 5 conditioned to $e_n \leq N^{[n]}$ could take many tries. In practice, we found $\tau = \frac{\lambda}{2}$ to be a good trade-off.

The clonal death counts are not utilized in later steps, as dead cells cannot be ancestral nodes for cells at a later time. Therefore, the storage requirement for events in all clones at time point $t$ is restricted to

$$\left\{ \left( w^{[r]}, g_0^{[r]}, g_1^{[r]}, g_2^{[r]} \right) : r = 1 \ldots, \mathcal{R}_t \right\} \tag{6}$$

where there are $\mathcal{R}_t$ distinct division groups. Each group $r$ consists of $w^{[r]}$ cells of clone $g_0^{[r]}$, each of which divides into one cell of clone $g_1^{[r]}$ and one cell of clone $g_2^{[r]}$. For instance, the cell divisions restricted within clone $n$ would constitute group $r$ with $w^{[r]} = e_{b-old}$, and $g_0^{[r]} = g_1^{[r]} = g_2^{[r]} = n$. Clone $n$ is furthermore associated with $e_{b-new}$ other groups, where $w^{[r]} = 1$, $g_0^{[r]} = n$ and $g_1^{[r]}$ and $g_2^{[r]}$ denote the daughter clones arriving as results to CNA or mutational mechanisms.

#### 2.3 Sampling

After 2.2, at any given time $t$, we have the total clone count $\mathcal{N}_t$ and cell count $N^{[n]}$ of each clone $n$. The sample of $N_{\text{sample}}$ cells at time $t_{\text{sample}}$ is uniformly drawn from the $\sum_{n=1,\ldots,\mathcal{N}_{\text{sample}}} N^{[n]}$ cells present. We then have $\left\{ N_{\text{sample}}^{[n]} \right\}$, where $N_{\text{sample}}^{[n]}$ is the cell count of clone $n$ in the sample.

#### 2.4 Sampled cell phylogeny

The goal of this step is to build the phylogeny for the $N_{\text{sample}}$ cells found in section 2.3, using information stored in section 2.2. The sampled cell phylogeny is initialized at time $t_{\text{sample}}$ as containing $\sum N^{[n]}_{\text{sample}}$ nodes, each with known clonal index.

At any time $t+\tau$, the phylogeny is at state

$$\left\{\left(N^{[n,t+\tau]}, N^{[n,t+\tau]}_{\text{nodes}}\right)\right\}_{n=1,\ldots,\mathcal{N}_{t+\tau}}$$

where $\mathcal{N}_{t+\tau}$ is the number of clones present in the total population, clone $n$ has $N^{[n,t+\tau]}$ cells in the total population and is represented by $N^{[n,t+\tau]}_{\text{nodes}}$ nodes in the phylogeny. From 6, there are $\sum_r w^{[r]}$ cell divisions in the total population within $[t, t+\tau)$, and each division of any group $r$ can be indexed as

$$\left(i^{[d]}_0\right)_{g^{[d]}_0} \to \left(i^{[d]}_1\right)_{g^{[d]}_1} + \left(i^{[d]}_2\right)_{g^{[d]}_2} : d = 1, \ldots, \sum_r w^{[r]} \tag{7}$$

where a cell in clone $g^{[d]}_0$ divides into a cell of clone $g^{[d]}_1$ and a cell of clone $g^{[d]}_2$. If the former daughter cell is a node then $i^{[d]}_1$ denotes its node index, otherwise $i^{[d]}_1 = \emptyset$, and similarly for $i^{[d]}_2$. Updating the phylogeny to time $t$ requires first simulating $\left\{i^{[d]}_1\right\}$ and $\left\{i^{[d]}_2\right\}$.

For a given clone $n$, let

$$\begin{aligned} h^{[n]} &= \sum \mathbf{1}_{i^{[d]}_y : y\in\{1,2\}, g^{[d]}_y = n} \\ \widetilde{h}^{[n]} &= \sum \mathbf{1}_{i^{[d]}_y : y\in\{1,2\}, g^{[d]}_y = n, i^{[d]}_y \neq \emptyset} \end{aligned}$$

In other words, $h^{[n]}$ is the count of freshly divided cells in clone $n$ at time $t+\tau$ within the total population, and $\widetilde{h}^{[n]}$ is the count of those $h^{[n]}$ cells that are in the current node list. We then have

$$\widetilde{h}^{[n]} \sim \text{Binomial}\left(h^{[n]}, p = \frac{N^{[n,t+\tau]}_{\text{nodes}}}{N^{[n,t+\tau]}}\right)$$

conditioned with $\widetilde{h}^{[n]} \leq N^{[n,t+\tau]}_{\text{nodes}}$. A random sample of size $\widetilde{h}^{[n]}$ among the cells $i^{[d]}_y$ in 7 satisfying $g^{[d]}_y = n$ and $y \in \{1,2\}$ are then assigned node indices drawn randomly from $\left\{1, \ldots, N^{[n,t+\tau]}_{\text{nodes}}\right\}$ without replacement.

After all the daughter cell indices are simulated, the phylogeny is updated by going through the cell divisions $d$ in 7:

- If $i_1^{[d]} = \emptyset$ and $i_2^{[d]} = \emptyset$: This means both daughter cells of the division $d$ are not part of the current node list, therefore the cell phylogeny is not updated.

- If either $i_1^{[d]} \neq \emptyset$ or $i_2^{[d]} \neq \emptyset$: Without loss of generality, assume $i_1^{[d]} \neq \emptyset$. The node $i_1^{[d]}$ then contains one more cell generation and is updated to clone $g_0^{[d]}$.

- If both $i_1^{[d]} \neq \emptyset$ and $i_2^{[d]} \neq \emptyset$: The nodes $i_1^{[d]}$ and $i_2^{[d]}$ then merge into one node indexed $i_0^{[d]}$ containing one cell generation of clone $g_0^{[d]}$.

This process runs backward in time until $t = t_0$ and the phylogeny for the $N_{\text{sample}}$ sampled cells is finished.

#### 2.5 Neutral Copy Number Aberrations

By this step, the algorithm has a simulated cell phylogeny, tracing the ancestry of sampled cells at $t_{\text{sample}}$ back to initial time. Each edge in the phylogeny can harbor events (CNAs or driver mutations), as described in sections 1.3 and 1.4, each of which molded the selection landscape by changing affected cells' fitness, as formulated in section 1.5, and consequently led to the expansion of the edge. Nodes downstream of a given edge belong to the same clone and therefore have the same CN profile, unless another event occurs within them. However, single-cell DNA-seq data commonly features a lot of cell-to-cell variation among each clone. This step of the algorithm attempts to simulate these variations, assuming that they are neutral, i.e. they do not affect the cell's fitness rate in comparison to the clone's.

Going down the phylogeny tree from $t = t_0$, for each cell generation in each edge, we sample the counts of neutral CNAs similarly as in section 1.3, except with probability $p_{WGD}^{neu}$, $p_{misseg}^{neu}$, $p_{arm-misseg}^{neu}, \ldots$ instead of $p_{WGD}$, $p_{misseg}$, $p_{arm-misseg}, \ldots$ The CN profile of the edge is updated accordingly, with two restraints. First, we require the neutral CNAs to not affect the selective CNAs downstream, i.e. the chromosome strands, arms, and bin locations chosen for neutral CNAs are conditioned to exclude those known to change in any progeny of the edge. Second, even though neutral CNAs do not affect the cells' fitness, the CN profile resulting from them cannot violate the viability checkpoints in section 1.6. If neither of these requirements can be satisfied in an edge, then it cannot harbor any neutral CNAs.

### 3 Inferring missegregation and chromosome-arm selection parameters

#### 3.1 Parameter fitting scheme

Assume that we have a library of $\mathcal{S}$ samples undergoing bulk DNA-sequencing, resulting in $\left\{f_r^{[\text{gain}]}, f_r^{[\text{loss}]}\right\}$, where $f_r^{[\text{gain}]}$ is the frequency of gains of chromosome arm $r$ and $f_r^{[\text{loss}]}$ is the frequency of its losses across the $\mathcal{S}$ samples. A chromosome arm is considered gained or lost if its CN is higher or lower than the cell's average ploidy. Here we present a scheme to fit the chromosome arm selection model (Section 1.5.1).

Three CNA mechanisms described in section 1.3 affect entire chromosome arms: Whole Genome Duplications, whole-chromosome missegregations and chromosome-arm missegregations. However, Whole Genome Duplication doubles all chromosome strands, therefore it does not change the gain/loss frequency of chromosome arms in a given cell. Therefore, only probability $p_{misseg}$ of whole-chromosome missegregation and probability $p_{arm-misseg}$ of chromosome-arm missegregation during a cell division are relevant to the parameter fitting routine. The target for fitting is therefore $p_{misseg}$, $p_{arm-misseg}$ and chromosome arm selection rates $\{\lambda_r : r = 1\text{p}, 1\text{q}, \dots\}$.

The fitting routine employs `abcrf` [4], an Approximate Bayesian Computation method relying on the random forest methodology. The first step of `abcrf` requires a set of reference data. We sample $p_{misseg}$, $p_{arm-misseg}$ and $\{\lambda_r\}$ from their respective prior distributions, then create $\mathcal{S}$ simulations. Each simulation is characterized by the CN profile of the clone with the highest population, assumed to be equivalent to the result of bulk DNA-seq. The gain/loss frequencies from the simulations then form the statistics for the corresponding parameter set. In the second step, `abcrf` is trained on the reference data for each parameter, then predicts the posterior distribution for the parameter based on $\left\{f_r^{[\text{gain}]}, f_r^{[\text{loss}]}\right\}$. Finally, we choose one parameter set $\hat{p}_{misseg}$, $\hat{p}_{arm-misseg}$ and $\left\{\hat{\lambda}_r\right\}$ to compare against the data, where each parameter is chosen from the mode of their posterior distribution. Algorithm 1 outlines the fitting routine.

We experimented with two variations of Algorithm 1: (i) both $p_{misseg}$ and $p_{arm-misseg}$ vary, and (ii) $p_{arm-misseg}$ varies while $p_{misseg}$ is fixed. The second scheme leads to better results. This is because the gain/loss frequencies can

**Algorithm 1** Parameter fitting for the chromosome arm selection model (section 1.5.1)

**function** FREQUENCY$(p_{misseg}, p_{arm-misseg}, \{\lambda_r\})$
  **for** $j \leftarrow 1, \ldots, \mathcal{S}$ **do**
    Create one simulation
    Clonal CN profile $\leftarrow$ CN profile of most populous clone
  **end for**
  Compute $\left\{f_r^{[\text{gain;i}]}, f_r^{[\text{loss;i}]}\right\}$ from the clonal CN profiles
**end function**

**for** $i \leftarrow 1, \ldots, N_{\text{simulations}}$ **do** ▷ Create reference data
  $p_{misseg}^{[i]}, p_{arm-misseg}^{[i]}, \left\{\lambda_r^{[i]}\right\} \sim$ prior distributions
  $\left\{f_r^{[\text{gain;i}]}, f_r^{[\text{loss;i}]}\right\} \leftarrow$ FREQUENCY $\left(p_{misseg}^{[i]}, p_{arm-misseg}^{[i]}, \left\{\lambda_r^{[i]}\right\}\right)$
**end for**
**for** $p \leftarrow p_{misseg}, p_{arm-misseg}, \lambda_{1\text{p}}, \lambda_{1\text{q}}, \ldots$ **do** ▷ ABC random forest
  Train `abcrf` on $\left\{p^{[i]}; f_r^{[\text{gain;i}]}, f_r^{[\text{loss;i}]}\right\}$
  Posterior distribution for $p \sim$ `abcrf` prediction for $\left\{f_r^{[\text{gain}]}, f_r^{[\text{loss}]}\right\}$
  $\hat{p}$ = mode of posterior distribution
**end for**
$\left\{\hat{f}_r^{[\text{gain}]}, \hat{f}_r^{[\text{loss}]}\right\} \sim$ FREQUENCY $\left(\hat{p}_{misseg}, \hat{p}_{arm-misseg}, \left\{\hat{\lambda}_r\right\}\right)$ ▷ Validation
Compare $\left\{\hat{f}_r^{[\text{gain}]}, \hat{f}_r^{[\text{loss}]}\right\}$ against $\left\{f_r^{[\text{gain}]}, f_r^{[\text{loss}]}\right\}$

be captured with $f = \frac{p_{arm-misseg}}{p_{misseg}}$. If $f$ is low, then whole-chromosome missegregations are more likely to occur than chromosome-arm missegregations, therefore both arms of a chromosome tend to be gained or lost together. On the other hand, there is less correlation between chromosome arms when $f$ is high. Assuming $f$ is known, then $p_{misseg}$ and $\{\lambda_r\}$ compensate each other, as a given $f_r^{[gain]}$ can be approached by either increasing $p_{misseg}$ or increasing $\lambda_r$, and similarly for $f_r^{[loss]}$, therefore it is not necessary to fit both $p_{misseg}$ and $\{\lambda_r\}$. Therefore, in the following sections, we implement the variation (ii) mentioned above, which is equivalent to fitting $f$ and $\{\lambda_r\}$.

The prior distributions used for the following sections are:

- $p_{misseg} = 5 \times 10^{-5}$
- $p_{arm-misseg} \sim \text{Uniform}\left(10^{-5}, 10^{-4}\right)$
- $\lambda_r \sim \text{Uniform}\left(0.5, 1.5\right)$

and the reference data for `abcrf` comes from $N_{\text{simulations}} = 10{,}000$.

In order to not overfit, we limit the parameter inference to only chromosome arms with clear signals for selection, by only inferring the selection rates of chromosome arms $r$ with $\left|f_r^{[\text{gain}]} - f_r^{[\text{loss}]}\right| \geq 0.1$. For chromosome arms not satisfying this condition, we assume $\lambda_r = 1$, as the noise from neutral CNAs is indistinguishable from true selection signals.

#### 3.2 Application for TCGA pan-cancer data

The data in this study comes from [2], consisting of the gain and loss frequency as well as gene balance scores for each chromosome arm (excluding 13p, 14p, 15p, 21q, 22p, X and Y) from > 8,200 pan-cancer tumor-normal pairs in The Cancer Genome Atlas (TCGA). The gene balance score is computed based on the gene density on an arm. Charm(TSG,OG) considers the imbalance between Tumor Suppresor Genes and Oncogenes, and Charm(TSG,OG,Ess) additionally recognizes the Essential genes. Algorithm 1 is employed to find the selection rates for chromosome arms, given their gain/loss frequencies. We choose $\mathcal{S} = 100$ as using $\mathcal{S} > 8{,}200$ requires much more computing resources without improving the fitting quality appreciably. The parameters' posterior distributions are shown in **Supplementary**

**Fig. 18a**, and the comparison between simulations computed with parameters' posterior modes and the gain/loss data is presented in **Supplementary Fig. 18b**.

#### 3.3 Application for PCAWG cancer-specific data

`pcawg_specimen_histology_August2016_v9.xlsx` in folder `clinical_and_histology` at `https://dcc.icgc.org/releases/PCAWG/` contains the histology of cancer samples in PCAWG. We also retrieve the WGD status for each sample (`consensus_cnv/consensus.20170218.purity.ploidy.txt`). For the inference of each cancer type, we limit the data to include only non-WGD samples (`wgd_uncertaion=FALSE` and `wgd_status=no_wgd`) with high-quality CN data (at least one CN region with `star` $\geq 2$). The CN profile of each sample is downloaded from `consensus_cnv/consensus.20170119.somatic.cna.annotated.tar.gz`.

We analyze cancer types with at least 10 samples that satisfy these conditions. To find the gain/loss frequencies, we normalize each CN profile to diploid, and measure the frequencies of CN $> 2$ or $< 2$ for each chromosome arm.

For each cancer type, we use Algorithm 1 to infer the chromosome-arm selection rates. To evaluate the biological relevance of the inferred selection rates, we compare the mean selection rates and counts of selective chromosome arms with WGD proportions, as the inference was performed only on non-WGD samples. For each cancer type, we compute the mean selection rates for GAIN arms:

$$\widetilde{\lambda}^{[\text{gain}]} = \mathbb{E}\left\{\lambda_r : \lambda_r > 1\right\}$$

and for LOSS arms:

$$\widetilde{\lambda}^{[\text{loss}]} = \mathbb{E}\left\{\lambda_r : \lambda_r < 1\right\}$$

as well as the count of GAIN arms:

$$\widetilde{N}^{[\text{gain}]} = |\lambda_r : \lambda_r > 1|$$

and of LOSS arms:

$$\widetilde{N}^{[\text{gain}]} = |\lambda_r : \lambda_r < 1|$$

and the WGD proportions:

$$\%\text{WGD} = 100 \times \frac{\text{count of WGD samples}}{\text{count of all samples}}$$

**Supplementary Fig. 1-17** present for each cancer type, the parameters' posterior distributions, comparisons between simulations computed with parameters' posterior modes and PCAWG gain/loss data, and correlations between inferred chromosome arm selection rates and amplification/deletion frequencies.

### 4 Parameter studies of the chromosome arm selection model

We study the impacts of different components in the chromosome arm selection model on the statistics observable from the cancer samples.

The parameter sweep studies are designed as follow:

- The population starts at $t_0 = 0$ with $\mathcal{N}_0 = 1$ clone consisting of $N = N_0$ cells with diploid CN profile and no driver mutations.
- The selection model for chromosome arms (1.5.1) is applied. We do not consider driver gene mutations. The selection rates for chromosome arms are $\{\lambda_r\}$.
- Viability checkpoints: $\text{ploidy}_{\text{max}} = 6$, $\text{CN}_{\text{max}} = 8$, $\text{nullisomy}_{\text{max}} = 0$.
- Cells' turn-over rate is $\lambda = 1$ day.
- The total population is expected to follow $\{t_i, P_i\}$, i.e. the total cell count at time $t_i$ is expected to be $P_i$. Each simulation ends at $t_f = 300$ days, i.e. after approximately $(t_f - t_0)/\lambda = 300$ cell generations.
- 1000 cells at $t_{\text{sample}} = t_f$ are sampled.

For every parameter combination, we create 1,000 simulations and consider the following statistics:

- **Clone count:** for each simulation, we find distinct total CN profiles and their populations in the sample. Clone count is the count of different total CN profiles (ignoring convergent evolution).
- **Shannon diversity index:** computed from the count and sizes of the distinct total CN profiles using function `diversity` in package `vegan` (CRAN).

- **Clonal missegregation count:** equals count of missegregations that the most recent common ancestor (MRCA) in the sampled cell phylogeny had acquired. Clonal missegregations can be further divided into gains and losses of single chromosomes.

- **Subclonal missegregation count:** averaged over the count of missegregations dating from the MRCA to each sampled cell. Subclonal missegregations can be divided into gains and losses.

- **Age of MRCA:** assuming the MRCA last divided at $t_{\mathrm{MRCA}}$, the age of MRCA is computed as $a_{\mathrm{MRCA}} = -\frac{t_f - t_{\mathrm{MRCA}}}{t_f - t_0} \in (-1, 0)$. The MRCA is more recent, implying there is more selection, as $a_{\mathrm{MRCA}} \to 0$.

- **Distribution of ploidy:** a cell's ploidy is defined as the average bin-level CN. A sample's distribution of ploidy shows the percentages of its cells belonging to different ploidy after rounding. For instance, a normal cell with only a few gains or losses has ploidy 2. After WGD, its ploidy changes to 4.

#### 4.1 Probability of missegregation versus scale of selection rates

We seek to analyze the changes resulting from different probabilities of missegregation, representing chromosome-specific CNAs that impact the gene balance in a cell, and selection rates. The "base" for the selection rates $\{\bar{\lambda}_r\}$ comes from fitting the pan-cancer chromosome-arm gains and losses (Section 3.2).

This study is set up as follows:

- $N_0 = P_i = 1000$, therefore the cell population is expected to stay constant at 1000 cells throughout time. All remaining cells at $t_f$ are sampled.

- Whole Genome Duplication is not included: $p_{WGD} = 0$.

- The range for the probability of missegregation: $p_{misseg} = 10^{-3}, 2 \times 10^{-3}, \dots, 9 \times 10^{-3}, 10^{-2}$.

- The range for the scale of selection rates: $\gamma = 1, 2, \ldots, 9, 10$. For a given scale $\gamma$, the selection rates $\{\lambda_r\}$ used in the simulations are computed as:

$$\lambda_r = \begin{cases} 1 + \gamma(\bar{\lambda}_r - 1), \text{ if } \lambda_r > 1 \\ \frac{1}{1+\gamma\left(\frac{1}{\lambda_r} - 1\right)}, \text{ if } \lambda_r < 1 \\ 1, \text{ if } \lambda_r = 1 \end{cases} \tag{8}$$

which increases the selective strength of the gains and losses equally.

#### 4.2 Scale of selection rates for gains versus scale of selection rates for losses

- $N_0 = P_i = 1000$, therefore the cell population is expected to stay constant at 1000 cells throughout time. All remaining cells at $t_f$ are sampled.
- $p_{WGD} = 10^{-3}$.
- $p_{misseg} = 10^{-2}$.
- The scale for selection rates for gains $\gamma_g$ and for losses $\gamma_l$ range from $\gamma_g, \gamma_l = 1, 2, \ldots, 9, 10$. For each pair of $(\gamma_g, \gamma_l)$, the selection rates $\{\lambda_r\}$ used in the simulations are computed as:

$$\lambda_r = \begin{cases} 1 + \gamma_g(\bar{\lambda}_r - 1), \text{ if } \lambda_r > 1 \\ \frac{1}{1+\gamma_l\left(\frac{1}{\lambda_r} - 1\right)}, \text{ if } \lambda_r < 1 \\ 1, \text{ if } \lambda_r = 1 \end{cases} \tag{9}$$

which increases the selective strength of chromosome arm gains by a factor of $\gamma_g$ and losses by a factor of $\gamma_l$.

#### 4.3 Probability of Whole Genome Duplication versus probability of missegregation

- $N_0 = P_i = 1000$, therefore the cell population is expected to stay constant at 1000 cells throughout time. All remaining cells at $t_f$ are sampled.

- $p_{WGD} = 10^{-3}, 2 \times 10^{-3}, \ldots, 9 \times 10^{-3}, 10^{-2}$.
- $p_{misseg} = 10^{-3}, 2 \times 10^{-3}, \ldots, 9 \times 10^{-3}, 10^{-2}$.
- $\lambda_r = \bar{\lambda}_r$.

##### 4.4 Growth mode versus avegage cell count

- Growth mode $g = 1, 2, \ldots, 9, 10$.
- Average cell count $\bar{N} = 200, 400, \ldots, 1800, 2000$.
- For each growth mode $g$ and average cell count $\bar{N}$, the cell population dynamics $P(t)$ is calibrated to satisfy

$$\begin{cases} P(0) = 1000\frac{g}{10} \\ \int_0^{t_f} P(t) = \bar{N} \end{cases}$$

  assuming exponential growth, i.e. $P(t) = P(0) \cdot \exp(kt)$, the growth rate $k$ can then be found, and the population input for the simulator is $\{t_i, P_i\}$ where $P_i = P(t_i)$. Therefore, higher growth mode $g$ necessitates higher exponential growth rate of the total cell population.
- 1000 cells at $t_f$ are sampled.
- $p_{WGD} = 10^{-3}$.
- $p_{misseg} = 10^{-2}$.
- $\lambda_r = \bar{\lambda}_r$.

### 5 Inferring WGD parameters

#### 5.1 Parameter fitting scheme

We investigate the increased level of aneuploidy associated with Whole-Genome Duplication (WGD). We consider two distinct reasons for higher CNA counts in WGD sample observations. First, the WGD cells have twice as many chromosome copies, therefore there are more chances for chromosomes to be missegregated during cell divisions. To model this, we use a slightly different missegregation probability model than that in 1.3. Instead of fixing $p_{misseg}$ per cell division, we define $p_{misseg}^{homolog}$ as the probability that an individual chromosome homolog is missegregated during a division. In a given cell with CN profile $\{J_i\}$, where $J_i$ is the homolog count of chromosome $i$ (in a normal cell, $J_i = 2$ for $i = 1, \ldots, 22$, $J_i = 1$ for $i =$ X or Y), the relationship between the two probabilities is:

$$p_{misseg} = 1 - \left(1 - p_{misseg}^{homolog}\right)^{\sum_i J_i} \tag{10}$$

If $p_{misseg}^{homolog}$ is constant, the WGD cells would still missegregate at a higher rate, due to increased $J_i$.

The second reason for increased aneuploidy in WGD cells is due to their genomic instability. Let $p_{misseg}^{homolog;non-WGD}$ and $p_{misseg}^{homolog;WGD}$ be the chromosome homolog missegregation probabilities, defined as above but conditioned on whether the cell has undergone WGD. We define the WGD-aneuploidy rate as

$$\alpha = \frac{p_{misseg}^{homolog;WGD}}{p_{misseg}^{homolog;non-WGD}}$$

For $\alpha = 1$, the increased aneuploidy in WGD samples is solely as a result of higher genomic content for cells to duplicate and distribute during divisions. For $\alpha \gg 1$, heightened cellular failure to properly segregate is taken into account. The same formulation is performed for $p_{arm-misseg}$.

For each data cohort, the selection rates $\{\lambda_r\}$ and missegregation rates $p_{misseg}$ and $p_{arm-misseg}$ are taken from applying Algorithm 1 on non-WGD samples. We then convert the missegregation rates to $p_{misseg}^{homolog;non-WGD}$ and $p_{arm-misseg}^{homolog;non-WGD}$ using equation 10. The goal of parameter fitting is then limited to finding the WGD-aneuploidy rate $\alpha$ and $p_{WGD}$ (probability of WGD in each cell division).

The $\mathcal{S}$ samples in the data are grouped into $\mathcal{S}^-$ diploid samples and $\mathcal{S}^+$ samples with WGD. For each sample $k$ with CN profile $\left(\left\{J_i^{[k]}\right\}, \left\{K_{i,j}^{[k]}\right\}\right)$, we define the total CN vector as

$$\mathcal{T}^{[k]} = \left(\sum_{j=1,\ldots,J_1^{[k]}} K_{1,j}^{[k]}, \ldots, \sum_{j=1,\ldots,J_{\mathcal{N}}^{[k]}} K_{\mathcal{N},j}^{[k]}\right)$$

and then measure the fraction of genome altered (FGA) as the fraction of the genome with total CN different from 2:

$$\text{FGA}(k) = \frac{\left|t : \mathcal{T}^{[k]}(t) \neq 2\right|}{|\mathcal{T}^{[k]}|}$$

Finally, the data cohort is summarized by two statistics:

$$\text{WGD proportion} = \frac{|\mathcal{S}^{+}|}{[\mathcal{S}^{-}] + |\mathcal{S}^{+}|}$$

and

$$\text{WGD FGA difference} = \frac{\mathbb{E}\left(\text{FGA}(k) : k \in \mathcal{S}^{+}\right)}{\mathbb{E}\left(\text{FGA}(k) : k \in \mathcal{S}^{-}\right)}$$

which are used to infer $\alpha$ and $p_{WGD}$. Algorithm 2 outlines the fitting routine.

The prior distributions we used are:

- $\alpha \sim \text{Uniform}(0, 300)$
- $\log_{10}(p_{WGD}) \sim \text{Uniform}(-6.5, -3.5)$

and the reference data for `abcrf` comes from $N_{\text{simulation}} = 3{,}000$, where the statistics for each parameter set are computed from $\mathcal{S} = 100$ simulations.

##### 5.2 Application for PCAWG cancer-specific data

For the parameter inference for each cancer type, the PCAWG files were downloaded from `https://dcc.icgc.org/releases/PCAWG/` as in Section 3.3. Similarly, we removed cancer types with less than 10 non-WGD samples. Further, we removed cancer types with WGD proportion $\leq 10\%$, as cohorts with few WGD samples would lead to significant noise in the average FGA. **Supplementary Fig. 22-23** show the posterior distributions for $\alpha$ and $\log_{10}(p_{WGD})$ for each cancer type.

#### 6 Inferring driver gene parameters

##### 6.1 Parameter fitting scheme

Assume that a cancer type does not frequently have large CNAs (affecting chromosome arms or of larger extents) and are only associated with $N_{\text{drivers}}$

**Algorithm 2** Parameter fitting for Whole-Genome Duplication

**function** STATS($\alpha, p_{WGD}$)
  **for** $j \leftarrow 1, \ldots, \mathcal{S}$ **do**
    Create one simulation
    Find WGD status and compute FGA
  **end for**
  Compute WGD proportion and WGD FGA difference
**end function**

**for** $i \leftarrow 1, \ldots, N_{\text{simulations}}$ **do** ▷ Create reference data
  $\alpha^{[i]}, p_{WGD}^{[i]} \sim$ prior distributions
  $\left\{\text{WGD proportion}^{[i]}, \text{WGD FGA difference}^{[i]}\right\} \leftarrow \text{STATS}\left(\alpha^{[i]}, p_{WGD}^{[i]}\right)$
**end for**
**for** $p \leftarrow \alpha, p_{WGD}$ **do** ▷ ABC random forest
  Train `abcrf` on $\left\{p^{[i]}; \text{WGD proportion}^{[i]}, \text{WGD FGA difference}^{[i]}\right\}$
  Posterior distribution for $p \sim$ `abcrf` prediction for $\{\text{WGD proportion}, \text{WGD FGA difference}\}$
  $\hat{p}$ = mode of posterior distribution
**end for**
$\{\text{WGD proportion}^*, \text{WGD FGA difference}^*\} \sim \text{STATS}(\hat{\alpha}, \hat{p}_{WGD})$ ▷ Validation
Compare $\{\text{WGD proportion}^*, \text{WGD FGA difference}^*\}$ against $\{\text{WGD proportion}, \text{WGD FGA difference}\}$

driver genes, which are frequently mutated, or gained and lost via small focal CNAs. The cancer type can then be studied under the driver gene selection model (Section 1.5.2).

There are two steps in parameter inference. First, we model the length of the focal amplification and deletion events. We assume that the length $\mathcal{M}_{foc-amp}$ of an amplification event follows the Beta distribution with respect to the length $\mathcal{M}_{arm}$ of the chromosome arm that it occurs on:

$$\frac{\mathcal{M}_{foc-amp}}{\mathcal{M}_{arm}} \sim \text{Beta}(\alpha_{foc-amp}, \beta_{foc-amp})$$

(see Section 1.3). The shape parameters $\alpha_{foc-amp}$ and $\beta_{foc-amp}$ can then be inferred from all focal amplifications observed in the data cohort. Similarly, we can find the Beta shape parameters $\alpha_{foc-del}$ and $\beta_{foc-del}$ for the focal deletion events. We use function `fitdist` in the R library `fitdistrplus` to infer these parameters.

The second step consists of fitting the driver gene selection model. Assume that for each gene $r = 1, \ldots, N_{\text{drivers}}$, $f_r^{[mut]}$ is its mutation frequency in the data, and $f_r^{[gain]}$ and $f_r^{[loss]}$ are frequencies that it is focally amplified and deleted, respectively. We want to find the driver gene selection rates $\lambda_r$, and probabilities $p_{foc-amp}$ and $p_{foc-del}$ that a focal amplification or deletion occurs in a given cell division. Algorithm 3 outlines this step, consisting of two phases (building simulation library and inferring parameters with `abcrf`), similarly to 1.

#### 6.2 Application for CLLE-ES

CLLE-ES was excluded from our chromosome arm study (Section 3.3) because the PCAWG cohort did not exhibit any frequent CNAs affecting either whole chromosomes or chromosome arms, making the cancer type ideal for analysis with our driver gene selection model.

The list of focal CNAs is downloaded from `consensus_cnv/GISTIC_analysis/focal_input.rmcnv.pt_170207.seg.txt.gz` at `https://dcc.icgc.org/releases/PCAWG/`. We collect total CN change events associated with CLLE-ES by extracting CNAs associated with any sample in the cohort with "Seg.CN" $\neq 0$. $\{\alpha_{foc-amp}, \beta_{foc-amp}\}$ are then inferred by fitting the Beta distribution, as described above, to CNAs with "Seg.CN" $> 0$. Similarly, $\{\alpha_{foc-del}, \beta_{foc-del}\}$ are inferred by fitting the Beta distribution to CNAs with "Seg.CN" $> 0$.

**Algorithm 3** Parameter fitting for the driver gene selection model (Section 1.5.2)

**function** FREQUENCY$(p_{foc-amp}, p_{foc-del}, \{\lambda_r\})$
  **for** $j \leftarrow 1, \ldots, \mathcal{S}$ **do**
    Create one simulation
    Clonal CN profile ← CN profile of most populous clone
  **end for**
  Compute $\left\{f_r^{[\text{mut;i}]}, f_r^{[\text{gain;i}]}, f_r^{[\text{loss;i}]}\right\}$ from the clonal CN profiles
**end function**

**for** $i \leftarrow 1, \ldots, N_{\text{simulations}}$ **do** ▷ Create reference data
  $p^{[i]}_{foc-amp}, p^{[i]}_{foc-del}, \left\{\lambda_r^{[i]}\right\} \sim$ prior distributions
  $\left\{f_r^{[\text{mut;i}]}, f_r^{[\text{gain;i}]}, f_r^{[\text{loss;i}]}\right\} \leftarrow \text{FREQUENCY}\left(p^{[i]}_{foc-amp}, p^{[i]}_{foc-del}, \left\{\lambda_r^{[i]}\right\}\right)$
**end for**
**for** $p \leftarrow p_{foc-amp}, p_{foc-del}, r = 1, \ldots, N_{\text{drivers}}$ **do** ▷ ABC random forest
  Train `abcrf` on $\left\{p^{[i]}; f_r^{[\text{mut;i}]}, f_r^{[\text{gain;i}]}, f_r^{[\text{loss;i}]}\right\}$
  Posterior distribution for $p \sim$ `abcrf` prediction for $\left\{f_r^{[\text{mut}]}, f_r^{[\text{gain}]}, f_r^{[\text{loss}]}\right\}$
  $\hat{p}$ = mode of posterior distribution
**end for**
$\left\{\hat{f}_r^{[\text{mut}]}, \hat{f}_r^{[\text{gain}]}, \hat{f}_r^{[\text{loss}]}\right\} \sim \text{FREQUENCY}\left(\hat{p}_{foc-amp}, \hat{p}_{foc-del}, \left\{\hat{\lambda}_r\right\}\right)$ ▷ Validation
Compare $\left\{\hat{f}_r^{[\text{mut}]}, \hat{f}_r^{[\text{gain}]}, \hat{f}_r^{[\text{loss}]}\right\}$ against $\left\{f_r^{[\text{mut}]}, f_r^{[\text{gain}]}, f_r^{[\text{loss}]}\right\}$

The list of driver events comes from `TableS3_panorama_driver_mutations_ICGC_samples.controlled.tsv` in folder `driver_mutations` at `https://dcc.icgc.org/releases/PCAWG/`. For each gene associated with at least one sample in the CLLE-ES cohort, we get the genomic location from Cancer Gene Census (CGC, `https://cancer.sanger.ac.uk/census#cl_sub_tables`). Our selection model requires that each driver gene must be classified as either Tumor Suppressor Gene (TSG) or oncogene, under the assumption that TSGs are associated with CN losses and oncogenes are associated with CN gains (Section 1.5.2). For each gene $r$, we follow these steps to assign a role:

1. Find frequencies of gain $f_r^{[gain]}$ and loss $f_r^{[loss]}$ in the CLLE-ES cohort.
2. If $f_r^{[gain]} > f_r^{[loss]}$, assign the gene as oncogene.
   If $f_r^{[gain]} < f_r^{[loss]}$, assign the gene as TSG.
3. If $f_r^{[gain]} = f_r^{[loss]}$, the gene role is assigned according to CGC.
4. If the CGC classification is both TSG and oncogene or neither, then assign gene as TSG if $f_r^{[gain]} = 0$.

Because we limit to autosomes, we remove driver genes that are either not listed in CGC or located on chromosomes X and Y. No driver gene in our list was affected by focal amplifications, therefore we remove $p_{foc-amp}$ when applying Algorithm 3. **Supplementary Fig. 24** shows the posterior distributions of each driver gene's selection rate.
